# KLHL41 orchestrates sarcomere assembly and size to drive skeletal muscle hypertrophy *in vivo*

**DOI:** 10.1101/2025.05.30.655367

**Authors:** Hyewon Han, Janet Shi, Alfredo H Bongiorno, Arian Mansur, Jeffrey Widrick, Grace Tate, Sehajpreet Gill, Esmat Karimi, Henk Granzier, Jeffrey R Moore, Vandana A Gupta

## Abstract

Sarcomere assembly and growth are fundamental processes essential for the development, function, and repair of skeletal muscle. However, the mechanisms underlying sarcomere formation and assembly *in vivo*, which are critical for the formation of functional myofibers in vertebrates, remain poorly understood. Defects in sarcomeres contribute to muscle dysfunction in numerous genetic and acquired myopathies, yet the lack of a clear understanding of specific sarcomeric defects has hindered the development of effective therapies. Nemaline myopathy (NM) is caused by mutations in genes that primarily affect sarcomere structure and function. The disease is clinically heterogeneous, with the congenital form being the most severe. Using zebrafish models of congenital forms of NM, we investigated sarcomere assembly in *vivo* during skeletal muscle development to identify key steps contributing to myofibril formation and muscle growth under both normal and disease conditions. Our findings demonstrate that the gene encoding the sarcomeric protein KLHL41 plays a critical role in the formation and directional organization of new sarcomeres within developing myofibrils, facilitating both radial and longitudinal muscle growth. Dysregulation of sarcomeric proteins and impaired protein turnover in the KLHL41-NM zebrafish model resulted in the development of sarcomeric defects. Moreover, we show that the interaction between the KLHL41-troponin complex is essential for muscle growth and cross-bridge regulation in developing muscle. These studies address a significant gap in the understanding of sarcomeric defects in myopathies and will help guide the development of targeted therapies aimed at rescuing these processes.

## INTRODUCTION

The sarcomere is the fundamental contractile unit of striated muscle, playing a central role in muscle structure and function [1, 2]. Within the sarcomere, thick filaments composed of myosin interact with actin filaments, generating the cross-bridge cycles that drive muscle contraction. This intricate structure ensures the precise alignment and coordination necessary for efficient force generation. Sarcomeres are arranged end-to-end to form long myofibrils, which bundle together to create myofibers. Despite their well-defined structural organization, the precise mechanisms governing their formation and assembly into myofibrils remain incompletely understood.

Sarcomere assembly and growth involve a complex interplay of structural proteins, mechanical forces, signaling, metabolic pathways, epigenetic processes, and molecular chaperones that coordinate the sequential organization of actin and myosin filaments into repeating sarcomeric units [3–12]. Several models, including the pre-myofibril and stress fiber models, propose different mechanisms for myofibril assembly [13–16].

However, the exact sequence of events and regulatory factors governing vertebrate muscle formation *in vivo* remains unclear. A deeper understanding of sarcomere formation could provide critical insights into muscle development, regeneration, and the molecular basis of human myopathies.

Defects in sarcomere structure and function play a pivotal role in the development of genetic and acquired myopathies, which are characterized by muscle weakness and dysfunction [17, 18]. Mutations in genes encoding key sarcomeric proteins—such as titin, nebulin, myosin, actin, and tropomyosin—can disrupt the structural integrity and contractile function of muscle fibers, leading to hypotrophy, atrophy, and impaired force transmission [19–22]. Nemaline myopathy (NM) is a congenital neuromuscular condition marked by muscle weakness, hypotonia, and the presence of rod-like nemaline bodies in muscle fibers [23]. NM arises from mutations in genes encoding essential sarcomeric and cytoskeletal proteins, which destabilize thin filaments and compromise force generation and muscle contraction [24–34]. While the severity of NM ranges from mild cases with near-normal lifespan to severe neonatal forms resulting in respiratory failure, there is currently no cure. Elucidating the molecular mechanisms underlying these defects could pave the way for targeted therapies aimed at restoring sarcomere integrity and improving patient outcomes.

To investigate *in vivo* sarcomere assembly, growth, and the pathogenic mechanisms of NM, we developed zebrafish models of NM. Zebrafish models of skeletal muscle disorders closely recapitulate the pathological features observed in human patients, making them valuable tools for uncovering the molecular and cellular processes driving disease progression. Our studies reveal that KLHL41 plays a crucial role in skeletal muscle hypertrophy by regulating sarcomere size—and when defective, it results in thinner myofibrils with reduced force generation. We further demonstrate that KLHL41 is a key regulator of sarcomere assembly and growth, as its absence significantly reduces the kinetics of sarcomere formation. Most importantly, our findings establish that congenital nemaline myopathy arises primarily from defects in sarcomere assembly rather than sarcomere maintenance, challenging previous assumptions and offering a more precise understanding of disease pathology. This refined perspective provides a foundation for developing more effective therapeutic strategies aimed at restoring sarcomere function and improving clinical outcomes for patients with NM.

## MATERIALS AND METHODS

### Zebrafish Maintenance and Husbandry

Zebrafish were maintained and bred using standard methods as described [35] All experiments and procedures were approved by the Institutional Animal Care and Use Committee at Brigham and Women’s Hospital. Wild-type fish were obtained from Tubingen (TU) line and staged by hours (h) or days (d) post-fertilization at 28.5°C. Zebrafish embryonic, larval, juvenile, and adult stages of development have been described previously [35].

### Creation of Zebrafish Lines

sgRNAs were designed using the web-based ZiFiT Targeter program (http://zifit.partners.org/) and targeting distinct sites in exon 1 or exon 2 of zebrafish Kelch genes. The first 2 bp GG sequences at the 5’ end of the target site are a constraint 20 imposed by T7 promoter sequence requirements in addition to the NGG protospacer adjacent motif (PAM) sequence requirement immediately 3’ to the target site. Two oligonucleotides, each of 22 nucleotides in length, were used to construct the guide RNA for each target site. Forward and reverse primers were annealed to create a sgRNA oligonucleotide duplex. Primer sequences are summarized in Table S2. The zebrafish guide RNA expression vector pDR274 was used to create the sgRNAs expression system using the T7 promoter followed by In Vitro Transcription as previously described [36]. sgRNAs and Cas9 protein (Thermo Scientific, CA) were co-injected into the yolk sac of one- and two-celled stage zebrafish embryos. Each embryo was injected with a 5ul injection solution containing 2ul of 400ng/nl Cas9 protein and 3ul of 100ng/ul sgRNA. Injected embryos were inspected under the microscope for 3 days and were classified as dead, deformed, or normal phenotypes. Embryos displaying normal phenotypes were analyzed to test the efficacy of sgRNAs by identifying target site mutations. To analyze injected fish, genomic DNA was extracted from 6-8 pooled embryos at 3-5 days post fertilization (dpf) and used for DNA sequencing experiments by topo cloning.

### Identification of Founder Fish and Generation of Isogenic Stable Mutant Fish

Because a fertilized zebrafish embryo develops quickly, direct delivery of sgRNA-Cas9 protein via injection results in chimeric embryos. Founder fish were determined by genotyping tail fin clips of the F0 generation and observing mosaicism at the target site. The mosaic F0 generation was outcrossed to wild-type TU fish for at least three generations for studies presented in this work. The sequences of sgRNAs are listed in Table S1, and sequencing primers are listed in Table S2.

### Kinematic Measurements in Zebrafish Larvae

The methods used to analyze escape response kinematics have been previously described in detail [37]. Briefly, a single larva was placed in a small arena, and an escape response was evoked using a brief electrical field pulse. The response was captured at 1000 frames/s by a high-speed camera. Recording began 10 ms prior to the stimulus and concluded after we had sufficient data to evaluate the initial 60 ms of escape response movement. A deep-learning neural network developed using the open-source toolkit DeepLabCut identified seven anatomical key points along the body and tail on each video frame [38, 39]. The x- and y-coordinates of these key points were used to calculate the linear kinematics of the larvae’s putative center of mass (COM, defined by a key point at the posterior edge of the swim bladder) and the angular kinematics of the propulsive tail. The average of three separate escape responses obtained per larva was used for analysis.

### Sample Preparation for Mass Spectrometry

Samples for protein analysis were prepared essentially as previously described [40, 41]. Proteomes were extracted using a buffer containing 200 mM EPPS pH 8.5, 8M urea, 0.1% SDS, and protease inhibitors. Following lysis, 50 µg of each proteome was reduced with 5 mM TCEP. Cysteine residues were alkylated using 10 mM iodoacetamide for 20 minutes at RT in the dark. Excess iodoacetamide was quenched with 10 mM DTT. A buffer exchange was carried out using a modified SP3 protocol [42]. Briefly, ∼250 µg of Cytiva SpeedBead Magnetic Carboxylate Modified Particles (65152105050250 and 4515210505250), mixed at a 1:1 ratio, were added to each sample. 100% ethanol was added to each sample to achieve a final ethanol concentration of at least 50%. Samples were incubated with gentle shaking for 15 minutes. Samples were washed three times with 80% ethanol. Protein was eluted from SP3 beads using 200 mM EPPS pH 8.5 containing Lys-C (Wako, 129-02541). Samples were digested overnight at room temperature with vigorous shaking. The next morning, trypsin was added to each sample and further incubated for 6 hours at 37° C. Acetonitrile was added to each sample to achieve a final concentration of ∼33%. Each sample was labeled, in the presence of SP3 beads, with ∼125 µg of TMT reagents (ThermoFisher Scientific). Following confirmation of satisfactory labeling (>97%), excess TMT was quenched by the addition of hydroxylamine to a final concentration of 0.3%. The full volume from each sample was pooled, and acetonitrile was removed by vacuum centrifugation for 1 hour. The pooled sample was acidified, and peptides were de-salted using a Sep-Pak 50mg tC18 cartridge (Waters). Peptides were eluted in 70% acetonitrile 1% formic acid and dried by vacuum centrifugation.

### Basic pH Reversed-Phase Separation (BPRP)

TMT labeled peptides were solubilized in 5% acetonitrile/10 mM ammonium bicarbonate, pH 8.0, and ∼300 µg of TMT labeled peptides were separated by an Agilent 300 Extend C18 column (3.5 μm particles, 4.6 mm ID and 250 mm in length). An Agilent 1260 binary pump coupled with a photodiode array (PDA) detector (Thermo Scientific) was used to separate the peptides. A 45 minute linear gradient from 10% to 40% acetonitrile in 10 mM ammonium bicarbonate pH 8.0 (flow rate of 0.6 mL/min) separated the peptide mixtures into a total of 96 fractions (36 seconds). A total of 96 Fractions were consolidated into 24 samples in a checkerboard fashion and vacuum-dried to completion. Each sample was desalted via Stage Tips and re-dissolved in 5% formic acid/ 5% acetonitrile for LC-MS3 analysis.

### Liquid chromatography Separation and Tandem Mass Spectrometry (LC-MS3)

Proteome data were collected on an Orbitrap Eclipse mass spectrometer (ThermoFisher Scientific) coupled to a Proxeon EASY-nLC 1000 LC pump (ThermoFisher Scientific). Fractionated peptides were separated using a 120 min gradient at 550 nL/min on a 35 cm column (i.d. 100 μm, Accucore, 2.6 μm, 150 Å) packed in-house. MS1 data were collected in the Orbitrap (60,000 resolution; maximum injection time 50 ms; AGC 4 × 10^5^). Charge states between 2 and 5 were required for MS2 analysis in the ion trap, and a 120 second dynamic exclusion window was used.

Top 10 MS2 scans were performed in the ion trap with CID fragmentation (isolation window 0.5 Da; Turbo; NCE 35%; maximum injection time 35 ms; AGC 1 × 10^4^). Real-time search was used to trigger MS3 scans for quantification. [PMID: 32126768] MS3 scans were collected in the Orbitrap using a resolution of 50,000, NCE of 55%, maximum injection time of 250 ms, and AGC of 1.25 × 10^5^. The close out was set at two peptides per protein per fraction [43]

### Data Analysis for Mass Spectrometry

Raw files were converted to mzXML, and monoisotopic peaks were re-assigned using Monocle [44]. Searches were performed using the Comet search algorithm against a zebrafish (*Danio rerio*) database downloaded from Uniprot in February 2023. We used a 50 ppm precursor ion tolerance, 1.0005 fragment ion tolerance, and 0.4 fragment bin offset for MS2 scans collected in the ion trap. TMT on lysine residues and peptide N-termini (+229.1629 Da) and carbamidomethylation of cysteine residues (+57.0215 Da) were set as static modifications, while oxidation of methionine residues (+15.9949 Da) was set as a variable modification.

Each run was filtered separately to 1% False Discovery Rate (FDR) on the peptide-spectrum match (PSM) level. Then, the proteins were filtered to the target 1% FDR level across the entire combined data set. For reporter ion quantification, a 0.003 Da window around the theoretical m/z of each reporter ion was scanned, and the most intense m/z was used. Reporter ion intensities were adjusted to correct for isotopic impurities of the different TMT reagents according to manufacturer specifications.

Peptides were filtered to include only those with a summed signal-to-noise (SN) ≥ 60 across all TMT channels. For each protein, the filtered peptide TMT SN values were summed to generate protein quantification values. The signal-to-noise (S/N) measurements of peptides assigned to each protein were summed (for a given protein).

These values were normalized so that the sum of the signal for all proteins in each channel was equivalent, thereby accounting for equal protein loading.

### Native Thin filament Preparation

Buffers were prepared for native thin filament (NTF) isolation as described [45]. The relaxing solution consisted of 50 mM NaCl, 5 mM MgCl₂, 2 mM EGTA, 1 mM dithiothreitol (DTT), 2.5 mM ATP, and 2.5 mg/mL phosphocreatine disodium salt hydrate, with the pH adjusted to 7.0 using phosphate buffer. The homogenization buffer contained 25 mM imidazole, 1 mM EGTA, 100 mM KCl, 5 mM MgCl₂, 5 mM ATP, 10 mM DTT, and 0.1% Triton X-100, adjusted to a pH of 6.45. The suspension buffer consisted of 25 mM imidazole, 1 mM EGTA, 100 mM KCl, 5 mM MgCl₂, and 5 mM ATP, adjusted to pH 7.9 with 0.55 mM Tris. The storage buffer consisted of a homogenization buffer without ATP.

Euthanized zebrafish larvae (5 dpf) were centrifuged at 10,000g to remove excess water. Zebrafish (<10 mg) were dissected in 500 µL of relaxing solution and teased apart with Dumont tweezers over the course of an hour to isolate single myofibers. Further dissection was performed in 500 µL of relaxing solution containing 0.5% Triton X-100. Subsequently, homogenization was performed in 150 µL of homogenization buffer using a 0.2 mL glass homogenizer. Samples were centrifuged for 30 minutes at 40,000g, and the pellet was discarded. The supernatant was centrifuged for 45 minutes at 200,000g to isolate NTFs. The resulting pellet was resuspended in 75–100 µL of suspension buffer and centrifuged for 30 minutes at 40,000g to remove residual debris. The final supernatant was centrifuged again at 200,000g for 45 minutes to obtain the purified NTF pellet. The NTF pellet was solubilized overnight in 40–60 µL of storage buffer.

Protein concentration was determined using the Bradford assay and labeled with TRITC-phalloidin (tetramethyl rhodamine) 1uM by overnight incubation at 4°C. All centrifugation steps were performed at 4°C. Dissections were conducted either on ice or in a 4°C environmental room. A TLA-100 rotor was used for centrifugation at 40,000g (32,198 rpm), while centrifugation at 200,000g was performed at 71,996 rpm.

### *In vitro* Motility Assay

The in-vitro motility assays were performed as previously described [46], with modifications to accommodate NTFs. The experiments were conducted in triplicate using the same dual-well flow cell. Flow cells were assembled by fastening a nitrocellulose-coated coverslip to a glass slide with double-sided tape.

Chicken skeletal muscle myosin was purified as previously described and diluted to 50 μg/mL in high-salt buffer (300 mM KCl, 25 mM BES, 5 mM EGTA, 4 mM MgCl₂, and 10 mM DTT, pH 7.4) and added to the flow cell [47]. Unbound or weakly bound myosin was removed by washing the flow cells with actin buffer (55 mM KCl, 25 mM BES, 5mM EGTA, 4 mM MgCl2, 10 mM DTT, pH 7.4), followed by the addition of 1 mg/mL bovine serum albumin (BSA) diluted in high-salt buffer to prevent nonspecific binding of thin filament proteins to the nitrocellulose membrane.

Unlabeled NTFs sourced from control or *klhl41b* zebrafish larvae (1 μM) were diluted in low-salt buffer (55 mM KCl, 25 mM BES, 5 mM EGTA, 4 mM MgCl₂, and 10 mM DTT, pH 7.4) and added into the myosin-coated flow cell and incubated for 2 minutes. A wash step was performed with 1 mM ATP to block any remaining non-functional myosin, followed by four additional low-salt buffer washes to remove excess ATP. Finally, regulated movement of labeled NTFs was initiated by adding motility buffer (low-salt buffer with 10 mM DTT, dextrose, and 0.5% methylcellulose) containing 1 mM ATP and calcium buffers at varying concentrations (pCa 4.0 to pCa 10.0). A scavenger mix containing glucose oxidase and catalase was also included to prevent photobleaching. The sliding velocity of the labeled NTFs and the percentage of motile filaments were analyzed using the ImageJ plug-in MTrackJ (https://imagescience.org/meijering/software/mtrackj/).

### Sample Preparation and Western Blotting Analysis

C2C12 Cells (10-15mg) were placed in RIPA Lysis and Extraction Buffer (Thermo Fisher Scientific) with a cocktail of protease inhibitors and homogenized (2 X 15 seconds) using the Tissuemiser homogenizer (Thermofisher Scientific). Samples were separated on an SDS-PAGE and blotted onto polyvinylidene difluoride (PVDF) membranes. The membranes were blocked using 5% non-fat milk powder in 1X Tris Buffered Saline (Boston Bioproducts, MA) and 0.1% TWEEN® 20 (Sigma Aldrich, cat. no. P9416) (TBST) for 1 hour at room temperature and incubated with primary antibodies overnight at 4°C. The membranes were subsequently washed and incubated with polyclonal anti-mouse-IgG antibody conjugated to horseradish peroxidase. The antibody pairings and associated dilutions are: Anti-FLAGM2 1: 250 (F1804, Sigma-Aldrich); TNNT3 (Sigma Aldrich, AV51287), TNNI2 (Thermofisher, TA807853), TNNC2 (Millipore Sigma, HPA043174), GAPDH (Cell Signaling Technology, 2118S). Secondary antibodies were anti-rabbit 1:1000 (170-6515, Biorad) and anti-mouse, 1:1000 (170-6516, Biorad). The quantification of protein bands was performed using Image J.

### Whole mount Immunofluorescence

Whole mount immunofluorescence on zebrafish was performed as previously described [48]. 1:100 Laminin (Millipore MAB1922) and 1:40 FITC-phalloidin (Thermofisher Scientific) were used. The secondary antibody was anti-mouse,1:250 (A11005, Thermofisher Scientific).

### C2C12 Cell Culture Studies

Coimmunoprecipitation of KLHL41 and troponins was performed using the previously described method [9]. To study the reciprocal interaction between KLHL41 and troponins, C2C12 cells (ATCC, #CRL-1772) were transfected with different amounts of KLHL40-pEZYFLAG plasmid. Cells were differentiated for two days and cell lysates were prepared in RIPA buffer, and proteins were analyzed by Western blot analysis.

### Electron Microscopy

Muscle tissue was dissected from juvenile and adult zebrafish, deskinned, and fixed in formaldehyde–glutaraldehyde– picric acid in cacodylate buffer overnight at 4°C, followed by osmication and uranyl acetate staining. Subsequently, tissue samples were dehydrated in a series of ethanol washes and finally embedded in Taab epon (Marivac Ltd., Nova Scotia, Canada). Ninety-five nanometer sections were cut with a Leica ultra cut microtome, picked up on 100 m formvar-coated Cu grids, and stained with 0.2% lead citrate. Sections were viewed and imaged by Joel 1200EX Transmission Electron Microscope (Electron Microscopy Core, Harvard Medical School).

### qRT-PCR

Total RNA was extracted from the control or mutant zebrafish embryos (2 dpf) using RNeasy mini kit (Qiagen) and cDNA was synthesized using Superscript 1V Reverse Transcriptase kit using random hexamers. RT-PCR was performed as previously described [48]

## Data analysis

The statistical significance and *p* values (or non-significant, n.s.) between groups were calculated using GraphPad Prism 9 using t-test or one-way ANOVA and reported in the figures.

## Materials Availability

Newly created zebrafish lines and plasmids generated in this work are available on request.

## Data Availability

Mass Spectrometry Data has been submitted to ProteomeXchange (Project Accession: PXD061858) These datasets will be made public upon acceptance of the manuscript.

Alternatively, reviewers can access the dataset by logging in to the PRIDE website (http://www.ebi.ac.uk/pride) using the following account details:

**Username:**, **Password:** m06c3U4ayBoR

## RESULTS

### KLHL41 deficiency results in skeletal muscle dysfunction in zebrafish

Mutations in *KLHL41* result in severe muscle weakness at birth, underscoring its essential role in normal skeletal muscle development [32]. To uncover the pivotal role of KLHL41 in normal and disease states in skeletal muscle, we focused on identifying the specific defects during myogenesis that are regulated by KLHL41. The differentiation of myogenic precursors to myofibers during myogenesis is evolutionarily conserved in vertebrates [49]. Therefore, we created a series of loss-of-function *klhl41* alleles for *KLHL41* zebrafish orthologs using CRISPR/Cas9 gene editing that allows the early *in vivo* analysis of the skeletal muscle development (Supplemental Table 1, Figure 1A-B). The *KLHL41* ortholog in zebrafish is duplicated as *klhl41a* and *klhl41b*. While Klhl41a deficiency did not affect survival, Klhl41b and Klhl41ab deficiency resulted in reduced survival and death of mutants by 7 dpf. Phenotypic analysis of the control and mutant groups showed no obvious differences by light microscopy (Figure 1C, left panel).

**Figure 1.**
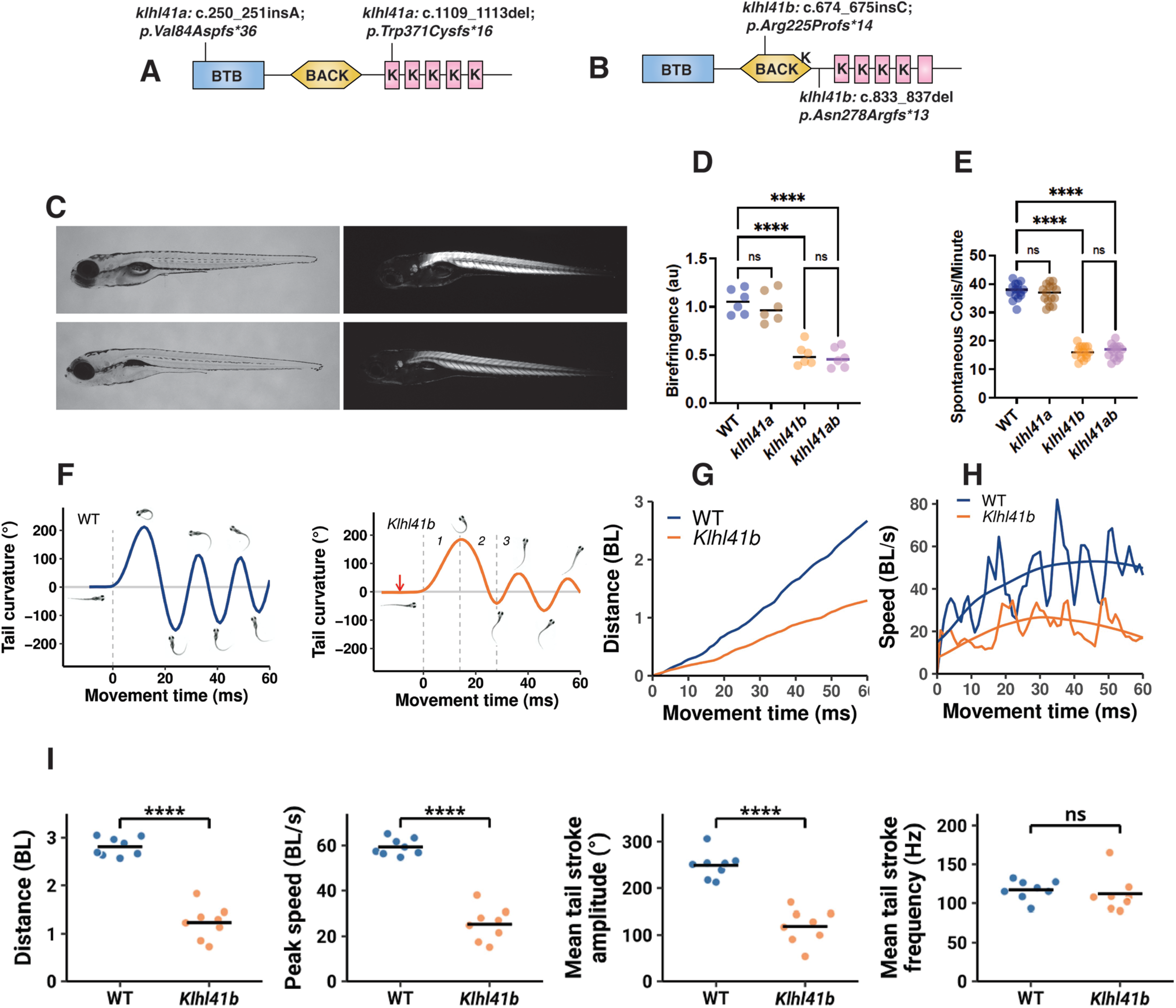
*klhl41b* is required for early skeletal muscle development by initiating myofiber assembly. (A-B) Zebrafish *klhl41* series of knockout mutant alleles were generated by the CRISPR-Cas9 approach and analyzed after outcrossing for 4-5 generations. The *klhl41* gene is duplicated in zebrafish, and knockout mutant alleles were created for both duplicated genes (*klhl41a* and *klhl41b*). (C-D) Phenotypic (left panel) and birefringence analysis (right panel) of control, *klhl41a*, *klhl41b,* and *klhl41ab* larval fish in normal (left panel) and polarized light (right panel) demonstrating normal birefringence in *klhl41a* mutants but reduced brightness in the *klhl41b* and *klhl41ab* mutants compared to controls (5 dpf, n=6 in each group from three different clutches). (E) Quantification of the spontaneous coiling at 18 hpf demonstrating early muscle weakness in *klhl41b* and *klhl41ab* embryos (n=20-24 in each group from three different clutches). (F) Tail curvature of a wild-type and a *klhl41b* mutant larva during an escape response. The escape response stimulus is represented by the red arrow. The vertical dashed lines delineate the onset of movement (movement time = 0) and the three subsequent escape response stages: the C-start (1), the power stroke (2), and burst swimming (3). The images show the larvae prior to movement, at the stage 1 to 2 transition, the stage 2 to 3 transition, and during each subsequent stroke of stage 3 burst swimming. (G) Distance traveled by the larval vs. movement time for the wild-type and mutant larvae shown in (E). (H) The instantaneous speed of the wild-type and mutant larvae shown in (E). Speed oscillates due to the undulatory nature of larval swimming. The speed response was smoothed for further analysis. (I) Cumulative distance covered in 60 ms of escape response swimming, peak instantaneous speed, and the mean tail stroke amplitude and frequency of stage 3 burst swimming for wild-type and *klhl1b* larvae. Data are mean ± S.E.M; unpaired t-test, parametric for two groups or one-way analysis of variance (ANOVA) and Tukey’s HSD test for multiple groups comparison. **** p<0.0001; ns: not significant (D-E) . Each point represents a single larva (the mean of three separate trials) with a horizontal bar indicating group means. **** p<0.0001; ns: not significant (F-I).

Examination of skeletal muscle organization by polarized light showed no significant difference in birefringence between the control and *klhl41a* mutants (Figure 1D).

However, there was reduced birefringence in the myotome of *klhl41b* and *klhl41ab* mutants compared to controls, indicating disrupted sarcomere organization (5 dpf).

Primary myogenesis leads to the formation of primary axial musculature during embryonic development in zebrafish that can contract to produce spontaneous coiling in the muscle as the first measure of motor function between 20-28 hours post fertilization (hpf). The frequency of spontaneous body coiling was reduced in *klhl41b* and *klhl41ab* embryos compared with the control siblings at 20 hpf, suggesting a defect in primary skeletal muscle contractile machinery (Figure 1E). As no significant differences were seen between *klhl41b* and *klhl41ab* double mutants, all following studies were performed in *klhl41b* mutants. One of the duplicated genes for human orthologs is often redundant in zebrafish [50], the normal structure and function of skeletal muscle in *klhl41a* mutants and similar phenotypes in *klhl41b and klhl41ab* suggest that zebrafish *klhl41a* gene is redundant in zebrafish while *klhl41b* is an ortholog of the human *KLHL41* gene and, similar to patients, mutations in the *klhl41b* gene result in an early onset myopathy.

To evaluate the effect of Klhl41b deficiency on the motor function, escape swimming response of the control and *klhl41b* mutants was studied at 6 and 7 dpf using high speed videography. [37]. Escape responses were partitioned into an initiating C-start(stage 1) where the larva curves to bring the head and tail together, a counter bend or power stroke (stage 2), and a concluding burst of fast undulatory tail strokes (stage 3)(Figure 1F). *klhl41b* mutants attained a significantly slower peak instantaneous speed and covered significantly less overall distance than wild-type larvae (Figure 1G-I). Of particular note was the inability of *klhl41b* larvae to bend to the same extreme angles as wild-type larvae during their C-start (Figure S1 - Stage 1 Final tail curvature), their power stroke (Figure S1 - Stage 2 Tail stroke amplitude), and during stage 3 burst swimming (Figure 1-I).

To compensate for these truncated tail strokes, larvae could perform strokes at a faster angular velocity. However, their tail stroke frequency during stage 3 was similar to wild-type (Figure 1-F) and the maximal tail stroke angular velocity was reduced in all three stages. Therefore, the impaired locomotion of the *klhl41b* mutants reflects their inability to bend the propulsive tail to the same extreme angles observed for wild-type controls coupled with a failure to increase tail stroke angular velocity and stroke frequency to compensate. These results show that deficiency of Klhl41b in zebrafish leads to reduced survival, changes in skeletal muscle organization, and reduced muscle function.

### KLHL41 is required for the initiation of myofibril assembly but dispensable for myoblast fusion

KLHL41 deficiency results in hypotonia and low muscle tone in affected patients. *KLHL41* is expressed in developing mouse (E9.5) and zebrafish somites (E9.5), suggesting a requirement in vertebrate skeletal muscle during early myogenesis [32, 51]. To identify the initial events regulated by KLHL41 during skeletal muscle development, we analyzed the myotome of control and *klhl41b* mutants at 16-18 hpf for any fusion defects as myoblasts are fused and multinucleated myotubes are attached to the myotendinous junction (Figure S2). No significant differences were identified in the myonuclei number of control and *klhl41b* mutant myotubes (16 hpf), suggesting that KLHL41 is not required for primary myoblast fusion *in vivo* and the defects observed in skeletal muscle potentially originated during sarcomere formation.

Sarcomere assemblies begin as myotubes are attached to myotendinous junctions in developing somites. Several models have been proposed to explain the periodic organization of sarcomeres into myofibrils (myofibrillogenesis). From *in vitro* studies or invertebrate models, they can be broadly categorized into 1) *de novo* sarcomere formation and their subsequent assembly into myofibrils and 2) immature actin filaments organizing into premyofibrils and progressively maturation into functional myofibrils [5, 15, 52]. Therefore, to identify specific stages of myofibrillogenesis regulated by KLHL41, we analyzed the developing sarcomere assemblies in control and *klhl41b* mutants by electron microscopy (Figure 2). Ultrastructural examination in control embryos showed the earliest presence of sarcomeres in developing somites around 16 hpf, with parallel rows of myofibrils emerging from the myotendinous junctions at the periphery of the myofibers (Figure 2A). No evidence of small, isolated actin-myosin assemblies or immature actin filaments was seen in the control embryos, suggesting the formation of rows of myofibrils is the first step towards sarcomere formation in normal skeletal muscle. Compared to single rows of myofibrils in control myotubes, a significant number of the myotubes (∼28%) in the *klhl41b* mutant embryos showed the absence of myofibrils at this stage (Figure 2B-C) or the presence of much thinner myofibrils compared to controls (Figure 2D-E) as indicated by the sarcomere height (Figure 2F).

**Figure 2.**
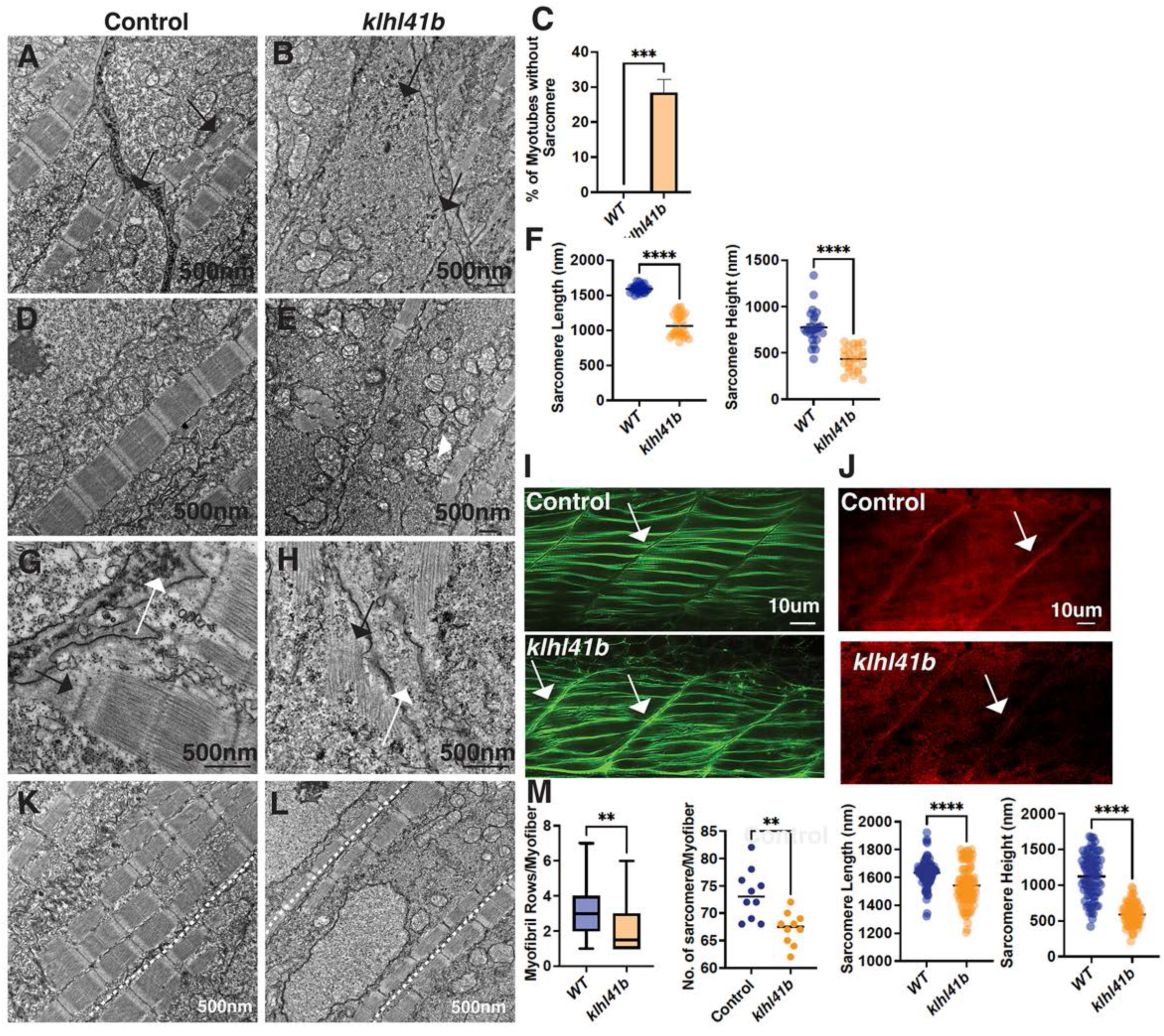
KLHL41 regulates the initiation of myofibril formation and sarcomere size during primary myogenesis. (A-B) Transmission electron microscopy (16 hpf) revealed the presence of single rows of myofibrils in differentiating myotubes in the control myotome. A significant number of myotubes in the *klhl41b* mutants failed to show the presence of myofibrils (quantified in C). (D-E) Compared to the control, myotubes containing myofibrils in *klhl41b* mutants contained smaller sarcomeres with reduced length as well as height (Quantified in F). (G-H) Myotubes lacking myofibrils exhibited electron-dense protein aggregates near myotendinous junctions (arrows) or single sarcomeres originating from the myotendinous junctions (arrow) in *klhl41b* mutants and were not seen in controls. n=3 in each group. (I) Confocal microscopy of the whole mount staining with phalloidin (labeling F-actin) revealed the accumulation of F-actin in the vicinity of the myotendinous junction in *klhl41b* mutants (white arrows). n= 15-20 in each group. (J) Confocal microscopy of the whole mount staining with laminin exhibited reduced staining in klhl41b mutants (while arrow) compared to control at MTJ. (K-L) Transmission electron microscopy of myotome at 24 hpf showed the presence of 3-4 rows of myofibrils in each myotube in controls compared to single rows of myofibrils in the *klhl41b* mutants (n=3). (M) Quantification of myofibril rows in each myofiber (n=20), number of sarcomeres in each myofiber (n=20), sarcomere length and height (n=100) on control and klhl41b mutants showed a significant decrease in all parameters. Data are mean ± S.E.M (unpaired t-test, parametric). **** p<0.001; *** p<0.01; ns: not significant.

The sarcomere length in the *klhl41b* mutants was also reduced compared to controls (Figure 2F). Moreover, myotubes without any myofibril rows showed the presence of electron-dense protein aggregates in the vicinity of myotendinous junctions (Figure 2B, black arrows) or the presence of filamentous protrusions emanating from myotendinous junctions (Figure 2H). Whole-mount immunofluorescence with phalloidin showed that these protein aggregates and filamentous structures are composed of F-actin (Figure 2I, white arrows). The myotendinous junction in the controls contained electron-dense collagen-rich fibrils and finger-like projections that were mostly blunted in the *klhl41b* mutants (Figure 2G-H, white arrows). Whole-mount immunofluorescence showed a decrease in laminin signal at MTJ in *klhl41b* mutants compared to controls, suggesting a defect in the formation of MTJ (Figure 2J). Within a few hours (24hpf), control myotubes exhibit 3-4 rows of myofibrils along the longitudinal periphery in myotubes. In contrast, *klhl41b* mutant muscles showed mostly a single row of myofibril in the myotubes (Figure 2K-L). The quantification of sarcomeres in a series in each row of myofibril showed a reduced number of sarcomeres in *klhl41b* mutants compared to the control group (Figure 2I). Quantification of sarcomere size showed a small but significant decrease in sarcomere length in the mutant muscle. Sarcomere height in the mutant myofibrils showed a much more severe decrease in sarcomere size than controls (Figure 2I).

These findings suggest that KLHL41 is required for the initiation of the myofibril assemblies and the addition of new sarcomeres during muscle growth in the existing myofibrils after myoblast fusion. In contrast to previously published models of myofibrillogenesis, myofibrils are added in rows during skeletal muscle development and not as immature actin assemblies or premyofibrils reported using *in vitro* models. As normal muscle contraction is required for MTJ growth and morphogenesis [53], these defects in myofibril formation and skeletal muscle contractions may eventually result in MTJ structural defects, as seen in *klhl41b* mutants.

### KLHL41 is required for the precise alignment of sarcomeres after primary myogenesis

Primary myogenesis in zebrafish skeletal muscle is completed by 24 hpf, establishing functional, contracting muscle with multiple rows of myofibrils within myotubes. As development progresses, new myofibrils are dynamically added, and new sarcomeres are added to the existing myofibrils during secondary muscle growth. As new sarcomeres are added at myofiber ends during skeletal muscle development, we examined the myofibril ends that are attached to the myotendinous junction as development proceeds (3 dpf) [54]. While most of the control myofibrils showed normal sarcomeric organization at the myofibril ends attached to the MTJ, *klhl41b* mutants revealed a lack of discreet attachment sites and disorganized sarcomeres with thickened and disrupted Z-lines in the sarcomeres (Figure 3A-B, black arrows). By 5 dpf, these regions of myofibrils in close proximity to MTJ showed highly misaligned sarcomeres rather than maintaining the parallel alignment observed in control myofibers (Figure 3C-D, white arrows). Most of these sarcomeres lacked well-developed structures and exhibited the presence of electron-dense bodies similar to nemaline bodies seen in skeletal muscles in NM patients (Figure 3D, inset). As muscle development progresses, MTJs typically form finger-like projections that increase the interface surface between terminal sarcomeres and the myoseptum, enhancing mechanical strength and force transmission as seen in the control embryos. In contrast, *klhl41b* mutants exhibited severely disrupted MTJ morphology, with thin, irregularly serrated junctions devoid of deep finger-like extensions (Figure 3E-F, white arrows).

**Figure 3.**
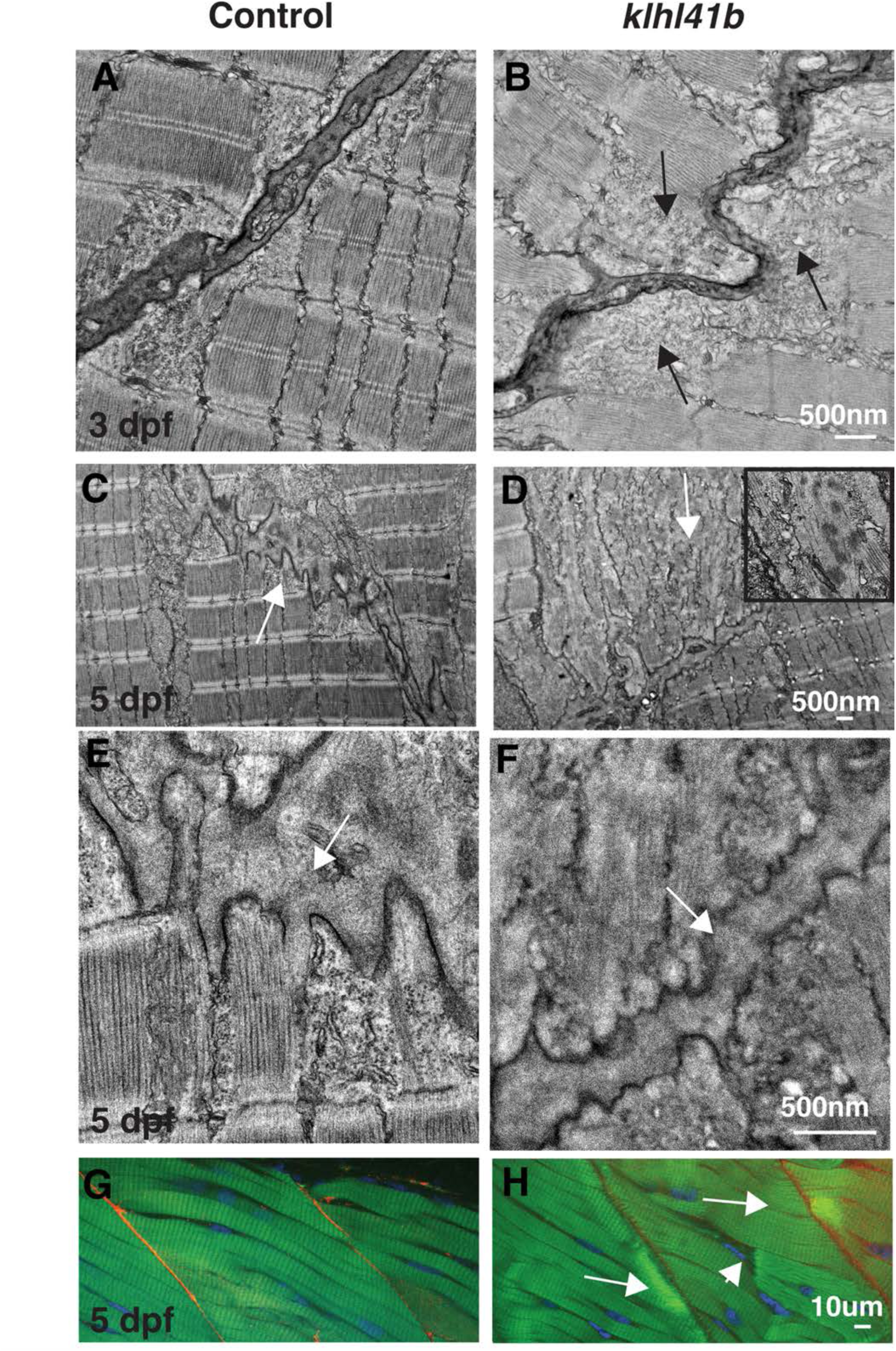
KLHL41 regulates addition and orientation of sarcomere during skeletal muscle growth. (A-B) Control myofibrils are attached to MTJ through terminal sarcomeres that showed damaged sarcomeres in the terminal region in klhl41b mutants at 3 dpf (black arrows). (C-D) Transmission electron microscopy of myotome at 5 dpf revealed the presence of misaligned sarcomeres in proximity to MTJ in *klhl41b* mutants compared to the parallel orientation of control sarcomeres. Moreover, the skeletal muscle showed the presence of electron-dense bodies in the skeletal muscle (n=3). (E-F) MTJ in controls showed finger-like projections, whereas *klhl41* mutants lacked these projections. (G-H) Confocal microscopy of the whole mount immunofluorescence in zebrafish with phalloidin (F-actin, green) and laminin (red) revealed the accumulation of F-actin in the periphery of the myotendinous junction and reduced signal for laminin (n=20). Data are mean ± S.E.M (unpaired t-test, parametric). **** p<0.001; ** p<0.05; ns: not significant.

Further examination of the myotome by whole mount phalloidin (green) and laminin (red) staining showed the accumulation of F-actin at the periphery of myofibers as well as overly stretched broken myofibers in *klhl41b* mutants (Figure 3G-H). Moreover, the intensity of laminin staining was also reduced in *klhl41b* mutants compared to controls. To investigate the effect of structural changes on force production, muscle physiological analysis was performed. Compared to controls, *klhl41b* mutant showed lower force production at all frequencies (1-200Hz) (Figure S3). We also examined the kinetics of the force production by measuring the activation and relaxation kinetics of single twitches at optimal sarcomere length at 1 Hz frequency. While no significant changes were seen in the activation kinetics of peak force development, a significant increase was observed in the relaxation kinetics, showing an increase in the amount of time for the peak force to reduce to the baseline level in the mutant muscle. This suggests that KLHL41 is required for the assembly and precise orientation of sarcomeres into mature myofibrils and MTJ architecture, and a defect in this process contributes to reduced force generation in Klhl41b deficient skeletal muscle.

### Klhl41b deficiency results in changes in sarcomeric, translational and proteolytic proteome in skeletal muscle

KLHL41 is a CUL3 E3 ubiquitin ligase interacting substrate-specific adapter that regulates the stability of skeletal muscle proteins by ubiquitination. To examine how the protein composition of skeletal muscle is altered in KLHL41 deficiency during skeletal muscle development, we performed mass spectrometry (MS)-based quantitative proteome of skeletal muscle from control and *klhl41b* mutant embryos (3 dpf) (Figure 4A). Proteins with a significant differential response between control and mutant samples (Log2Fold change=1.25, p<0.05) were analyzed by the String pathway analysis to reveal enriched proteome abundance patterns across experimental groups (Figure 4B-C). A comparison of the lower abundance proteins in the mutants in functionally enriched categories identified striated muscle contraction as the most enriched group. Within this group, the lowest abundance in mutants compared to controls was observed for the members of the skeletal muscle troponin complex: Tnnt3a, Tnnt1, Tnnc2, Tnni1a, Tnni1b, and Tnni2a, Tnni2b suggesting that KLHL41 is required for the stability of the troponin protein complex (Figure 4D). Nemaline myopathy is caused by mutations in genes regulating or forming the structure of the actin thin filament proteins; therefore, the *klhl41b* mutant proteome was also analyzed for changes in other genes that cause NM. The results showed a downregulation of Neb, Tpm2, and Klhl40a in the mutants, while *A*cta1, Lmod3, Kbtbd13*, and* Cfl2 exhibited no significant differences. In contrast, Tpm3 *and* Mypn were slightly elevated in *klhl41b* mutants compared to controls (Table S3). In addition, other protein groups that showed lower abundance in the *klhl41b* mutants in comparison to controls were associated with phospholipid metabolic pathway and translation processes.

**Figure 4.**
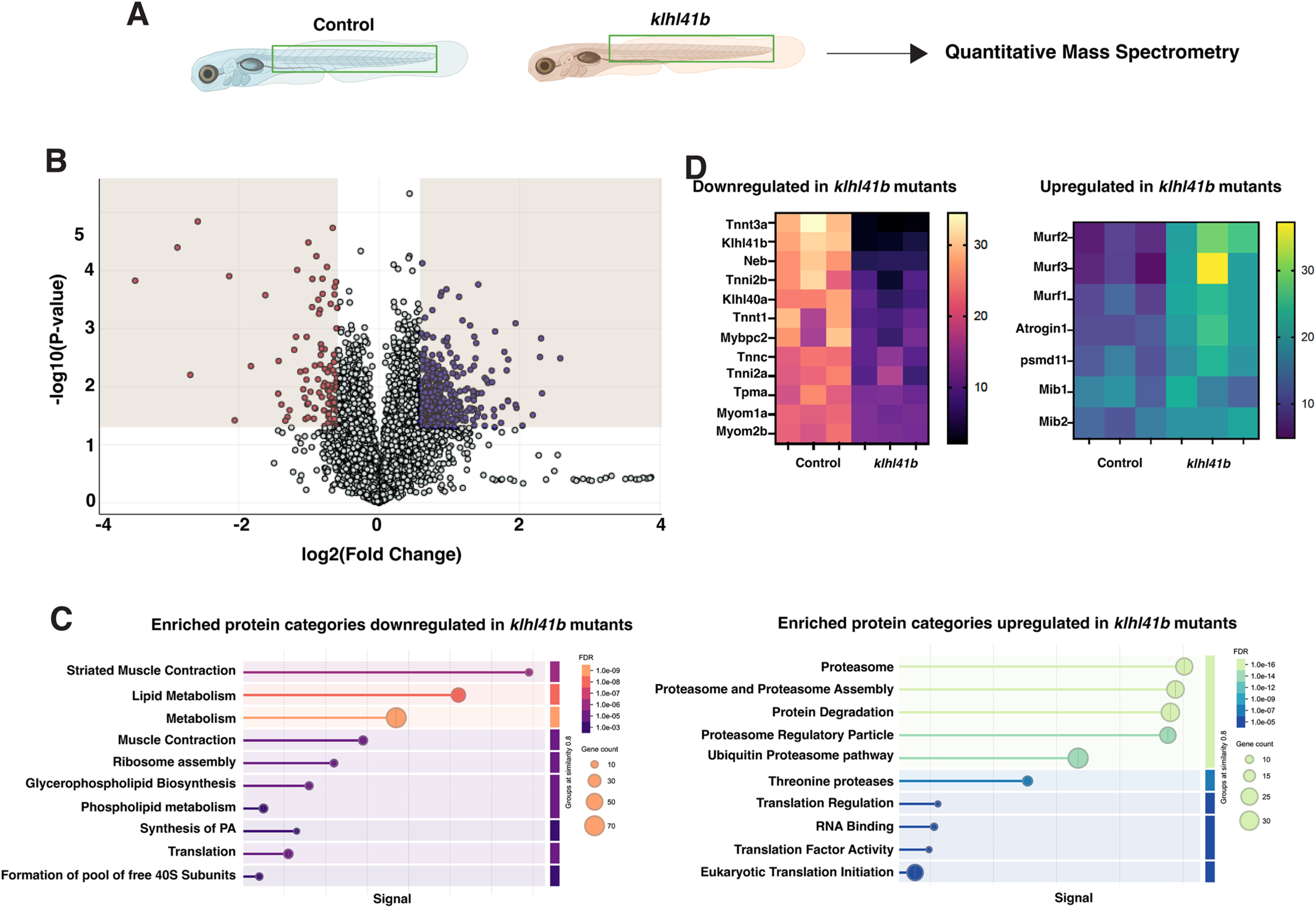
Proteomic changes in KLHL41 deficiency in skeletal muscle. (A) Skeletal muscle tissue from control and *klhl41b* mutants was dissected (3 dpf) and proteome was analyzed by quantitative mass spectrometry (n=3 replicates of 50 embryos in each group). (B) Volcano plot showing protein abundance in control vs *klhl41b* mutant groups (cut off Log2 fold change=1.25 and p<0.05). (C) Reactome pathway enrichment in control or *klhl41b* mutants revealed downregulation of sarcomeric contraction, metabolic and translational pathways in mutants while proteasomal degradation and translational pathways proteins are more abundant in the *klhl41b* mutants compared to controls. (D) Proteins that are down regulated (left,) or upregulated (right) in the *klhl41b* mutants in the top enriched categories.

Conversely, mutant muscle revealed a significant increase in proteasome-associated proteins, including several skeletal muscle-specific E3 ubiquitin ligases implicated in protein degradation during atrophy such as *Murf1, Murf2, Murf3, Atrogin,* and *Mib* (Figure 4C-D). Proteins that showed an increased abundance in *klhl41b* mutant muscles also showed enrichment in translation-related processes, with most proteins in these groups involved in ribosome assembly and translational initiation, suggesting a disruption in the translation processes in *klhl41b* mutants.

### KLHL41 regulates the stability of the troponin protein complex

Our previous studies have shown that KLHL41 regulates the dynamics of the protein complexes by regulating their protein stability through ubiquitination by the CUL3 E3 ligase-KLHL41 complex [9]. In Klhl41b deficient skeletal muscle, all the members of the troponin protein complex showed a significant downregulation. The Troponin complex is the Ca2+ regulating complex for skeletal muscle contraction and is localized to thin filaments in sarcomeres similar to KLHL41 (Figure 5A) [55]. Mutations in skeletal muscle troponin complex proteins are associated with severe myopathy [29, 56–60]. Therefore, we set out to identify if KLHL41 is a regulator of the stability of the troponin protein complex and, hence, contributes to skeletal muscle weakness. While all skeletal muscle troponin proteins were downregulated in the mutants, the highest down-regulation in the *klhl41b* muscle was observed for fast muscle troponin protein complex (TNNT3, TNNI2, and TNNC2). Therefore, we focused on understanding the effect of KLHL41 on the fast muscle-specific troponin protein complex. To identify any direct interaction between KLHL41 and different troponin protein members, KLHL41 was overexpressed in the C2C12 muscle cells, and co-immunoprecipitation was performed in the differentiating myotubes (day 2; two days in the differentiation media). Western blot analysis revealed that KLHL41 interacts with the endogenous TNNT3 but not with the TNNC2 or TNNI2 (Figure 5B). As KLHL41-CUL3 complex is shown to regulate the stability of the sarcomeric protein by ubiquitination, we also analyzed the ubiquitination of TNNT3 in the presence of KLHL41 and CUL3 ligase in C2C12 myotubes (Figure 5C). However, no ubiquitination was observed for TNNT3 in the presence of KLHL41 and CUL3.

**Figure 5.**
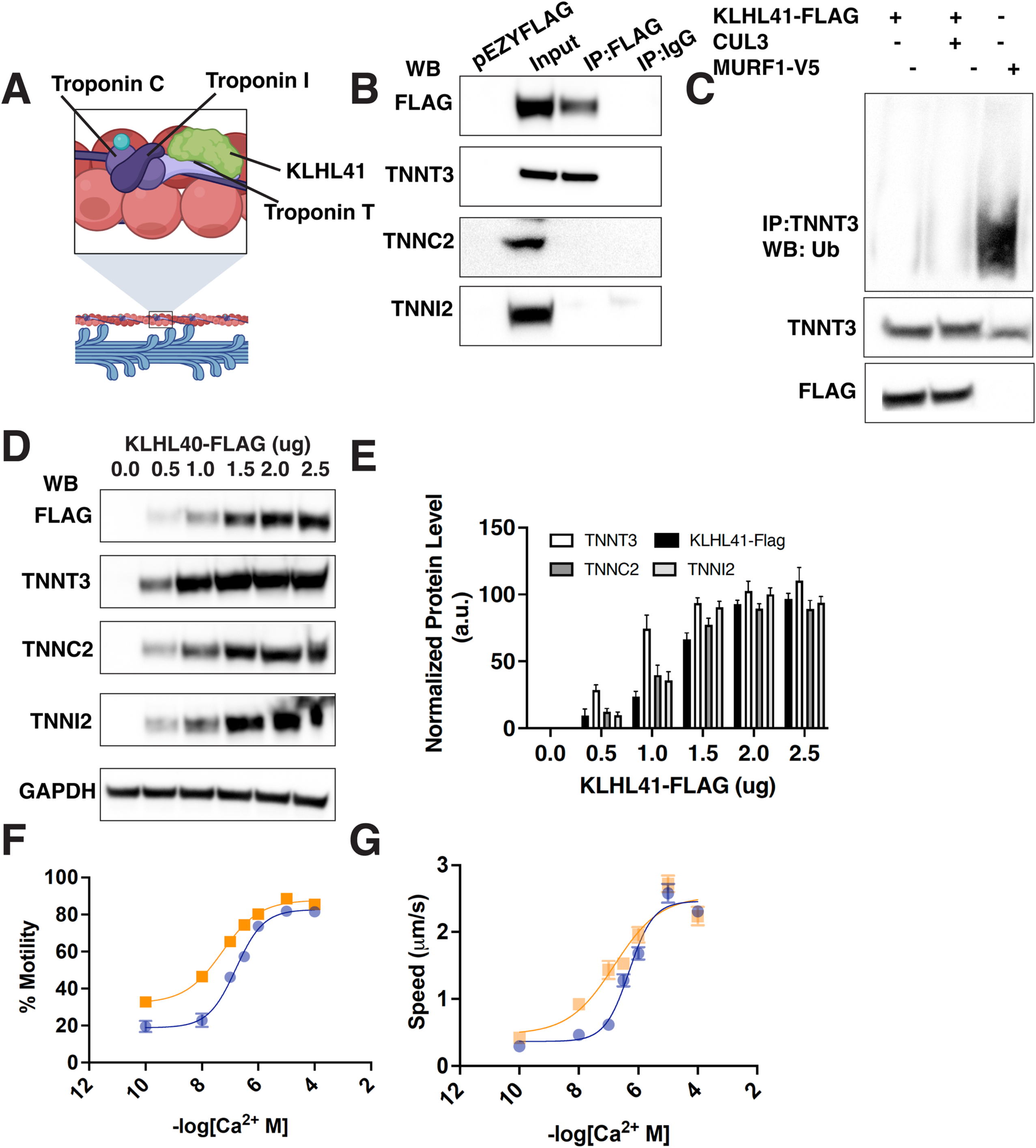
Troponin protein complex is destabilized in Klhl41 deficiency and leads to increased Ca2^+^ sensitivity and reduced cooperativity. (A) Troponin protein complex. Red: thin filaments, blue: thick filaments, purple and green: troponin proteins in skeletal muscle. (B) Coimmunoprecipitation in C2C12 myotubes revealed that KLHl41 directly interacts with TNNT3 but not with other members (TNNC2 and TNNI2) in the troponin protein complex. (C) KLHL41-FLAG with or without CUL3 E3ubiquitin ligase was expressed in C2C12 cells and immunoprecipitation was performed with TNNT3 antibody. MURF1-V5 was also overexpressed as a positive control for TNNT3 ubiquitination. Western blot of immunoprecipitated protein with ubiquitin antibody revealed the absence of TNNT3 ubiquitination in the presence of KLHL41 and CUL3, while ubiquitinated TNNT3 was seen in the presence of MURF1-V5. (D) Western blots of protein extracts from C2C12 myotubes demonstrate the expression of increasing amounts of KLHL41-FLAG in C2C12 myotubes, which results in an increase in the protein levels of all troponin protein members. (E) Quantification of the protein levels by Western blots. N= 3 independent experimental replicates. Data are mean ± S.E.M; unpaired t-test, one-way analysis of variance (ANOVA) and Tukey’s HSD test for multiple groups comparison. (F) The sliding speed of fluorescently labeled native thin filaments with native bound Troponin and Tropomyosin purified from either control or *klhl41b* was measured with increasing calcium concentration. Fit parameter*: pCa_50_* (control: 6.31 ±0.29, klhl41b: 6.803 ±0.91), *n_H_* (control: 1.07 ±0.6131, *klhl41b*: 0.5655 ±0.6044), *Offset* (control: 0.364 ±0.3145, *klhl41b*: 0.4740 ±0.781), *V_max_* (control: 2.461 µm*sec^-1^ ±0.299, *klhl41b*: 2.535 µm*sec^-1^ ±0.54) (G) Percent of motile filaments plotted against calcium concentration purified for control zebrafish or *klhl41b* zebrafish. Fit parameters for panel B: *pCa_50_* (control: 6.793 ±0.293, *klhl41b*: 7.273 ±0.465), *n_H_*(control: 0.8864±0.4658, *klhl41b*: 0.6535 ±0.372), *Offset* (control: 18.85%±9.49, *klhl41b*: 32.12% ±11.32), and *Max % Motile* (control: 82.68 % ± 6.9, *klhl41b*: 87.78 % ± 6.78).

TNNT3 has previously been reported to be a target of MURF1 E3 ubiquitin ligase and therefore, MURF1 was used as a positive control. Immunoprecipitation of TNNT3 in C2C12 myotubes showed ubiquitinated TNNT3 in the presence of MURF1. This suggests that KLHL41 regulates the stability of the troponin protein complex independent of the CUL3-mediated ubiquitination.

Next, we evaluated the effect of KLHL41 protein on TNNT3 protein stability. Increasing amounts of KLHL41-FLAG plasmid were overexpressed in C2C12 myotubes, and western blotting was performed to quantify the level of the troponin protein complex (TNNT3, TNNI2, and TNNC2) (Figure 5D). Increasing the amount of KLHL41 protein levels led to an increase in TNNT3 protein levels in the differentiating myotubes.

Similarly, an increase in KLHL41 also increased TNNC and TNNI levels (Figure 5D-E). This suggests that KLHL41 is required to stabilize the troponin protein complex in differentiating myotubes by direct interaction with TNNT3.

### Calcium dysregulation in KLHL41 deficiency

The troponin protein complex serves as the primary regulator of skeletal muscle contraction. When calcium binds to troponin C in the complex, structural changes occur within the thin filaments, revealing myosin binding sites on actin and allowing cross-bridge formation. Therefore, to assess if the reduced abundance of troponin complex members affects cooperative cross-bridge formation in *klhl41b* mutants, *in vitro* motility assays were performed on thin filaments purified from control or *klhl41b* mutant fish (Figure 5F-G). The calcium sensitivity of thin filaments was quantified as the pCa₅₀ value, which represents the calcium concentration required for half-maximal filament speed. The Hill coefficient (nH) was also calculated to assess the cooperativity of thin filament activation. As shown in Figure 5, the motility of thin filaments purified from control (wild-type) animals was inhibited at low calcium concentrations. Increasing calcium levels resulted in a greater fraction of motile filaments and increased filament speed. Similarly, thin filaments purified from *klhl41b* mutants were activated at high calcium concentrations, showing no significant difference in maximum filament speed compared to Tpm-WT (Figure 5G). However, unlike control thin filaments, *klhl41b* mutant thin filaments did not fully inhibit motility at low calcium, leading to a significant increase in pCa₅₀. Additionally, the *klhl41b* mutant thin filaments exhibited reduced cooperativity, as indicated by a significantly lower Hill coefficient (Figure 5), suggesting that thin filament activation is more gradual in response to increasing calcium levels in the *klhl41b* mutants than controls.

The increase in pCa50 observed in *klhl40b* mutant zebrafish is consistent with an impairment in skeletal muscle activation and relaxation by calcium. During normal swimming, zebrafish generate alternating contractions along the body axis. For this to be efficient, each muscle contraction must be coordinated with the relaxation of the opposing muscle groups to allow full bending to each side. In the *klhl40b* mutant, the increased calcium sensitivity suggests that mutant muscles might remain partially contracted for longer periods of time, reducing their ability to relax fully between contraction cycles consistent with shorter sarcomeres (Figure 2M). As a result, when one side of the fish contracts, the opposing side does not fully relax, limiting the degree of bending, consistent with the reduction in tail curvature observed *in vivo* in the *klhl41b* but absent in WT zebrafish (Figure 1F). Thus, the impaired activation and/or relaxation of thin filaments isolated from mutant larvae provide a potential mechanistic explanation for the reduced curvature amplitude seen *in vivo* swimming.

### KLHL41-Troponin complex is required for sarcomere assembly and alignment

*TNNT3* mutations have been previously reported in arthrogryposis with neonatal lethality and congenital nemaline myopathy, but suitable vertebrate animal models are unavailable to investigate the function of TNNT3 in skeletal muscle development. The function of skeletal muscle troponin is mostly postulated from studies on cardiac TNNT2 in regulating muscle contraction by binding Ca^2+,^ but its role in sarcomere formation is not known. Therefore, to understand if the TNNT3 deficiency in *klhl41b* mutants affects the Ca^2+-^mediated muscle contraction or also contributes to sarcomere development in developing myotubes through interaction with Klhl41, we studied the knockout zebrafish model of human *TNNT3* orthologue (*tnnt3a*). *tnnt3a* knockout fish (c.A529T; NM_131565.1 and p.k177*) were obtained from the Zebrafish International Resource Center and predicted to result in a truncated protein (Figure 6A). Ultrastructure analysis of the skeletal muscle of *tnnt3a* mutants (2 dpf) by electron microscopy revealed a reduced number of myofibril rows in myotubes compared to controls, suggesting that Tnnt3a is required for myofibril formation in developing skeletal muscle (Figure 6B-C, H). Control myotubes contain sarcomeres organized in a parallel orientation. Skeletal muscle in *tnnt3a* mutants exhibited relatively normal sarcomere in the middle of the myofibers with misaligned sarcomeres at myofibril ends in the proximity of MTJ, similar to *klhl41b* mutants (Figure 6D-E, white arrows). Moreover, Z-lines in these sarcomeres are irregular and thicker than controls (Figure 6E, black arrows), as seen in the skeletal muscle of patients with nemaline myopathy [61] . The terminal sarcomeres at the myofibril end connecting to MTJ appear to be thinner compared to the control, suggesting a defect in addition to new sarcomeres in the *tnnt3a* mutant muscle (Figure 6 F-G). Quantifying the sarcomere size showed a decrease in sarcomere length in the *tnnt3a* mutant compared to control sarcomeres (Figure 6I). No significant differences in sarcomere height between control and tnnt3a myofibers were observed except at the myofiber ends (Figure 6J-K). The direct interaction between KLHL41 and TNNT3 and a similar phenotype in skeletal muscle suggests that the KLHL41-TNNT3 complex is critical for adding new sarcomeres and their precise alignment in myofibrils.

**Figure 6.**
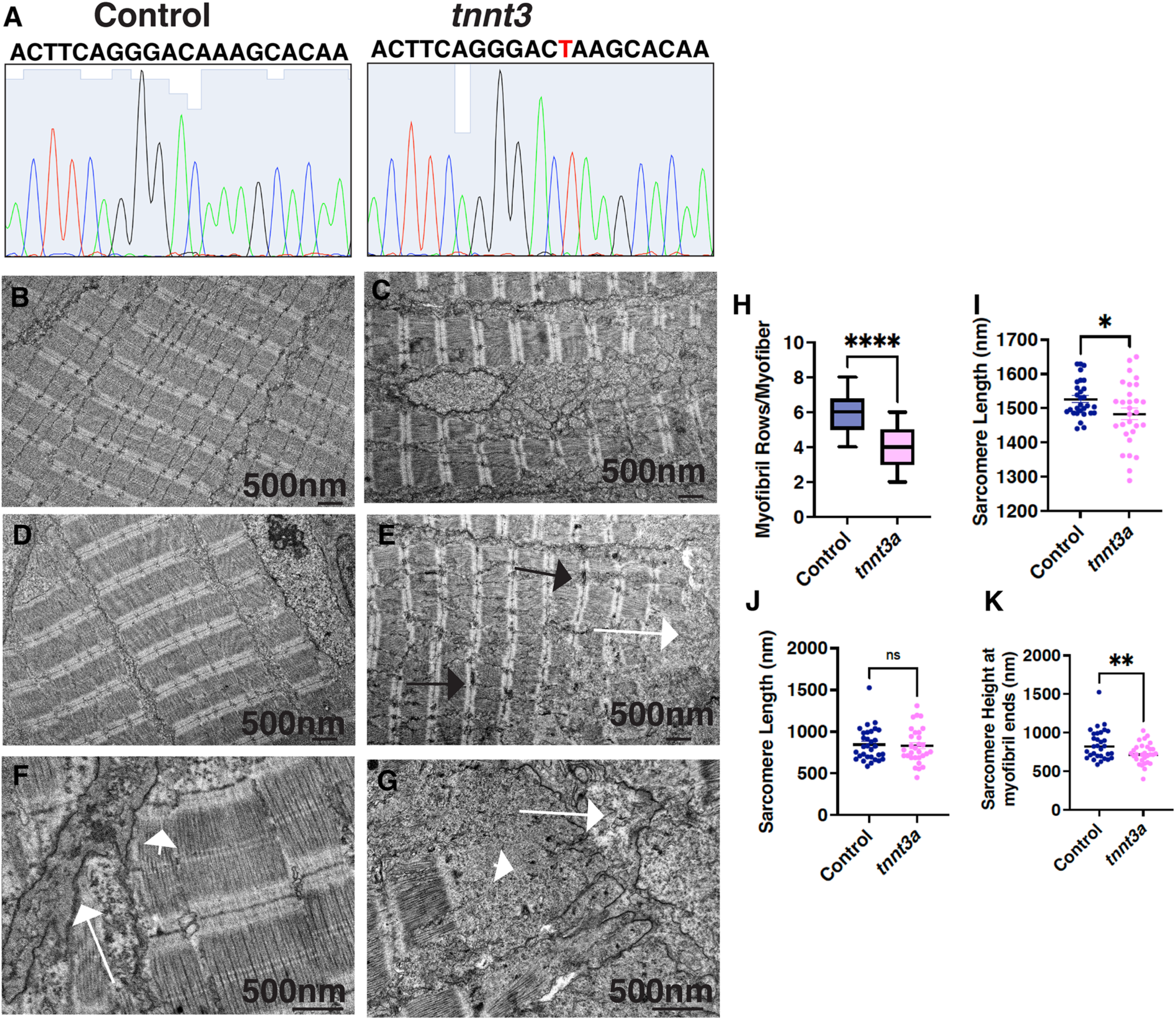
Troponin complex is required for additions of myofibril rows and new sarcomeres during skeletal muscle growth. (A) *tnnt3a* mutant fish harbors a stop loss of function allele in *tnnt3a* gene that is predicted to result in a truncated protein. (B) *tnnt3a* mutants exhibit reduced rows of myofibrils compared to controls (2 dpf). (C-D) sarcomeres in control skeletal muscle are arranged parallelly, while *tnnt3a* mutant sarcomeres lacked parallel organization and exhibited disorganized thicker z-lines (black arrows) in proximity to the MTJ areas (white arrows). (F-G) Myofibers are attached to MTJ by terminal sarcomeres in control but exhibit wide regions lacking sarcomeres (white arrow) and contain the presence of electron-dense bodies near MTJ (white arrowhead). (H-I) quantification of reduced rows of myofibrils in each sarcomere and reduced sarcomere length in the *klhl41b* mutants. Sarcomere height showed a significant difference in sarcomeres near MTJ areas but not in other regions of the myofibers compared to controls. Data are mean ± S.E.M (unpaired t-test, parametric). **** p<0.001; ** p<0.05; ns: not significant.

### Disorganized sarcomere alignment signals the onset of muscular atrophy

*Klhl41b* mutant proteome revealed enrichment of proteins regulating proteolytic processes as well as high abundance in the *klhl41b* mutant muscle compared to controls. Muscular atrophy is widely associated with myopathies and dystrophies, but the origin of atrophic changes in skeletal muscle is mostly unknown. Most of the proteolytic enzymes and skeletal muscle E3 ubiquitin ligases are present in the sarcomere, suggesting that sarcomeric defects may signal the activation of the skeletal muscle atrophy program in defective myofibers. To test this hypothesis, we analyzed the expression of these atrophy genes in different neuromuscular disease-causing zebrafish mutants with exhibiting sarcomeric defects (*klhl41b*, *tnnt3a*) and dystrophic mutants (*dmd*, *dag1*) that exhibit normal sarcomeres during early development (2 dpf). RT-PCR analysis revealed a significant upregulation of *murf1, murf2, murf3, atrogin, and mib1* transcripts in *klhl41b* mutants, suggesting that the regulation of atrophy genes is controlled at the transcriptional level (Figure 7). Similarly, these atrophy genes showed a significant increase in mRNA level in Tnnt3a deficiency. *Sapje* fish exhibit normal skeletal muscle development followed by skeletal muscle degeneration after 3 dpf. Similarly, *dag1* zebrafish mutants exhibit normal sarcomere structure during early development, as reported previously. No differences in transcript expression for murf1, murf2, murf3, atrogin, and mib1 was observed in *spaje* or *dag1* mutants compared to controls, suggesting that structural perturbation in sarcomere results in the upregulation of atrophy genes further contributing to skeletal muscle weakness in sarcomeric diseases.

**Figure 7.**
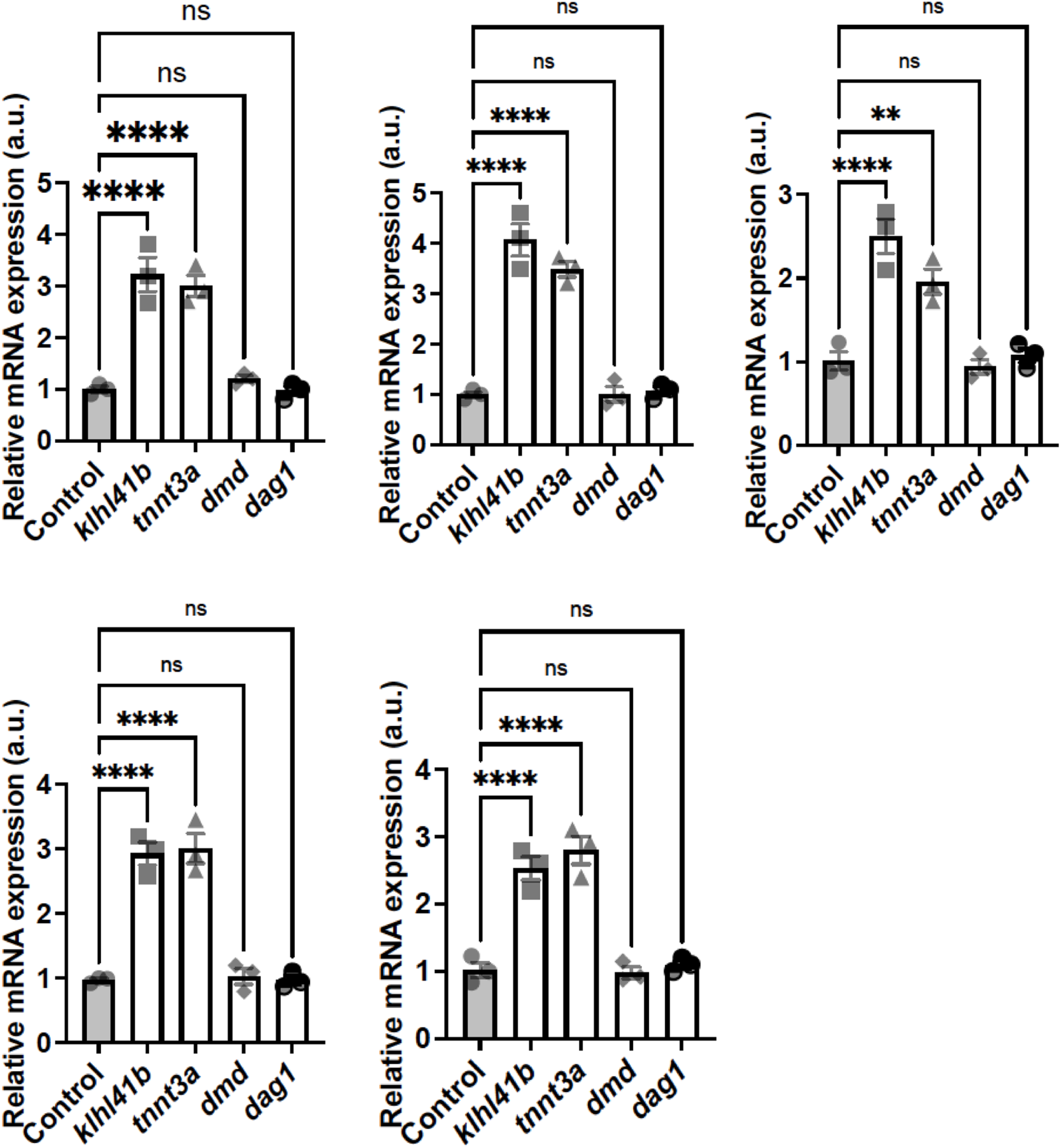
**Sarcomeric disruption in skeletal muscle is associated with increased expression of skeletal muscle atrophy genes**. Quantitative RT-PCR analysis of control and klhl41b, tnnt3, dmd and dag1 mutants for skeletal muscle atrophy transcripts (3 dpf) (n=20 in each group). Data are mean ± S.E.M; with one-way analysis of variance (ANOVA) and Tukey’s HSD test. **** p<0.001; ** p<0.05; ns: not significant.

## DISCUSSION

Mutations in approximately 200 genes contribute to the diverse array of sarcomeric diseases affecting sarcomere assembly, maintenance, or growth (https://www.musclegenetable.fr/). However, no therapies are available for these disorders due to a lack of understanding of how specific sarcomeric defects contribute to skeletal muscle weakness. Therapeutic development of sarcomeric diseases is challenging as many structural proteins encoding genes are extremely large, while other sarcomeric genes result in rare or ultra-rare forms of myopathies. By understanding the specific sarcomeric defects in vertebrate animal models of human myopathies, we may devise better therapeutic strategies that may have a wider application by restoring the structure and function of sarcomeres for these diseases.

Nemaline myopathy is a sarcomeric disease where mutations in 11 genes, including *KLHL41,* result in disorganized sarcomeres, the presence of nemaline bodies, and hypotonia in affected patients. However, it is still unclear if the sarcomeric defects observed in patients are due to a defect in the formation or maintenance of myofibrils.

Previous studies on KLHL41 deficient mice, zebrafish, and cells showed sarcomeric defects in skeletal muscle, but the mechanism is not clear [32, 62, 63]. In this work, we identified that KLHL41 is required for the initiation of sarcomeric assembly and regulating sarcomere size *in vivo*, which was also reflected in reduced spontaneous coiling in mutants during early development as muscles start to contract after establishing initial sarcomere assemblies. Similarly, many KLHL41 patients and congenital NM patients with mutations in other genes experience fetal akinesia and reduced fetal movement, suggesting muscle weakness is initiated during fetal development.

We identified that sarcomeres are assembled as single rows of myofibrils adjacent to the sarcolemmal membrane and attached to MTJ on both ends. These first-formed sarcomeres exhibit well-defined Z lines, thick and thin filaments, A and I-bands, and an H zone similar to sarcomeres in the adult muscle. This shows that sarcomere assembly *in vivo* does not require intermediate steps of immature actin assemblies or pre-myofibril formation seen *in vitro* models. This suggests that vertebrate myotubes inherently contain all the necessary proteins to assemble mature sarcomeres rapidly and efficiently to support the primary myofibrillogenesis within a few hours of myotubes’ attachment to the MTJ. After the first rows of myofibrils is established, subsequent rows are added dynamically adjacent to these rows, thus filling the core of the myotubes with myofibrils by 48 hpf. KLHL41 deficiency results in delayed onset of the first row of myofibrils as well as the addition of subsequent rows. We and others have shown that KLHL41 is required to form functional sarcomeres by regulating the protein stability of other key sarcomeric proteins such as nebulin or NRAP [9, 62]. However, KLHL41 may have additional roles in the initiation of sarcomeric assembly, as seen by delays in the formation of the first rows of sarcomeres. Our proteomic analysis identified the reduced abundance of proteins required for protein translation machinery. This suggests that changes in the ribosome biosynthesis and translation initiation program in the Klhl41b proteome may affect the translation of sarcomeric and other regulatory proteins required to form sarcomeres. Recent work has shown that β_2_-adrenergic-mediated muscle hypertrophy, which causes increased ribosomal protein production, is significantly impaired in KLHL41 deficiency in a cellular model [64]. While further studies may identify the specific role/s of KLHL41 in regulating sarcomere assembly *in vivo*, our data suggests KLHL41 may also contribute to sarcomere initiation by regulating functional ribosomal assembly and function.

Klhl41 deficient myofibrils displayed a significant reduction in the sarcomere height from the beginning of myofibrillogenesis. Regulation of sarcomere length is well studied, and many sarcomeric proteins such as LMOD3, TMOD, CAPZ, NEB, tropomyosins, CFL2, and TTN contribute to different steps of sarcomere elongation by regulating the assembly and stability of actin thin filaments [19, 65–69]. Sarcomere height is a critical contributor to radial myofiber growth and skeletal muscle hypotrophy, but how sarcomere height is regulated during normal muscle development and growth is not clear. As skeletal muscle hypotrophy is associated with increased sarcomere height, we compared the control and *klhl41b* mutants’ proteome for change in these pathways. No significant changes were observed in key regulators of skeletal muscle hypertrophy pathways such as Igf1r, mTOR pathways, and TGFβ in the proteome of *klhl41b* mutants compared to controls. As the skeletal developmental proceeds, *klhl41b* mutants exhibited a delay in adding new myofibrillar rows. Our studies showed that KLHL41 interacts with TNNT3 in the differentiating myotubes and *tnnt3a* mutants stabilizes TNNT3. While no evidence of KLHL41-mediated ubiquitination was found on the TNNT3 protein, this interaction may prevent TNNT3 from other ubiquitination enzymes such as MURF1 and protein degradation in the muscle cells, resulting in the increased stability of TNNT3. The troponin protein complex plays a central role in calcium-mediated skeletal muscle contraction, and our studies show that, in addition to regulating calcium homeostasis, this complex is critical for sarcomere formation in the developing muscle. These myofibrillar defects in *tnnt3a* mutant zebrafish show that, like in *klhl41b* mutants, muscle hypotrophy and weakness in congenital NM are caused by sarcomere formation defects. Similarly, deficiency of cardiac Troponin T results in defective sarcomeric assembly in zebrafish hearts with downregulation of other troponin complex members. However, the precise mechanism by which cardiac troponin T results in sarcomeric defects is not clear [70]. As the troponin complex is the key protein complex on thin filaments, deficiency of TNNT3 may prevent the assembly of thin filaments and, therefore, functional sarcomeres in developing skeletal muscle.

Moreover, TNNT3 may function together with KLHL41 in maintaining the regulation of other molecular processes, such as ribosomal function and muscle atrophy processes that are perturbed in *klhl41b* mutant zebrafish thereby resulting in similar sarcomeric defects.

Contractures are usually present with severe clinical forms of NM caused by mutations in other genes; however, most of the KLHL40 and KLHL41 NM patients exhibit contractures irrespective of their clinical severity [32, 71]. Previous work has shown that defects in the addition of new sarcomeres in series result in the development of contractures [72]. We observed the presence of actin-rich protein aggregates at the myofiber ends. Further, we observed the sarcomeric misalignment in the developing skeletal muscle that also begins at the myofiber ends, finally resulting in the development of nemaline bodies in these misaligned sarcomeric regions. Previous studies have shown the presence of subsarcolemmal nemaline bodies in skeletal muscle biopsies in NM patients [61]. Similar to *klhl41b* and *tnnt3a* mutants, deficiencies of the NM causing LMOD3 and Cofilin as well as capping protein CapZ also result in protein aggregation and sarcomeric disorganization defects at myofiber ends [66, 67], suggesting that each of these proteins is required for sarcomere addition during muscle growth. While no changes in the skeletal actin protein level were observed in Klhl41b deficiency, we observed abnormal protein aggregates at the myofiber ends in the *klhl41b* mutants. Recent work has shown that actin polymerization-driven membrane protrusions form strong integrin attachment sites in the developing embryos and MTJ defects in *klhl41b*, *tnnt3a* and other NM models could also be due to abnormal actin accumulation and decrease in functional sarcomeres in the MTJ area [73].

Finally, we identified significant upregulation of E3 ubiquitin ligases and other proteins that contribute to skeletal muscle atrophy and dysfunction. Most of these proteins are localized on the sarcomeres in the skeletal muscle and are upregulated during muscle wasting. The specific factors triggering the activation of muscle atrophy in skeletal muscle during disease conditions is not completely understood. Analyzing a number of zebrafish models of neuromuscular diseases revealed that the upregulation of the skeletal muscle atrophy program is signaled by structural defects in sarcomeres. This could be due to changes in the mechanical properties in myofiber thus signaling the nuclei to switch on the atrophic pathways. Changes in cellular metabolites through mitochondrial dysfunction have recently been shown to be regulators of the assembly and activity of the proteasome complex [74]. As mitochondria are localized in close proximity to sarcomeres, any defects in sarcomere structure may also result in changes in the metabolites and signal to activate the protein degradation processes in skeletal muscle and future studies may be able to identify these specific signals.

Therefore, our studies show that KLHL41 is critical for initiating sarcomere assembly and myofibril rows during muscle growth by stabilizing sarcomeric proteins and regulating protein turnover processes (summarized in Figure 8). Moreover, KLHL41 is required for the alignment and addition of sarcomeres in series during muscle growth, a defect that results in hypertrophic muscle with misaligned sarcomeres with abnormal protein aggregates, leading to weak skeletal muscle in NM.

**Figure 8.**
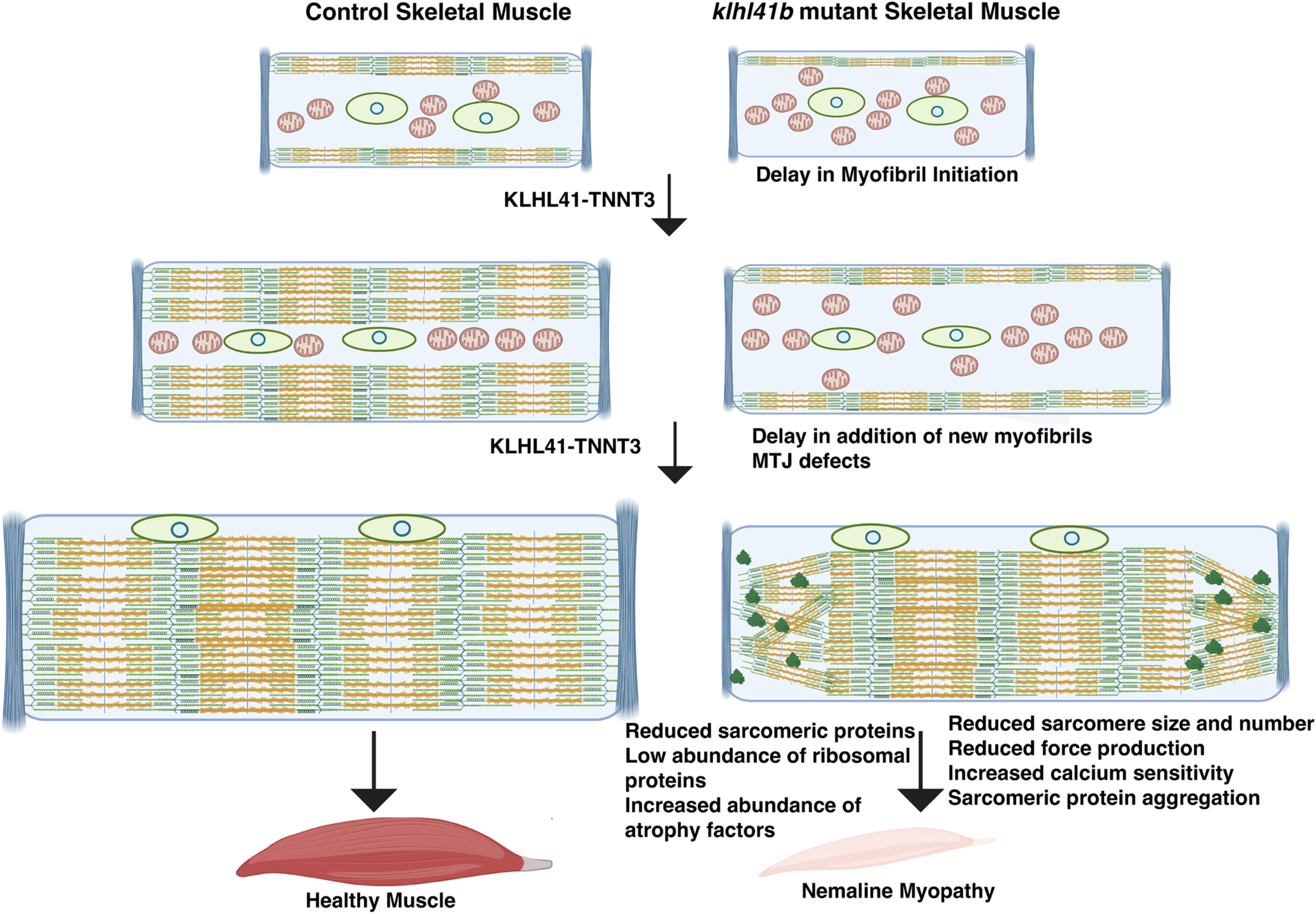
Proposed Role of KLHL41 in Normal and NM skeletal muscle. KLHL41 is required for initial sarcomere formation and subsequent addition of new myofibril rows for radial growth and terminal sarcomere addition for the longitudinal growth of skeletal muscle. Defects in these processes result in hypertrophic skeletal muscle with weakness and reduced force production. Dynamics and stability of sarcomeric proteins and translational and atrophic processes are altered in KLHL41 deficiency and underlie the disease state.

## Acknowledgments

This work was supported by NIH R56AR077017 (VAG) and A Foundation Building Strength grant (VAG); NIH R01AR053897 (HG). We thank Jonathan Van Vranken at the Thermofisher Center for Multiplexed Proteomics at Harvard Medical School for consultation on sample preparation and performing mass spectrometry. We thank Louise Trakimas (HMS Electron Microscopy Core) for her excellent help in the sample preparation and microscopy resources on the North Quad (MicRon) core at Harvard Medical School for confocal microscopy.

---

**Figure S1:** Effect size estimates for 3 pre-stimulus variables, 4 overall performance kinematic variables, and 27 additional kinematic variables classified by escape response stage. Data were obtained using a bootstrap procedure (37) and indicate the estimated difference between genotypes (*klhl41b* minus wild-type) with the 95% confidence interval.

**Figure S2:** Klhl41b is dispensable for myoblast fusion. Confocal image of the whole mount fluorescence in control and *klhl41b* zebrafish embryos (16 hpf) labeled with phalloidin (F-actin green) and DAPI (pink). Quantification of the number of nuclei in each myofiber (n=20 embryos in each group) showed no significant differences between control and *klhl41b* mutants. Data are mean ± S.E.M (unpaired t-test, parametric), ns: not significant.

**Figure S3:** Muscle physiology in control and *klhl41b* mutants. (A) Force production in control and klhl41b mutants at all the frequency range of stimulation resulted in lower force producton in the mutants. (B) Activation and (C) relaxation of single twitch produced at 1 Hz frequence.

**Table S1.** : Oligonucleotides to generate *klhl41* mutant zebrafish lines

**Table S2:** Primer sequences for genotyping zebrafish lines.

**Table S3:** Proteomic changes in the *klhl41b* mutant muscle

