## Supplementary figures and images for "KLHL41 orchestrates sarcomere assembly and size to drive skeletal muscle hypertrophy *in vivo*"

### Supplemenatl Figure 2

#### Supplemental Figure S2

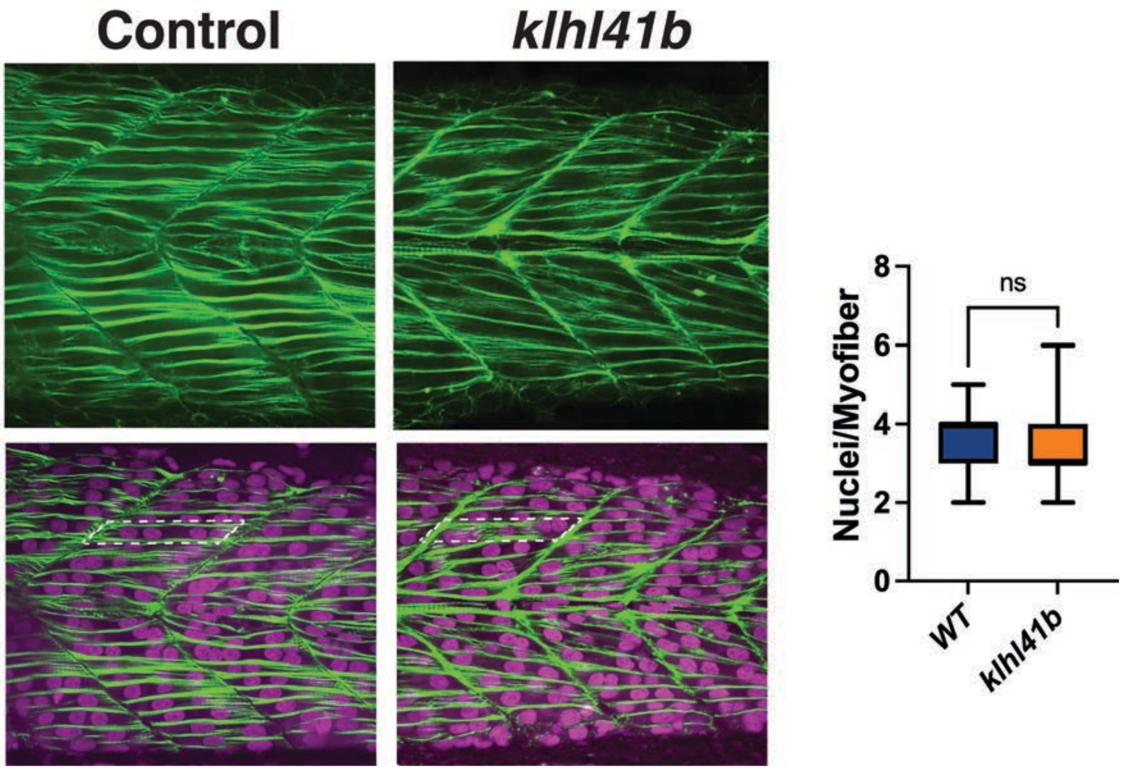

### Supplemental Figure 1

#### Supplemental Figure 1

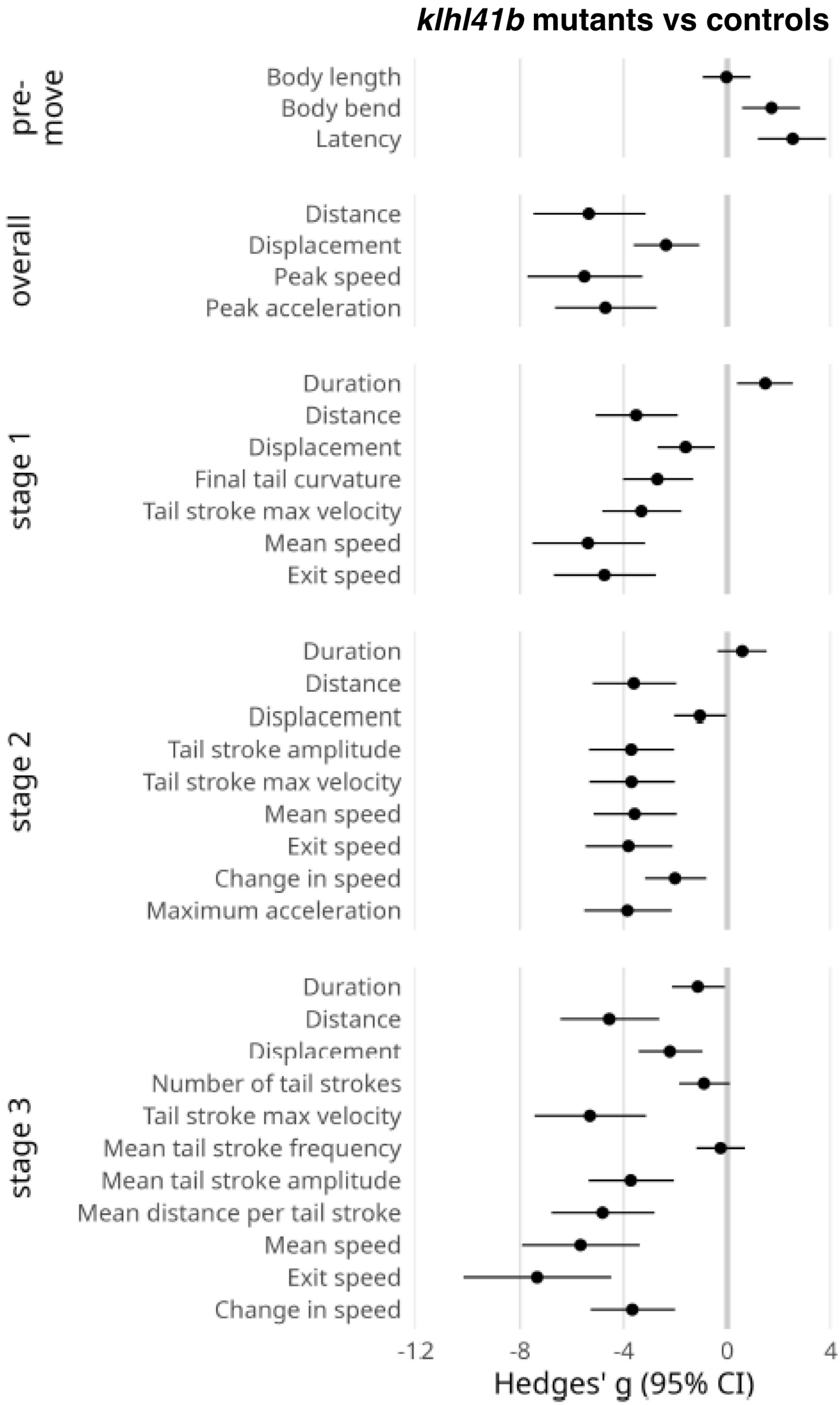

### Supplemental Figure 3

#### Supplemental Figure S3

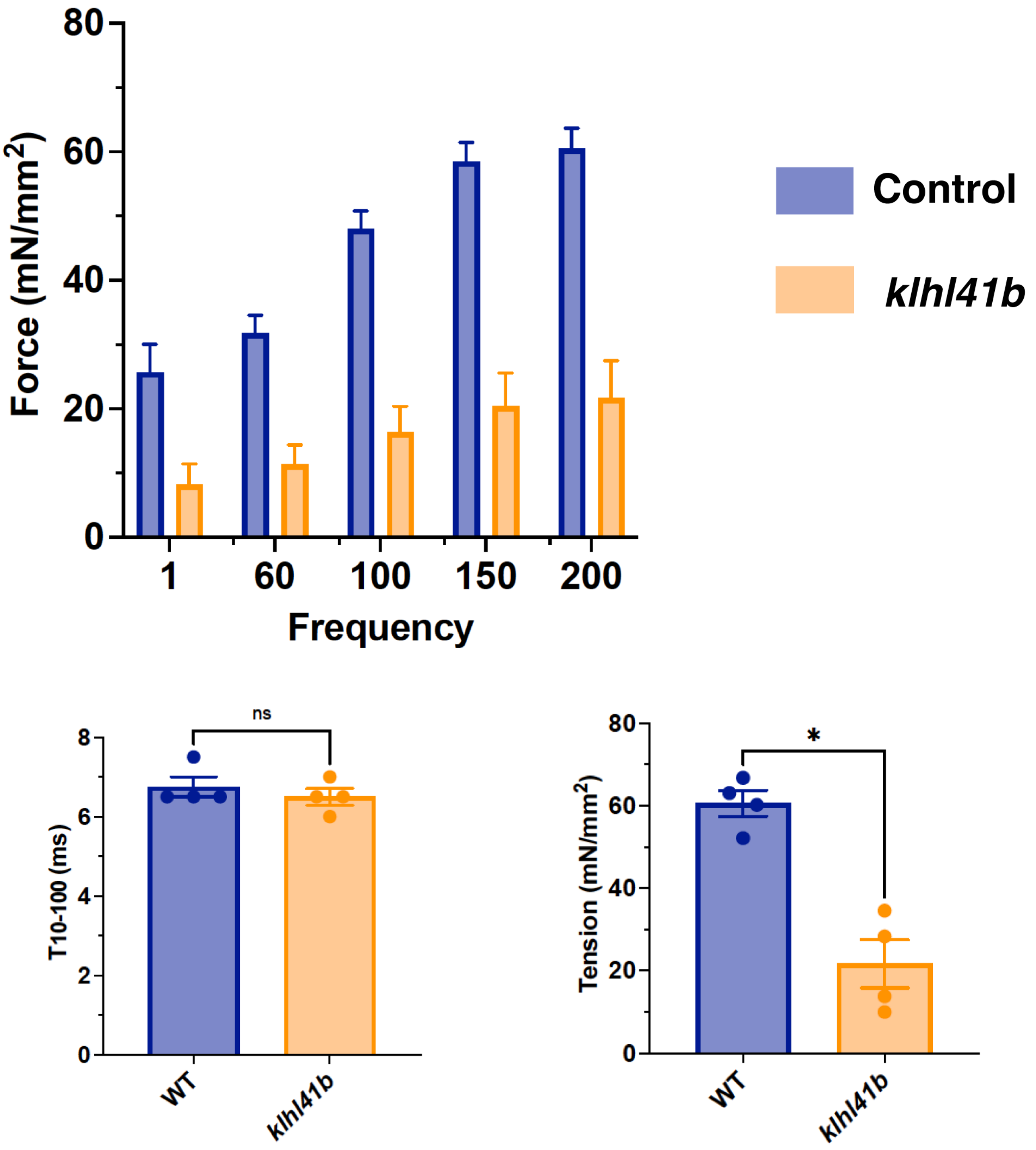
