## Supplemental Table 1 for "KLHL41 orchestrates sarcomere assembly and size to drive skeletal muscle hypertrophy *in vivo*"

**Table S1 : Oligonucleotides to generate *klhl41* mutant zebrafish lines**

| Gene | Target Site | Oligo1 | Oligo2 |
|---|---|---|---|
| *Klhl41a* | GGCTATCTTCAGGATGGGGC | TAGGCTATCTTCAGGATGGGGC | AAACGCCCCATCCTGAAGATAG |
| *Klhl41a* | GGAGGCAGAGCAATCCAGTT | TAGGAGGCAGAGCAATCCAGTT | AAACAACTGGATTGCTCTGCCT |
| *Klhl41b* | GGAAGGAGAGGTGAACGGAG | GGAAGGAGAGGTGAACGGAG | AAACCTCCGTTCACCTCTCCTT |
| *Klhl41b* | GGTGAACGGAGAGGAGGAGG | TAGGTGAACGGAGAGGAGGAGG | TAGGTGAACGGAGAGGAGGAGG |
