## Supplemental Table 2 for "KLHL41 orchestrates sarcomere assembly and size to drive skeletal muscle hypertrophy *in vivo*"

**Table 2 : Primers for genotyping zebrafish lines**

| Gene | Forward primer (5’-3’) | Reverse Primer (5’-3’) |
|---|---|---|
| *Klhl41a ex2* | TTCTTATGAACTCTCCATTACA AGC | TGACACAGTCGGCATGACTCAT AA |
| *Klhl41b* | GCTTCCGACTTCTCCCAGAGA | GGCGACTCTTTGCTTTCTTCA |
| *Tnnt3a* | GAGGACAAACTGAGGTGAGT | CAGGTTACTACACTGCGCAG |
