## Supplemental table 3 for "KLHL41 orchestrates sarcomere assembly and size to drive skeletal muscle hypertrophy *in vivo*"

Table S3: Differentially Regulated proteins between controls and klhl41b mutant muscle

| UniprotID | GeneSymb | Descriptior | NumPeps | neglogPVal | log2FC |
|---|---|---|---|---|---|
| A3KPW9 | Bag6 | Large proli | 2 | 1.620806 | 0.62405 |
| Q6YLH9 | CAV1 | Caveolin | 2 | 2.034626 | 0.716236 |
| B0R1D0 | CH211-265 | 26S protea | 13 | 2.10758 | 0.337172 |
| A5PF33 | CH211-51H | F-box-like/ | 4 | 1.446631 | 0.553987 |
| Q6PI45 | CH73-223D | Zgc:55347 | 4 | 1.768332 | 0.485837 |
| Q502E4 | DKEYP-88A | NADH dehy | 8 | 1.433695 | 0.392852 |
| D7RVR6 | EAAT3 | Amino acid | 1 | 1.51728 | 0.533418 |
| Q6TH05 | GABARAPL | GABA(A) re | 2 | 2.553531 | 0.860046 |
| Q6B337 | GIPC | GAIP c-terr | 2 | 1.392486 | 0.455964 |
| Q8AYE2 | HCII | Heparin co | 3 | 1.59823 | 0.351677 |
| Q6P0Z0 | LIIIb | Lamin L3 | 1 | 1.646315 | 0.446324 |
| B5DDE8 | LOC10000( | 40S ribosor | 1 | 1.589346 | 0.471525 |
| A0A8M3AN | LOC10033( | RNA-bindir | 2 | 1.658835 | 0.564277 |
| A0A8M1RD | LOC100331 | myosin-9-li | 1 | 1.510077 | 0.379795 |
| A0A8M9PK | LOC10053 | urokinase p | 1 | 2.365085 | 0.648265 |
| A0A8M9Q | LOC10188 | regulator o | 1 | 1.653906 | 1.75585 |
| A0A8M1P9 | LOC10798 | uncharacte | 1 | 1.737139 | 1.150267 |
| A0A8M6Z5 | LOC10817 | V-type prot | 1 | 1.542381 | 0.410561 |
| A0A8M1RH | LOC10817 | stonustoxir | 1 | 1.424716 | 0.704324 |
| A0A8M9QJ | LOC11044( | RNA-bindir | 8 | 2.035575 | 0.809222 |
| A3KNA6 | LOC56610 | Enhancer o | 2 | 2.448898 | 0.332733 |
| B3DJJ2 | LOC56795 | C-type natr | 1 | 1.314559 | 0.501755 |
| Q1RM69 | MMPLf | Collagenas | 1 | 1.662584 | 0.406425 |
| Q6NWH0 | P5436 | UPF0696 p | 2 | 1.734704 | 0.80074 |
| Q7ZU20 | PA28-gamr | Proteasom | 7 | 2.530361 | 0.470059 |
| Q6TGW1 | PDIP38 | Polymerase | 4 | 2.650087 | 0.648776 |
| B8JIP2 | PRIM2A | DNA prima | 1 | 1.532977 | 1.076624 |
| Q6TGV6 | PSMA5 | Proteasom | 5 | 1.918531 | 0.873438 |
| Q803F7 | Rbm4 | RNA bindin | 16 | 1.61803 | 0.43627 |
| Q7ZWH0 | Rnf30 | Zgc:56376 | 3 | 1.634743 | 1.957823 |
| E7FEB4 | SI:zC146F4 | procollager | 3 | 2.568782 | 0.636756 |
| Q803I0 | SMIF | 5'-(N(7)-me | 2 | 2.625121 | 0.426631 |
| Q7ZVB5 | SNRPD3 | Small nucle | 7 | 1.510437 | 0.333674 |
| A0A097 | Tbeta-a | Thymosin b | 1 | 2.296249 | 0.615749 |
| D3XD60 | Ugt1b1 | UDP-glucur | 1 | 1.640661 | 1.056771 |
| O57332 | Y | Proteasom | 9 | 1.539607 | 0.589091 |
| A0A8M3B6 | aak1b | AP2-associ | 5 | 1.439522 | 0.452955 |
| A0A8M1PZ | abcc1 | ABC-type g | 8 | 2.792762 | 0.535323 |
| A0A8M2BD | abi2b | abl interact | 4 | 1.53854 | 0.66753 |
| Q68EG8 | abra | Zgc:92005 | 3 | 1.69503 | 0.737183 |
| Q4V8X4 | acbd6 | Acyl-CoA-b | 2 | 1.901203 | 0.523634 |
| A0A8M2B9 | accs | 1-aminocyc | 1 | 1.747437 | 0.804911 |
| A0A8M2B6 | acot13 | acyl-coenzy | 3 | 1.637791 | 0.707653 |
| A0A8M1NF | acsl4b | acyl-CoA sy | 5 | 2.201483 | 0.83092 |
| A0A8M9QH | actn4 | alpha-actin | 22 | 1.557072 | 0.492551 |

| | | | | | |
|---|---|---|---|---|---|
| A0A8M3AN | acy1 | N-acyl-alipl | 4 | 3.205058 | 0.415311 |
| A0A8M3AS | add2 | beta-adduc | 2 | 2.282366 | 0.393181 |
| A0A8N7T6 | ahctf1 | protein ELY | 7 | 4.247147 | 0.433224 |
| Q7SY23 | aldh4a1 | Delta-1-pyr | 10 | 1.98596 | 0.870463 |
| Q8JH70 | aldocb | Fructose-bi | 23 | 1.403696 | 0.447116 |
| A0A8M9QS | alpk3a | alpha-prote | 16 | 1.418075 | 0.338506 |
| A0A8M9PZ | alpk3b | titin homol | 2 | 2.208188 | 0.916536 |
| A0A8M6Z0 | ampd1 | AMP deam | 35 | 1.639854 | 0.32489 |
| A9JRX1 | and2 | Actinodin2 | 29 | 1.637922 | 0.738351 |
| A0A8M9QE | ank3b | ankyrin-3 is | 10 | 1.73836 | 0.446176 |
| A0A8M2BJ | ankfy1 | rabankyrin- | 1 | 2.078699 | 0.343381 |
| E9QBZ2 | ankrd28b | Ankyrin rep | 1 | 1.941667 | 0.380353 |
| A0A8M9PH | ano5a_1 | Anoctamin | 4 | 1.83981 | 0.357526 |
| A0A8N1Z3I | anp32b | acidic leuci | 5 | 2.15063 | 1.505393 |
| Q6NUW5 | anp32e | Acidic leuci | 10 | 2.110233 | 0.699894 |
| A4FUN4 | anpepb | Aminopept | 23 | 2.018627 | 0.368584 |
| Q804H2 | anx1a | Annexin | 11 | 1.358174 | 0.585678 |
| A0A8M2BC | ap1b1_1 | AP comple: | 1 | 1.495486 | 0.639156 |
| A0A8M2B2 | apba1b | amyloid be | 1 | 1.737876 | 0.829784 |
| O42363 | apoa1 | Apolipopro | 6 | 2.097033 | 0.570109 |
| A0A8M9NZ | apoa4b.2 | apolipopro | 6 | 1.778191 | 1.043092 |
| A0A8M1P6 | apoc1 | apolipopro | 1 | 1.970802 | 0.851132 |
| Q7T3B8 | ard1a | N-terminal | 5 | 1.594195 | 0.364104 |
| A0A8M2B5 | arfgap2 | ADP-ribosy | 8 | 1.810747 | 0.510016 |
| A0A8M3AH | arhgef12a | rho guanin | 3 | 1.434029 | 0.999056 |
| A0A8M9PR | arhgef1a | Rho guanin | 2 | 2.81057 | 0.537942 |
| A0A8M6Z5 | arhgef7b | rho guanin | 1 | 1.728167 | 0.468898 |
| A0A8M3B3 | arid3a_1 | AT-rich inte | 1 | 2.35768 | 1.352365 |
| A0A8M3AV | armc10 | armadillo r | 1 | 1.884904 | 2.321186 |
| A0A8M9PII | arsb | arylsulfatas | 1 | 1.613093 | 1.356078 |
| Q6DG88 | atg4b | Cysteine pr | 3 | 2.058019 | 0.614129 |
| A0A8M1P2 | atg9b | Autophagy- | 1 | 1.901594 | 0.50782 |
| A0A8M1P5 | atp5f1c | ATP syntha | 7 | 1.453357 | 0.34278 |
| Q6PC77 | atp5h | ATP syntha | 13 | 1.676294 | 0.806426 |
| Q7ZTY2 | atrogin1 | F-box only | 3 | 2.34415 | 1.079748 |
| Q6P026 | banf1 | Barrier-to-a | 4 | 1.390908 | 0.493861 |
| Q502P7 | banp | Protein BAI | 2 | 1.355479 | 0.483131 |
| Q7SZF4 | bat3 | BCL2-assoc | 6 | 1.455732 | 0.549112 |
| A0A8N7TEI | baz1a | bromodom | 3 | 2.388085 | 0.392988 |
| A0A8M2B1 | bicd2 | protein bic | 1 | 1.518818 | 0.32836 |
| A0A8M3AT | bin1b | bridging int | 14 | 1.734455 | 0.564709 |
| A0A8M1NZ | bphl | valacyclovi | 1 | 1.535003 | 1.40877 |
| A0A8M3B5 | brcc3 | Lys-63-spe | 1 | 2.033887 | 0.819622 |
| A0A8M2BC | brd3b | bromodom | 6 | 1.522658 | 0.862897 |
| A0A8M2BH | bub3 | mitotic che | 5 | 1.324495 | 0.391835 |
| Q6IQA1 | c11orf15 | TMEM9 do | 1 | 1.859839 | 0.596405 |
| A0A8M1N5 | c1qbp | Compleme | 8 | 1.834545 | 0.8801 |

| | | | | | |
|---|---|---|---|---|---|
| A0A8M1P7 | c3a.1 | complemer | 29 | 1.369286 | 0.350983 |
| A0A8M3B7 | cadm3 | cell adhesio | 2 | 1.482682 | 0.689675 |
| A0A8M2BL | capn15 | calpain-15 | 1 | 1.910489 | 0.324349 |
| A0A8M9PN | caprin1b | caprin-1 iso | 5 | 3.624458 | 0.890352 |
| A0A8N7US | cars2 | cysteine--tl | 1 | 1.520159 | 0.914619 |
| Q98UI8 | casp3 | Caspase 3, | 5 | 2.011155 | 0.41712 |
| Q8AWD9 | catD | Cathepsin I | 14 | 2.208651 | 0.543825 |
| Q6IQJ1 | cb381 | Mitochond | 2 | 2.069262 | 0.407261 |
| Q6TLF9 | cb430 | AP-3 comp | 5 | 2.746288 | 0.329561 |
| Q8UWM4 | cb543 | Hexosyltra | 1 | 1.367256 | 0.469997 |
| Q7ZUY4 | cb638 | Nucleobinc | 4 | 1.57167 | 0.759141 |
| F1QI36 | cb896spalt | Spalt-like tr | 1 | 1.472035 | 0.783124 |
| Q7ZUR6 | cb985 | Muscle-spe | 1 | 2.861174 | 1.356409 |
| Q6NWY1 | cb997 | 55 kDa eryt | 10 | 1.62036 | 0.440962 |
| Q5RHZ2 | ccdc167 | Coiled-coil | 2 | 1.78911 | 0.849788 |
| A0A8M2B5 | ccdc28a | coiled-coil | 1 | 1.864291 | 1.017481 |
| Q6DRP4 | ccm2 | Cerebral ca | 2 | 1.702661 | 0.871931 |
| A0A8M2B2 | cct8 | T-complex | 20 | 1.320587 | 0.324038 |
| A0A8M1P0 | cdc73 | parafibrom | 7 | 2.439453 | 0.368625 |
| Q6IQA6 | cdkrap3 | CDK5 regul | 4 | 2.213571 | 0.544785 |
| Q6DGV1 | celf4 | CUGBP Elav | 2 | 1.968639 | 1.257716 |
| Q6PBS8 | chchd2l | Chchd2l pro | 2 | 2.289957 | 0.493983 |
| A0A8M2BI | chchd3b | uncharacte | 1 | 2.510239 | 2.291845 |
| Q7ZVB1 | chmp1b | Charged m | 1 | 1.526059 | 1.127304 |
| Q4G3H4 | chuk | Inhibitor of | 2 | 1.895549 | 0.531717 |
| A0A8N7TC | cilp2 | cartilage in | 2 | 1.941592 | 0.632019 |
| A0A8M3AS | clip1a | CAP-Gly do | 9 | 2.844476 | 0.652849 |
| A0A8M3AT | clpb | caseinolytic | 5 | 1.65458 | 0.532803 |
| A0A8M1RE | clpxb | ATP-depen | 6 | 2.969404 | 0.402894 |
| A0A8M9Q7 | cnn2 | Calponin | 5 | 2.727172 | 0.486721 |
| Q7T303 | cnn3 | Calponin | 5 | 1.37127 | 0.919433 |
| A4QP78 | cnot11 | CCR4-NOT | 2 | 2.244035 | 0.425107 |
| A0A8M2BA | cnot2 | CCR4-NOT | 5 | 1.513198 | 0.369472 |
| E7F568 | cobl | Protein cor | 4 | 2.034794 | 1.017224 |
| Q29RB1 | cog4 | Conserved | 3 | 1.762707 | 0.549665 |
| Q6P2U9 | cops3 | COP9 signa | 3 | 1.584592 | 0.772903 |
| Q6P0H6 | cops4 | COP9 signa | 3 | 1.582582 | 0.402671 |
| Q6PC30 | cops5 | COP9 signa | 7 | 1.463653 | 0.444961 |
| F1RAX8 | coq6 | Ubiquinone | 4 | 1.711209 | 1.012003 |
| A0A8M1P2 | cox6c | Cytochrom | 6 | 1.345586 | 0.504149 |
| A0A8N7T7 | csnk2a2b | casein kina | 2 | 1.742199 | 0.400509 |
| A0A8M1N9 | cspg5b | chondroitir | 1 | 1.486066 | 0.765667 |
| A0A8M3AH | cstf2 | cleavage st | 5 | 1.782321 | 0.479323 |
| F1QKH1 | ctr-1 | Copper tra | 4 | 1.45578 | 0.824591 |
| Q6DI42 | cutl1 | Protein CAS | 1 | 1.76866 | 1.024484 |
| Q7ZW86 | cwc27 | Spliceosom | 3 | 1.76064 | 0.577708 |
| Q5RGJ5 | cwf19l1 | CWF19-like | 2 | 1.579083 | 1.2532 |

| | | | | | |
|---|---|---|---|---|---|
| A0A8M1P9 | dab2 | DAB adapto | 2 | 1.45504 | 1.226682 |
| A0A8M2BF | dcaf6 | DDB1- and | 2 | 3.030207 | 0.520748 |
| Q7T3H1 | dctn2 | Dynactin su | 8 | 1.647123 | 1.161715 |
| A0A8M9Q( | ddhd2 | phospholip | 1 | 1.624359 | 0.462703 |
| A0A8M9Q( | ddi2 | protein DD | 11 | 2.141291 | 0.740348 |
| A0A8M1NH | ddx21 | RNA helicas | 3 | 2.333453 | 0.421068 |
| A0A8M1P8 | ddx39aa | RNA helicas | 7 | 1.613145 | 0.512666 |
| A0A8M1N8 | ddx52 | RNA helicas | 4 | 1.733744 | 0.430182 |
| A0A8M1PK | ddx5 | RNA helicas | 11 | 1.362851 | 0.503687 |
| A0A8M2B5 | dennd4a | C-myc pror | 1 | 2.429893 | 0.728807 |
| A0A8M1PA | desma | desmin isof | 27 | 2.293547 | 0.796856 |
| Q7T3E1 | diablo | Direct IAP-I | 1 | 2.19096 | 0.553964 |
| A0A8M9QI | dmd_1 | dystrophin | 1 | 1.627037 | 0.567986 |
| A0A8M9PC | dmgdh | dimethylgly | 15 | 2.018258 | 0.492471 |
| Q7ZVS0 | dnaja2l | DnaJ (Hsp4 | 6 | 1.709964 | 0.534701 |
| A0A8M9PN | dnaja | dnaJ homo | 6 | 1.782866 | 0.529398 |
| A0A8M9QI | dnajc16l | DnaJ homo | 1 | 1.449928 | 0.365062 |
| A0A8M3B8 | dnajc7 | dnaJ homo | 4 | 2.511803 | 0.535024 |
| A0A8M1N5 | dync1li2 | Dynein ligh | 4 | 1.756989 | 1.103746 |
| A0A8M2B7 | eef1db | elongation | 14 | 1.358219 | 0.442763 |
| A0A8N7UV | eef1e1 | eukaryotic | 2 | 2.797977 | 0.485059 |
| A0A8M9QI | ehbp1l1a | uncharacte | 16 | 2.411591 | 0.767286 |
| A0A8M9Q( | ehbp1l1b | EH domain | 11 | 1.517752 | 0.343305 |
| Q66HW2 | ehd1 | EH domain | 14 | 1.32517 | 0.472728 |
| Q7ZUI8 | eif1b | Eukaryotic | 6 | 1.740028 | 0.381504 |
| A0A8M1NX | eif3f | Eukaryotic | 6 | 2.116542 | 0.853096 |
| Q7T3B0 | eif3m | Eukaryotic | 9 | 1.449225 | 0.334382 |
| Q6ZM19 | eif6 | Eukaryotic | 3 | 1.919763 | 0.520514 |
| A0A8M2BII | elavl1 | ELAV-like p | 11 | 2.14408 | 0.715059 |
| A0A8N7UZ | elavl1b | ELAV-like p | 12 | 1.374465 | 0.460666 |
| A0A8M3AS | elmo1 | engulfmen | 1 | 2.348262 | 0.811831 |
| Q6GMI7 | enoph1 | Enolase-ph | 3 | 1.80653 | 1.344392 |
| A0A8M1RH | eps15 | epidermal g | 15 | 1.503238 | 0.532397 |
| A0A8M9P2 | eps15l1b | epidermal g | 4 | 1.53835 | 0.452901 |
| Q7T2D4 | ergic2 | Endoplasm | 1 | 1.885828 | 1.413288 |
| A0A8M1N( | esyt3 | extended s | 2 | 1.4979 | 0.616969 |
| A0A8M3AT | evi5b | EVI5-like pr | 3 | 1.666048 | 0.503122 |
| A0A8M9P2 | exoc1 | exocyst cor | 4 | 2.133576 | 0.752221 |
| A0A8M1PF | exosc10 | exosome co | 6 | 2.593176 | 0.381756 |
| A0A8M1NF | exosc1 | exosome co | 3 | 1.878165 | 0.641719 |
| Q6DRL1 | fa19d11 | RNA helicas | 8 | 2.224835 | 0.479361 |
| A0A8M1QL | fadd | FAS-associa | 4 | 1.383442 | 0.684625 |
| A0A8M1PN | faf1 | FAS-associa | 6 | 3.993887 | 0.475621 |
| A0A8M2B6 | fam13a | protein FAI | 1 | 1.437692 | 0.386276 |
| Q568K9 | fam50a | Protein FAI | 5 | 1.809003 | 0.367032 |
| Q6TEP1 | fam91a1 | Protein FAI | 1 | 1.35112 | 1.055261 |
| A0A8M9PN | far1 | Fatty acyl-C | 1 | 1.762446 | 1.835247 |

| | | | | | |
|---|---|---|---|---|---|
| Q7ZVF2 | fb11d08 | Glutamine | 4 | 3.562686 | 0.877206 |
| A8WFZ2 | fb13d02 | Zgc:172056 | 5 | 1.757678 | 0.559582 |
| Q6PUR8 | fb15c09 | Eukaryotic | 1 | 1.653241 | 0.550076 |
| Q802Z0 | fb49c02 | YTH N(6)-m | 3 | 1.767538 | 0.479337 |
| Q7ZU67 | fb54c08 | RNA helicas | 6 | 1.598451 | 0.53986 |
| Q568R2 | fb57e05 | ATG16 auto | 1 | 1.402988 | 0.554437 |
| Q7ZVG7 | fb60h05 | Fgg protein | 15 | 2.545193 | 0.569674 |
| Q66IB0 | fb75b09 | Protein lin- | 5 | 2.534566 | 0.483149 |
| Q9PW77 | fb77e11 | Pbx4 home | 2 | 1.96506 | 0.950383 |
| Q6PBJ7 | fbxl20 | F-box and l | 1 | 1.43595 | 1.114105 |
| F1QFK9 | fc22d10 | Cleavage st | 6 | 1.923612 | 0.591995 |
| Q803L7 | fc38a11 | Ras homolo | 2 | 1.464983 | 0.506011 |
| Q6IQD9 | fc67a03 | EF-hand ca | 1 | 2.036405 | 0.322267 |
| Q803M4 | fc79h10 | Zgc:55512 | 1 | 1.31725 | 0.488761 |
| E7FBF7 | fcho1 | F-BAR dom | 2 | 2.200494 | 0.419268 |
| Q6DEM2 | fi19g05 | V-crk avian | 9 | 1.457956 | 0.358769 |
| Q5XJJ3 | fj08b08 | ATP syntha | 15 | 1.484302 | 0.865791 |
| Q6PE34 | fj33a08 | Tubulin bet | 2 | 1.493372 | 1.63915 |
| F1QHG7 | fj33c07 | Inhibitor of | 2 | 1.567553 | 0.898998 |
| Q8AWF1 | fj84d04 | Tyrosine-pr | 1 | 1.483001 | 0.367698 |
| Q1JPR9 | fj86f05 | Rho GTPase | 3 | 1.497413 | 1.173307 |
| Q98870 | fk53b11 | Myocyte er | 2 | 1.416269 | 0.462677 |
| A0A8M1RN | flnca | filamin-C is | 36 | 2.651614 | 0.655722 |
| A8WG78 | foxred1 | FAD-depen | 3 | 1.962461 | 0.599006 |
| A0A8M1N6 | fpgt | fucose-1-pl | 1 | 1.922215 | 1.151939 |
| A0A8M1RL | fyco1b | FYVE and c | 2 | 1.776709 | 0.738221 |
| A0A8M3B9 | gak | cyclin-G-as | 3 | 2.149663 | 0.680269 |
| Q5MJ86 | gapdh-2 | Glyceraldeh | 21 | 1.520713 | 0.396297 |
| A0A8M2BE | gcc1 | GRIP and co | 1 | 2.374411 | 0.696373 |
| Q6P3I8 | gcdh | Glutaryl-Co | 6 | 1.620286 | 0.323736 |
| A0A8M1N1 | gga1 | ADP-ribosy | 4 | 1.415154 | 0.61809 |
| A0A8M6Z9 | ggps1 | geranylgera | 3 | 2.082444 | 0.395289 |
| O93430 | glra1 | Glycine rec | 5 | 1.646111 | 0.925984 |
| Q7SXP8 | gmppab | Mannose-1 | 2 | 1.781936 | 0.441408 |
| O42248 | gnb2l1 | Guanine nu | 28 | 2.444969 | 0.447618 |
| A0A8N7TC | gnpda2 | Glucosamir | 4 | 1.314647 | 0.76455 |
| A0A8M3AS | golga2 | golgin subf | 2 | 2.094421 | 0.593266 |
| A0A8M2BC | golgb1 | golgin subf | 6 | 2.250469 | 0.536356 |
| Q7SYK7 | got2a | Aspartate a | 16 | 1.708631 | 0.367645 |
| Q5CZM4 | gpb6bb | Glycoprote | 1 | 1.784791 | 0.622361 |
| A0A8M1N3 | gpx1a | Glutathione | 5 | 1.614178 | 0.511162 |
| A0A8M1N2 | gpx1b | Glutathione | 4 | 1.75883 | 0.610834 |
| A0A8M1N7 | gpx4b | Glutathione | 9 | 1.32641 | 0.43282 |
| A0A8N1Z0 | gripap1 | GRIP1-asso | 9 | 1.675963 | 0.455566 |
| Q6DGU9 | gstk1 | Glutathione | 7 | 1.549467 | 0.335245 |
| Q6IQP1 | gygl | Glycogenin | 2 | 1.625891 | 1.325398 |
| Q71PD7 | h2az2a | Histone H2 | 15 | 1.678936 | 0.68241 |

| | | | | | |
|---|---|---|---|---|---|
| Q6DI22 | hadhsc | Hydroxyacy | 9 | 1.575355 | 0.420647 |
| Q5BJJ5 | hdhd2 | Haloacid de | 3 | 1.421671 | 0.334581 |
| Q32PU2 | hm:zehn14 | Calponin | 3 | 1.40406 | 0.863304 |
| A0A8M2B3 | hmbsb | hydroxyme | 3 | 1.322818 | 0.542379 |
| A0A8M1PH | hnrnpa0a | heterogene | 11 | 1.718625 | 0.785423 |
| Q7ZU48 | hnrpa0l | Hnrpa0l pro | 10 | 1.934242 | 1.22342 |
| A0A8M1NC | homer1b | homer prot | 1 | 2.160576 | 0.967344 |
| A0A8M1P2 | hpgd | 15-hydroxy | 1 | 1.355575 | 0.328745 |
| Q6P5L8 | hsdl2 | Hydroxyste | 15 | 1.426528 | 0.377112 |
| Q6PH56 | hsp70 | Heat shock | 3 | 2.488216 | 1.373138 |
| Q90474 | hsp90a.1 | Heat shock | 28 | 1.388604 | 0.470429 |
| Q5RG12 | hsp90a2 | Heat shock | 3 | 1.365463 | 0.630291 |
| Q0R4G9 | hspa12b | HSPA12B | 2 | 1.524668 | 0.903845 |
| A0A8M1NE | hspa1b | uncharacte | 1 | 1.419514 | 1.194781 |
| A5JV83 | hspb11 | Heat shock | 1 | 2.064621 | 1.552662 |
| E7F6A5 | id:ibd1090 | 39S ribosor | 3 | 1.973955 | 0.438546 |
| A0A8N7T7I | ifi35 | interferon- | 2 | 1.784311 | 1.26333 |
| D2JGP1 | ifngr1 | Cytokine re | 1 | 1.786567 | 0.393843 |
| A0A8M1N9 | ighmbp2 | DNA-bindir | 4 | 1.37881 | 0.538958 |
| A0A8M9QC | il16 | pro-interleu | 1 | 2.227772 | 0.407917 |
| Q1JPY6 | im:689555 | Prefoldin su | 4 | 1.408113 | 0.823536 |
| Q6YLH7 | im:690290 | Caveolin | 2 | 1.635526 | 1.451066 |
| E7FFC4 | im:690405 | Apoptosis-i | 4 | 2.023149 | 0.440859 |
| Q4V9A4 | im:691233 | exo-alpha-s | 4 | 1.332815 | 0.701066 |
| Q58EC1 | im:697126 | Snx4 prote | 8 | 1.369859 | 0.446194 |
| B3DID0 | im:714806 | FAS-associa | 4 | 1.874773 | 0.536195 |
| A0A8N7TB | ints5 | integrator c | 3 | 1.421931 | 0.373864 |
| A0A8M1P1 | itpkcb | Kinase | 4 | 1.825893 | 0.450964 |
| Q7ZVQ8 | ivns1abpb | Influenza v | 4 | 2.982165 | 0.35047 |
| A0A0R4IXF | kat2a | Histone ace | 2 | 1.807924 | 0.529756 |
| A5PLC1 | kdelc2 | KDEL motif | 7 | 1.4179 | 0.415877 |
| A0A8M1NY | kdm1a | Lysine-spec | 7 | 1.929401 | 0.4346 |
| A0A8M1QH | kdm4aa | [histone H3 | 4 | 1.953089 | 0.690546 |
| A0A8M6Z4 | kdm5a | [histone H3 | 3 | 1.442098 | 0.36313 |
| A0A8M3AS | klhl43 | kelch-like p | 2 | 1.605106 | 1.634897 |
| A0A8M3AV | kpnb3 | importin-5 | 13 | 1.71384 | 0.466795 |
| A0A8M1P1 | ktn1 | kinectin iso | 24 | 3.116224 | 0.508411 |
| Q5BKX3 | lamp2 | Lysosomal | 1 | 1.326433 | 0.610008 |
| A0A8M1P7 | lcat | phosphatid | 4 | 1.341902 | 0.905296 |
| Q9PVK4 | ldhba | L-lactate de | 13 | 1.602699 | 0.515337 |
| Q24JV9 | lgals3bpa | Galectin-3- | 1 | 1.313764 | 0.980678 |
| Q6NY73 | lgals3bpb | Galectin-3- | 1 | 2.291102 | 1.301916 |
| A0A8M1RJ | lgals8a | Galectin | 2 | 2.209702 | 1.099011 |
| A0A8M9PH | limch1a | LIM and ca | 28 | 1.836233 | 0.380485 |
| Q3B748 | llph | Protein LLP | 1 | 2.918537 | 0.451784 |
| A0A8M3B2 | lmo7a_1 | LIM domain | 2 | 2.326936 | 0.569914 |
| A0A8M1P2 | lmtk2 | serine/thre | 3 | 1.330702 | 0.323736 |

| | | | | | |
|---|---|---|---|---|---|
| A0A8M2B5 | lpin1 | phosphatid | 4 | 1.48323 | 1.393159 |
| A0A8M2BC | lrrfip2 | leucine-rich | 6 | 2.649239 | 0.405466 |
| A0A8M1NF | lrsam1 | E3 ubiquiti | 7 | 1.890189 | 0.392814 |
| A0A8M2B5 | lsm14ab | protein LSN | 5 | 2.272353 | 0.333569 |
| A0A8M1PD | lyar | cell growth | 2 | 2.077492 | 1.610946 |
| Q90YS5 | lys-C | Lysozyme C | 7 | 1.694747 | 0.379586 |
| Q8UUZ1 | mab21l2 | Protein ma | 1 | 1.94702 | 0.841376 |
| A0A8M2BH | man2b1 | Alpha-man | 7 | 1.304297 | 0.448182 |
| Q9DGE2 | mapk14a | Mitogen-ac | 4 | 2.803725 | 0.4036 |
| A0A8M9PX | mark3a | non-specifi | 1 | 1.774005 | 0.696978 |
| P52161 | max | Protein ma | 2 | 1.776291 | 0.66536 |
| Q6P0E3 | mcrs1 | Microspher | 2 | 1.872621 | 0.58068 |
| Q7ZVN7 | med15 | Mediator o | 1 | 1.907943 | 0.769508 |
| Q6PFL0 | med27 | Mediator o | 1 | 1.421048 | 0.799046 |
| A0A8M3AK | meis2a | homeobox | 1 | 1.402052 | 0.632977 |
| Q803S3 | memo1 | Protein ME | 2 | 2.081093 | 0.698638 |
| Q7ZVJ8 | mettl26 | Methyltran | 1 | 1.695212 | 0.734094 |
| Q804S5 | mib1 | E3 ubiquiti | 2 | 1.927078 | 1.343481 |
| F1RA39 | mical2b | [F-actin]-m | 1 | 4.124483 | 0.617381 |
| A0A8M1NE | mief2 | mitochond | 1 | 1.571445 | 0.351056 |
| A0A8M1P6 | mmp13a | matrix met | 1 | 2.831495 | 2.30746 |
| Q789K9 | mp:zf637-2 | Ubiquitin-c | 2 | 1.409717 | 0.793225 |
| A0A8M2BH | mpp6b | MAGUK p5 | 5 | 1.524199 | 0.595552 |
| Q502J9 | mrpl30 | 39S ribosor | 1 | 1.970929 | 0.795946 |
| A0A8M1N7 | mrpl44 | 39S ribosor | 2 | 1.929772 | 0.757766 |
| Q503X2 | mrpl4 | 39S ribosor | 2 | 1.489196 | 0.964942 |
| A5PLF9 | mrps22 | 28S ribosor | 5 | 1.623421 | 0.669342 |
| Q498Z6 | mrps7 | 28S ribosor | 4 | 1.321927 | 0.87299 |
| A0A8M3B2 | msh3 | DNA misma | 1 | 2.287266 | 0.910838 |
| A0A8M3AI | msi2b | RNA-bindir | 3 | 1.836535 | 1.351808 |
| A0A8M2BK | mta2 | metastasis- | 3 | 2.848949 | 0.769015 |
| A0A8M1NX | mustn1b | Musculoske | 1 | 1.32755 | 1.505364 |
| Q6P3L0 | mvp | Major vault | 25 | 1.461805 | 0.362568 |
| Q6DKF0 | mxc | Interferon- | 1 | 2.060857 | 0.583592 |
| A0A8M1P8 | myh7ba | myosin, he | 9 | 1.502502 | 0.460469 |
| Q6NVA6 | myl9 | Myosin, lig | 4 | 2.258752 | 0.449551 |
| A0A8M1RF | myoz3a | myozenin-2 | 2 | 2.78585 | 0.924982 |
| A0A8N7UT | mypn | myopalladi | 9 | 2.281609 | 0.428492 |
| A0A8M1NL | naa30 | N-alpha-ac | 1 | 1.648802 | 0.337026 |
| Q6DBY2 | naa50 | N-alpha-ac | 8 | 1.853594 | 0.837346 |
| A1L1N2 | naf1 | TNFAIP3-in | 1 | 1.91934 | 0.457094 |
| A0A8M2BJ | nap1l1 | nucleosom | 7 | 1.350164 | 0.372044 |
| Q98881 | nar | Cleavage ar | 3 | 1.319379 | 0.793535 |
| Q5XLR3 | ncstn | Nicastrin | 3 | 1.998144 | 0.326077 |
| Q6PFP6 | ndor1 | NADPH-dep | 3 | 1.794612 | 0.454575 |
| A0A8M1P3 | necab2 | N-terminal | 4 | 1.312181 | 0.577131 |
| Q0II00 | nfkb2 | Nuclear fac | 3 | 1.748171 | 0.344721 |

| | | | | | |
|---|---|---|---|---|---|
| Q6PFJ1 | ngdn | Neuroguidi | 1 | 1.680685 | 0.562359 |
| Q7ZV23 | nhp2l1 | Ribonuclop | 1 | 1.616211 | 0.785987 |
| Q568R1 | nsrp1 | Nuclear spe | 1 | 1.428459 | 0.492794 |
| A0A8M9Q( | ntm | neurotrimi | 3 | 1.648022 | 0.399305 |
| A0A8M2BH | nub1 | NEDD8 ulti | 3 | 2.051528 | 0.599227 |
| Q3B7Q7 | nubp2 | Cytosolic F | 1 | 1.749355 | 1.836527 |
| A0A8M2BE | nucb1 | Nucleobinc | 4 | 3.067294 | 0.441201 |
| F1QNV4 | nup133 | Nuclear po | 10 | 1.66916 | 0.568204 |
| A0A8M9Q! | obsl1b | obscurin-lik | 3 | 1.9544 | 0.465762 |
| Q5U3P2 | ocma | Parvalbumi | 2 | 1.517569 | 0.356422 |
| Q29RB4 | olfml3a | Olfactomed | 5 | 1.689879 | 0.69791 |
| Q5RI56 | optn | Optineurin | 3 | 2.000242 | 0.701519 |
| Q4V920 | pacsin1b | Protein kin | 1 | 1.653183 | 0.673828 |
| A0A8M9PT | palld | palladin iso | 10 | 2.35732 | 0.84439 |
| A0A8M2BA | palm1b | uncharacte | 4 | 1.692789 | 0.500185 |
| A0A8M1Q1 | pargl | poly(ADP-r | 1 | 1.334902 | 1.357304 |
| A0A8M2BC | pbrm1l | polybromo | 4 | 1.548404 | 0.395562 |
| A0A8M9PZ | pck1 | phosphoen | 7 | 1.534852 | 0.62389 |
| A0A8M2B9 | pde7a | Phosphodie | 1 | 1.889332 | 0.594051 |
| A0A8M9QH | pdgfrb | receptor pr | 2 | 1.710412 | 0.36143 |
| A0A8M1NX | pdia3 | Protein disu | 24 | 2.554696 | 0.32603 |
| A0A8M3AV | pdlim5b | PDZ and LII | 3 | 1.877104 | 0.675738 |
| Q66HY8 | pdxdc1 | Pyridoxal-d | 2 | 1.726751 | 0.427083 |
| B8A4Q2 | pdxk | Pyridoxal k | 2 | 1.398389 | 0.892661 |
| B3DI21 | pdxp | Pyridoxal (p | 1 | 1.878826 | 0.49948 |
| A0A8M1RL | pfdn4 | Prefoldin su | 2 | 1.321246 | 0.46808 |
| A0A8M2BD | pfkfb4b | 6-phosphof | 3 | 3.139817 | 1.188827 |
| Q502L2 | pgam5 | Serine/thre | 5 | 2.191927 | 0.444917 |
| A0A8M9QI | phka2 | Phosphoryl | 2 | 2.404397 | 0.520166 |
| A0A8M2BD | phkb | Phosphoryl | 11 | 2.119663 | 0.340638 |
| A0A8M2BD | phkb_1 | Phosphoryl | 2 | 1.481444 | 0.369924 |
| Q6PE18 | pi4k2a | Phosphatid | 4 | 2.127667 | 0.409761 |
| A0A8M9PN | pik3r4 | non-specifi | 3 | 1.345414 | 0.677311 |
| Q6IQE1 | pip4k2c | Phosphatid | 2 | 1.964454 | 0.533555 |
| A0A8M9P8 | pld2 | Phospholip | 1 | 2.018448 | 0.585247 |
| B3DLH5 | plek2 | Pleckstrin-2 | 1 | 2.700139 | 0.979937 |
| A0A8M1QH | polr1a | DNA-direct | 10 | 2.128634 | 0.510427 |
| Q6DEG5 | polr2e | DNA-direct | 5 | 2.380524 | 0.493849 |
| A5D6S6 | polr2l | DNA-direct | 3 | 1.802017 | 0.897563 |
| Q642H5 | ppp2r1b | Protein pho | 16 | 2.099172 | 0.335613 |
| A2CF49 | ppp2r4 | Serine/thre | 3 | 1.351241 | 0.337781 |
| Q6P964 | ppp4r2a | Serine/thre | 3 | 1.502581 | 0.337374 |
| A0A8M3AV | ppp6r3 | serine/thre | 10 | 3.283002 | 0.540583 |
| Q6NWG4 | prmt6 | Protein arg | 3 | 2.577278 | 0.33556 |
| A0A8M1RC | proser2 | proline anc | 2 | 1.591853 | 0.770509 |
| M5BFV8 | prp | Collagen ty | 7 | 1.966407 | 0.518855 |
| Q1JPZ7 | prpf39 | Pre-mRNA- | 11 | 1.939766 | 0.408665 |

| | | | | | |
|---|---|---|---|---|---|
| Q5XJA3 | prrc1 | Protein PRI | 4 | 1.793813 | 0.42355 |
| A0A8M1NS | prune | exopolypho | 2 | 1.765846 | 0.807728 |
| A0A8M2BC | psmc3 | 26S protea | 24 | 1.661243 | 0.748769 |
| F6P3G4 | psmd11a | 26S protea | 8 | 1.457844 | 0.701517 |
| F1QGH9 | psmd11b | 26S protea | 17 | 2.260414 | 0.767158 |
| A0A8M1N5 | psmd5 | 26S protea | 3 | 1.665168 | 1.008464 |
| F1QFR9 | psme4a | Proteasom | 10 | 2.135835 | 0.382349 |
| A0A8M1P3 | ptbp2a | uncharacte | 1 | 1.487556 | 0.432201 |
| Q8QGV5 | ptgds | Lipocalin-ty | 1 | 1.37821 | 1.737466 |
| A0A8M1P7 | ptpn1 | Tyrosine-pr | 8 | 1.554546 | 0.61847 |
| A0A8M2B5 | ptx3a | pentraxin-r | 9 | 1.521625 | 0.824355 |
| Q6IQE0 | puf60b | Poly(U)-bin | 9 | 1.874376 | 0.379513 |
| Q6P971 | pycr1 | Pyrroline-5 | 1 | 2.153187 | 0.435098 |
| Q5SPD7 | pycr3 | Pyrroline-5 | 3 | 1.568646 | 1.003304 |
| Q7T2C6 | rab7 | RAB7, mem | 10 | 1.744815 | 0.581627 |
| A0A8M2BJ | rabep1 | rab GTPase | 2 | 2.314645 | 0.725333 |
| Q32PS7 | rad23ab | UV excision | 7 | 1.380655 | 0.436306 |
| Q7ZWF0 | rae1 | mRNA expo | 4 | 1.625923 | 0.819876 |
| A0A8M1NV | raf1b | non-specifi | 1 | 2.177527 | 0.720576 |
| Q1LUS8 | ranbp10 | Ran-binding | 2 | 2.327994 | 0.56042 |
| A1L252 | ranbp9 | Ran-binding | 2 | 2.143425 | 0.871763 |
| A9JTA8 | rangap1 | Ran GTPase | 5 | 2.109381 | 0.65327 |
| B1H1M1 | rap1gap | Zgc:175180 | 3 | 1.67285 | 0.555719 |
| A0A8M3B3 | rasa3 | ras GTPase | 1 | 1.52593 | 0.482283 |
| Q6P3H7 | rbbp4 | Histone-bir | 2 | 2.45368 | 0.617015 |
| Q7ZTY4 | rbbp7 | Histone-bir | 9 | 2.06425 | 0.505234 |
| A0A8M9Q/ | rbfox2 | RNA bindin | 1 | 2.058503 | 0.963097 |
| A0A8M1P3 | rfk | riboflavin k | 1 | 1.842063 | 0.333724 |
| D7PS89 | rfl | FRAS1-rela | 3 | 1.37582 | 0.346114 |
| A0A8M9QE | rictora | rapamycin- | 2 | 1.391295 | 0.796624 |
| A7E2K4 | rplp2l | 60S acidic r | 6 | 1.712794 | 0.583403 |
| A0A8M2B9 | rprd1b | regulation | 8 | 2.980347 | 0.336373 |
| A0A8M9PN | rrad | GTP-bindin | 1 | 1.429573 | 0.582153 |
| Q1LXR3 | rrbp1 | Ribosome-l | 2 | 2.230474 | 1.04678 |
| P79732 | rrm1 | Ribonucleo | 4 | 1.538212 | 0.645469 |
| Q4G5V0 | rtn6 | Reticulon | 7 | 1.339219 | 0.530857 |
| A0A8M2B6 | s100s | Protein S10 | 1 | 2.237084 | 0.83483 |
| Q6JWU7 | sb:cb121 | Coatomer s | 10 | 1.78283 | 0.554597 |
| Q6NYJ7 | sb:cb173 | Legumain | 3 | 2.133275 | 0.797644 |
| Q7ZTH9 | sb:cb425 | Isovaleryl-C | 10 | 1.912722 | 0.651433 |
| F1Q5Z8 | sb:cb621 | Sequestoso | 2 | 2.280298 | 1.817563 |
| Q6NZ32 | sb:cb897 | Flavin-cont | 4 | 1.735767 | 0.727721 |
| A0A8M1RE | scaf8 | protein SCA | 6 | 2.445141 | 0.338176 |
| Q802F3 | selenof | Selenoprot | 3 | 2.38028 | 0.417185 |
| Q1LVN8 | selenoo1 | Protein ade | 1 | 1.393399 | 0.355508 |
| Q7ZW38 | sephs1 | Selenide, w | 3 | 1.793274 | 0.888356 |
| A0A8M2BK | 6-Sep | Septin | 14 | 1.339813 | 0.330148 |

| | | | | | |
|---|---|---|---|---|---|
| Q5SNQ7 | serac1 | Protein SEF | 1 | 1.396825 | 0.517747 |
| A0A8N7V0 | serpina1l | serine (Or c | 23 | 2.641159 | 0.487289 |
| A0A8M2BL | serpinb1l3 | serpin pept | 6 | 2.10077 | 0.684014 |
| E9QGQ0 | setd1a | SET domair | 1 | 1.370719 | 0.452561 |
| Q6DHG0 | setd7 | Histone-lys | 3 | 2.233869 | 0.693985 |
| A0A8N7TA | sf3b1 | splicing fac | 20 | 2.525766 | 0.407316 |
| Q1LVE8 | sf3b3 | Splicing fac | 19 | 1.667914 | 0.363285 |
| Q803T6 | sh3gl2 | SH3 domai | 6 | 1.600708 | 0.552867 |
| A0A8M3AV | sh3glb1a | Endophilin- | 1 | 1.494526 | 0.853731 |
| A0A8M2BE | sh3tc1 | SH3 domai | 1 | 1.550868 | 1.165023 |
| Q5SPK4 | si:busm1-1 | F-box only | 1 | 1.484628 | 1.498709 |
| A0A8M9QJ | si:ch1073-5 | E3 SUMO-p | 13 | 2.190882 | 0.494954 |
| B3DIT4 | si:ch211-15 | valine--tRN | 3 | 1.871373 | 0.653825 |
| E7F569 | si:ch211-15 | ES1 proteir | 4 | 2.45095 | 0.68146 |
| A0A8M9Q4 | si:ch211-16 | clustered n | 8 | 1.45474 | 0.536634 |
| A0A8M9P9 | si:ch211-20 | disabled hc | 2 | 2.09679 | 0.401111 |
| A0A8M9PF | si:ch211-21 | uncharacte | 3 | 1.680683 | 0.434848 |
| A0A8M1N9 | si:ch211-21 | uncharacte | 2 | 1.537519 | 0.363268 |
| A0A8M9Q3 | si:ch211-21 | rho GTPase | 1 | 1.419797 | 1.096518 |
| A0A8M3B4 | si:ch211-24 | myomegali | 2 | 2.734661 | 0.393471 |
| A0A8N7UC | si:ch211-26 | zinc finger | 5 | 2.423262 | 0.343606 |
| X1WDU0 | si:ch211-28 | Si:ch211-28 | 2 | 1.428914 | 1.337693 |
| A0A8M3AY | si:ch211-39 | uncharacte | 2 | 3.05285 | 1.296797 |
| A0A8M2B9 | si:ch73-103 | centrosom | 5 | 2.975831 | 0.534713 |
| A0A8M1P6 | si:ch73-234 | vesicle-ass | 2 | 2.902056 | 0.419017 |
| A0A8M9PT | si:ch73-375 | kinesin-like | 1 | 2.573026 | 0.526155 |
| A0A8M1Q7 | si:ch73-63e | protein stra | 4 | 3.331538 | 0.574408 |
| A0A8M2BI5 | si:ch73-70c | epidermal g | 4 | 1.301919 | 0.536365 |
| Q1LXF9 | si:dkey-102 | Si:dkey-102 | 4 | 2.304238 | 0.68058 |
| A0A8M1PZ | si:dkey-105 | alpha-2-ma | 2 | 2.176552 | 0.754871 |
| A0A8N7TB | si:dkey-10f | transcriptic | 4 | 2.999266 | 0.545787 |
| A4VCF2 | si:dkey-12h | Si:dkey-12h | 3 | 2.227602 | 0.518667 |
| A0A8M6YS | si:dkey-12j | probable a: | 2 | 1.919334 | 1.045966 |
| A5PMM5 | si:dkey-193 | Sorting nex | 1 | 1.975802 | 0.568141 |
| A0A8M2BK | si:dkey-205 | AT-rich inte | 1 | 2.17417 | 1.116433 |
| A0A8M9Q2 | si:dkey-240 | NLR family | 2 | 1.344453 | 0.707038 |
| A0A8M9PN | si:dkey-250 | carcinoemb | 1 | 1.315778 | 0.562069 |
| A7E2I2 | si:dkey-254 | Arf-GAP wi | 3 | 1.505789 | 0.466967 |
| A0A8M1P8 | si:dkey-276 | uncharacte | 14 | 1.422184 | 0.523761 |
| A0A8M9QE | si:dkey-27p | uncharacte | 2 | 2.244058 | 0.902704 |
| A0A8M9Q0 | si:dkey-38l | uncharacte | 4 | 3.187159 | 0.654814 |
| Q08C89 | si:dkey-49c | Lectin, mar | 3 | 1.991355 | 0.509352 |
| A0A140LG1 | si:dkey-56n | Si:dkey-56n | 3 | 2.78611 | 0.962162 |
| Q5RHE5 | si:dkey-90n | Si:dkey-90n | 6 | 1.874351 | 0.696214 |
| A0A8M3AY | si:dkeyp-12 | uncharacte | 7 | 1.729488 | 0.349308 |
| A0A8M2BE | si:dkeyp-69 | uncharacte | 2 | 1.414112 | 1.230549 |
| Q90X40 | si:dz150f13 | DNA polym | 1 | 1.419794 | 0.51209 |

| | | | | | |
|---|---|---|---|---|---|
| F1QGJ3 | si:rp71-61h | Perilipin | 5 | 2.02079 | 0.430599 |
| Q6IQD0 | si:zc14a17. | Proteasom | 14 | 1.621098 | 0.342409 |
| A0A8M3B4 | sin3ab | paired amp | 2 | 1.609885 | 1.156945 |
| A0A8M1PZ | slc22a7b.3 | solute carri | 2 | 1.348255 | 0.899685 |
| Q6P5K6 | slc25a20 | Solute carr | 2 | 1.379827 | 0.832021 |
| Q6DHC3 | slc25a40 | Probable m | 1 | 3.426923 | 0.556105 |
| A0A8M9PN | slc44a2_2 | choline tra | 2 | 1.947673 | 1.000586 |
| A0A8M2BC | slmapa | sarcolemm | 6 | 1.581948 | 0.518625 |
| A0A8M9Q9 | smap1 | stromal me | 5 | 1.807853 | 0.616178 |
| Q5TZ66 | snap25a | Synaptosor | 12 | 1.870902 | 0.420189 |
| A0A8M1NZ | snap29 | synaptosor | 2 | 1.92699 | 0.381034 |
| Q6P2T1 | snrpa | U1 small nu | 4 | 1.546 | 0.449421 |
| Q5RID7 | snx17 | Sorting nex | 1 | 1.580403 | 0.41991 |
| Q566W7 | snx30 | Sorting nex | 1 | 1.870754 | 0.532753 |
| A0A8M6YV | soga3b_1 | uncharacte | 2 | 1.684603 | 0.57069 |
| A0A8M1NC | sort1b | sortilin 1b p | 1 | 1.308059 | 0.803175 |
| A0A8M1QL | soul5 | heme-bind | 4 | 1.665489 | 0.663529 |
| A0A8M1P8 | spag9a | C-Jun-amin | 3 | 2.563981 | 0.418699 |
| Q503Q1 | spata18 | Mitochond | 1 | 1.569269 | 0.494048 |
| A0A8M1N1 | srek1 | splicing reg | 1 | 1.875888 | 0.4235 |
| A0A8M6YZ | srpk1a | SRSF protei | 2 | 2.130568 | 0.427163 |
| A0A8M1NE | stag2a | Cohesin sul | 1 | 1.340972 | 0.6518 |
| A0A8M2BA | stam | signal trans | 4 | 1.327496 | 0.331376 |
| A0A8M2BF | stat4 | Signal trans | 2 | 2.020383 | 0.911517 |
| A0A8M1PJ | stau1 | double-stra | 8 | 2.057421 | 0.563276 |
| Q7ZW47 | stau2 | Double-stra | 10 | 1.486325 | 0.382385 |
| A5PLE0 | stoml3 | Stomatin (E | 1 | 1.51187 | 2.189745 |
| A0A8M9QC | stx16 | syntaxin-16 | 1 | 1.439861 | 0.655847 |
| A0A8M1PD | stx5al | syntaxin-5 | 1 | 2.891218 | 0.68427 |
| Q6DHL4 | sumo2 | Small ubiqu | 2 | 1.798385 | 0.842369 |
| A0A8M1NH | sv2ca | synaptic ve | 2 | 2.565412 | 0.607778 |
| A0A8M1PS | synpo2lb | synaptopoc | 3 | 1.911722 | 0.612897 |
| A0A8M2B3 | synrg | synergin ga | 3 | 2.267817 | 0.456453 |
| Q7SXR1 | tarbp2 | RISC-loadin | 2 | 1.604264 | 0.815689 |
| Q6GML7 | tatdn1 | Deoxyribor | 3 | 1.421224 | 0.342918 |
| A9JSV4 | tbc1d13 | TBC1 doma | 3 | 1.90628 | 0.581675 |
| A0A8M1P7 | tcap | telethonin | 3 | 2.466415 | 1.839381 |
| A0A8M3AU | tcerg1b | transcriptic | 2 | 1.371467 | 0.523684 |
| Q503G3 | tdg | G/T misma | 1 | 1.865051 | 0.54925 |
| B3DJW5 | th | Tyrosine 3- | 1 | 1.530649 | 0.925285 |
| A0A8M2B7 | thbs1b | thrombosp | 9 | 1.564138 | 0.453424 |
| Q6DI06 | timm10 | Mitochond | 1 | 1.76316 | 0.870698 |
| A0A8M1PC | tm9sf4 | Transmeml | 3 | 1.579252 | 0.408397 |
| B3DJK0 | tmem160 | Transmeml | 1 | 1.425736 | 1.192273 |
| A7E2V0 | tmem230b | Transmeml | 1 | 1.591338 | 0.722457 |
| A0A8M2B7 | tnfaip2a | tumor necr | 2 | 1.39894 | 0.363696 |
| A0A8M9PV | tpd52l2b | tumor prot | 4 | 1.841953 | 0.346512 |

| | | | | | |
|---|---|---|---|---|---|
| Q1MTI4 | tpi1a | Triosephos | 7 | 1.390243 | 0.627726 |
| A0A8M2B1 | tpm2 | tropomyos | 17 | 1.367069 | 0.501029 |
| Q7T3F0 | tpm4 | Tropomyos | 5 | 1.897167 | 0.356996 |
| A0A8M1N7 | tprb | Nucleoprot | 19 | 2.664443 | 0.401829 |
| Q6E2N3 | trim33 | E3 ubiquitin | 4 | 1.83953 | 0.336687 |
| A0A8M1PV | trim35-33 | tripartite m | 1 | 1.347876 | 0.658226 |
| Q6DHS0 | trim55 | Tripartite m | 8 | 3.546889 | 1.140479 |
| A0A8M2B9 | trim55b | tripartite m | 2 | 2.598988 | 1.784539 |
| A0A8M1QI | trim65 | tripartite m | 1 | 1.450255 | 0.485187 |
| A0A8M1P0 | trip11 | thyroid rec | 5 | 2.40905 | 1.051808 |
| A0A8M1N3 | trmt10c | tRNA meth | 1 | 1.484578 | 0.474546 |
| B3DHG0 | trub1 | tRNA pseud | 4 | 1.330895 | 0.533547 |
| A0A8M3B5 | tsc22d2 | TSC22 dom | 2 | 1.472761 | 0.344489 |
| A0A8M2BL | tshz2 | teashirt ho | 1 | 1.467207 | 0.488378 |
| A3KPN8 | ttc38 | Tetratricop | 4 | 1.907085 | 0.474099 |
| A0A8M1N7 | tubb2b | Tubulin bet | 59 | 1.892786 | 0.342734 |
| Q6P5M9 | tubb2c | Tubulin bet | 5 | 1.302779 | 1.645459 |
| Q6GMH3 | twf2 | Twinfilin-2 | 5 | 1.781149 | 0.574958 |
| Q5RIN4 | txlnba | Taxilin beta | 11 | 2.231921 | 1.200408 |
| A0A8M2B9 | txndc5 | protein disu | 18 | 1.411234 | 0.395732 |
| A0A8M1PF | txnl1 | thioredoxin | 11 | 2.238577 | 0.938165 |
| A0A8M9Q3 | tyrp1a | 5,6-dihydro | 3 | 1.336995 | 0.429714 |
| A0A8M2BC | uacab | uveal autoa | 9 | 2.842882 | 0.975331 |
| A0A8M6Z3 | ubap2a | ubiquitin-a | 2 | 1.966023 | 0.546268 |
| A0A8M2BE | ubap2b | ubiquitin as | 8 | 2.019681 | 0.815837 |
| Q9W6H5 | ube2ia | SUMO-conj | 3 | 1.452274 | 0.430776 |
| Q6PEH5 | ube2v2 | Ubiquitin-c | 9 | 1.455581 | 0.490139 |
| A0A8M9PK | ube4b | ubiquitin co | 10 | 2.079223 | 0.46551 |
| A0A8M9PJ | ubr5 | E3 ubiquitin | 6 | 2.818268 | 0.378353 |
| A0A8M9PC | uckl1b | Uridine-cyt | 1 | 1.394498 | 0.777158 |
| A0A8M3B8 | uhrf1bp1l | UHRF1-bind | 2 | 1.350194 | 0.389919 |
| A0A8M1NC | uqcrc2a | ubiquinol-c | 24 | 1.454136 | 0.509215 |
| A0A8N7UV | usp10 | Ubiquitin ca | 7 | 2.920177 | 0.602521 |
| F1QFS9 | usp13 | Ubiquitin ca | 5 | 2.354213 | 0.533431 |
| A0A8M1QI | usp28 | ubiquitin ca | 3 | 2.476756 | 0.889915 |
| A0A8M1RI | usp38 | ubiquitin ca | 2 | 1.723803 | 0.448694 |
| A0A8M1RK | usp4 | ubiquitinyl | 5 | 4.211008 | 0.421413 |
| A0A8M9PX | vcana | chondroitin | 11 | 1.423458 | 0.63616 |
| Q7ZU99 | vcp | Transitiona | 33 | 3.140274 | 0.569352 |
| Q7ZW40 | vdp | General ves | 16 | 2.360954 | 0.453489 |
| Q6TNP8 | vps26a | Vacuolar pr | 4 | 1.531654 | 0.492541 |
| A0A8M2B7 | vta1 | vacuolar pr | 4 | 1.598965 | 0.570784 |
| A0A8M9QC | vwa2 | von Willebr | 14 | 2.711466 | 0.347683 |
| Q7ZUK7 | waca | WW domai | 3 | 1.468851 | 0.915926 |
| Q5SP67 | wdr26 | WD repeat | 2 | 2.578264 | 1.015136 |
| A0A8M1QI | wdr26a | WD repeat | 3 | 1.834377 | 0.441547 |
| A0A8M1P7 | wdr36 | WD repeat | 2 | 2.156612 | 0.68107 |

| | | | | | |
|---|---|---|---|---|---|
| Q7ZTY9 | wdr57 | Small nucle | 6 | 1.371696 | 0.348907 |
| E7FFE0 | whsc2 | Negative el | 1 | 1.71709 | 0.59291 |
| A0A8M9PX | wnk1b | non-specifi | 2 | 1.71088 | 0.410084 |
| Q75XT1 | wu:fa01e0: | Acetylcholi | 2 | 1.530358 | 0.345991 |
| Q6AZC1 | wu:fa14g0: | 26S protea | 19 | 2.792549 | 0.480879 |
| Q503J3 | wu:fa17b1( | Zgc:110551 | 6 | 1.503609 | 0.622415 |
| E7F5V5 | wu:fa96a0! | Actinodin1 | 29 | 1.458818 | 0.330914 |
| Q6TH08 | wu:fb08f03 | vacuolar pr | 3 | 2.008325 | 0.519934 |
| Q6IQQ0 | wu:fb13h0 | 60S ribosor | 8 | 1.388105 | 0.724418 |
| Q6P9P3 | wu:fb30f01 | Ubiquitin-li | 25 | 1.747176 | 0.400717 |
| Q6NV07 | wu:fb34a0( | DNA replica | 4 | 1.71972 | 1.122288 |
| Q6PBM0 | wu:fb39a1: | NADH dehy | 18 | 1.863373 | 0.383336 |
| Q6P937 | wu:fb61d0 | Clathrin ligl | 6 | 1.814524 | 0.343691 |
| Q8JH29 | wu:fb62f06 | Angiotensir | 10 | 2.078748 | 0.524703 |
| B0S6V7 | wu:fb65b0. | Phosphatid | 1 | 3.460799 | 0.349897 |
| A5PLJ7 | wu:fb77f12 | non-specifi | 5 | 2.024287 | 0.711877 |
| Q7SZC9 | wu:fb97a0! | Heterogen( | 2 | 2.115441 | 1.532515 |
| Q803U0 | wu:fc11d0( | SCY1-like 3 | 1 | 1.990555 | 0.516331 |
| Q6P951 | wu:fc31e0: | 6-phospho | 1 | 2.277442 | 1.604715 |
| Q6GQL6 | wu:fc38g1( | Dynactin 3 | 5 | 1.68497 | 0.443907 |
| Q568M0 | wu:fc47b08 | RAS-relatec | 5 | 1.78513 | 0.464872 |
| Q6DRF3 | wu:fc51f06 | Proteasom | 8 | 1.876454 | 1.344909 |
| Q6IQM8 | wu:fc52e0: | Caveolin | 2 | 3.090291 | 1.943676 |
| Q8AYJ5 | wu:fc60a07 | Mkln1 prot | 1 | 1.719681 | 0.537398 |
| Q7ZW32 | wu:fc75g08 | Ube2g1 pro | 2 | 1.976654 | 0.515125 |
| Q6DRD2 | wu:fc85c1( | 26S protea | 11 | 1.973419 | 0.597986 |
| Q802C9 | wu:fd15g0: | RNA helicas | 15 | 1.825743 | 0.430575 |
| Q4VBU0 | wu:fd20d0 | 2-oxoisova | 8 | 1.883112 | 0.483603 |
| Q7ZUS2 | wu:fd59g0! | Nucleobinc | 2 | 1.494737 | 0.362254 |
| Q7ZV77 | wu:fe05d1( | Proteasom | 7 | 1.590481 | 0.333008 |
| Q7ZYX7 | wu:fi03c01 | 26S protea | 10 | 1.85306 | 0.437504 |
| Q7ZW96 | wu:fi22c06 | Low densit | 3 | 1.449726 | 0.809442 |
| Q08BA7 | wu:fi23e07 | Zgc:154073 | 2 | 3.135653 | 0.575753 |
| Q7ZUF5 | wu:fi43d12 | Transmeml | 1 | 1.644546 | 1.130888 |
| Q7T317 | wu:fi98c09 | Matrix met | 3 | 2.488018 | 2.58072 |
| Q6IQ72 | wu:fj14c10 | Proteasom | 16 | 2.326798 | 0.398638 |
| Q7ZVT4 | wu:fj17e04 | Kinesin ligh | 7 | 1.497205 | 0.47893 |
| F1R8J6 | wu:fj17f04 | Palmitoyl-p | 1 | 1.628797 | 0.335833 |
| Q6NWL6 | wu:fj17f09 | Ubiquitin c | 5 | 2.501781 | 0.348688 |
| Q6GML0 | wu:fj54b01 | Zgc:91860 | 3 | 1.776391 | 0.49141 |
| Q7SZZ5 | wu:fj61c06 | Vesicle-ass | 4 | 1.394538 | 0.386032 |
| Q7SX91 | wu:fj66c04 | Ribosome I | 8 | 1.592111 | 0.418393 |
| Q6PGU3 | wu:fk52b1: | Regulation | 2 | 1.953431 | 0.357383 |
| Q6P013 | wu:fl03a12 | Transcripti | 1 | 1.431039 | 0.734237 |
| Q567F7 | wu:fu56g1: | Zgc:112077 | 1 | 1.649424 | 0.76102 |
| Q5PZ43 | xirp1 | Xin actin-bi | 15 | 2.750108 | 1.384185 |
| A8DZH4 | xpr1 | Xenotropic | 1 | 2.544229 | 0.758666 |

| | | | | | |
|---|---|---|---|---|---|
| Q1L8J7 | yap1 | Transcriptio | 4 | 2.257561 | 0.384332 |
| A0A8M1P8 | ypel5 | Protein yip | 1 | 2.339749 | 1.158608 |
| E7F1H9 | ythdf2 | YTH domai | 3 | 1.977161 | 0.373324 |
| A0A8M1PC | zc3h13 | zinc finger | 3 | 1.363103 | 0.713608 |
| F1QXD3 | zdhhc15b | Palmitoyltr | 1 | 1.414967 | 0.554179 |
| Q90XA8 | zfHSF2 | Heat shock | 1 | 1.676635 | 0.607689 |
| A0A8M2BJ | zfand5a | AN1-type z | 1 | 3.758308 | 1.413122 |
| A0A8M9QF | zfyve26 | Zinc finger | 2 | 1.393242 | 0.603262 |
| Q6DC38 | zgc:10085 | BAG family | 4 | 2.856301 | 0.559749 |
| Q6DEI0 | zgc:100903 | Phospholip | 3 | 2.047807 | 0.86613 |
| Q641M4 | zgc:10091 | Zgc:10091 | 1 | 1.779816 | 0.66152 |
| Q68EJ4 | zgc:101131 | Transcriptio | 2 | 1.728262 | 0.874871 |
| Q5XJU9 | zgc:101541 | Galactokin | 1 | 1.658541 | 1.132206 |
| Q66IF0 | zgc:101565 | Zgc:10156 | 6 | 2.348945 | 0.352386 |
| F1RC64 | zgc:101574 | Glutathion | 2 | 1.38105 | 0.415706 |
| Q5RKQ5 | zgc:101581 | Zgc:101581 | 2 | 1.45115 | 0.836118 |
| Q5XJP4 | zgc:10171C | Zgc:10171C | 5 | 1.575741 | 0.430492 |
| A0A0R4ICJ | zgc:10178C | Beta-1,4-ga | 1 | 3.246361 | 0.365851 |
| Q76C07 | zgc:10179 | Actin-bindi | 2 | 1.313974 | 0.683841 |
| Q66IA7 | zgc:101894 | Ubiquitous | 1 | 1.993912 | 0.519689 |
| Q5XJD0 | zgc:10342 | Heat shock | 2 | 2.022352 | 0.861379 |
| Q5XJC6 | zgc:103434 | ethanolam | 7 | 2.840246 | 0.486731 |
| A2BEV6 | zgc:103442 | Novel prot | 9 | 2.247991 | 0.705922 |
| Q5XJB0 | zgc:103467 | Zgc:10346 | 18 | 1.3756 | 0.552508 |
| Q5PR58 | zgc:10348 | Armadillo r | 1 | 1.328795 | 1.030677 |
| Q66I87 | zgc:10349C | Glyoxalase | 5 | 1.440405 | 0.575513 |
| Q5XJ58 | zgc:103632 | Thioredoxi | 3 | 2.575526 | 0.547506 |
| Q5U3E8 | zgc:103652 | Coatomer s | 5 | 1.427652 | 1.193417 |
| Q568W5 | zgc:109763 | MCL1 apop | 1 | 1.698617 | 0.537315 |
| A0A8M1P3 | zgc:10988 | Wiskott-Al | 1 | 1.534808 | 0.598043 |
| Q503V2 | zgc:110064 | Apolipopro | 1 | 2.343208 | 0.617733 |
| Q503R3 | zgc:11027C | Zgc:11027C | 2 | 1.495142 | 0.347402 |
| Q568H8 | zgc:11028 | DNA-direct | 3 | 1.795131 | 0.652786 |
| Q568H2 | zgc:11030C | Cytochrom | 1 | 1.496729 | 0.822145 |
| B8A6A3 | zgc:11031 | NFU1 iron- | 3 | 1.454857 | 0.382514 |
| Q4VBR6 | zgc:110731 | Endoplasm | 3 | 1.47091 | 0.553403 |
| Q4VBI5 | zgc:11234C | NADH deh | 8 | 1.459865 | 0.607721 |
| Q4KMD0 | zgc:112352 | Coronin | 2 | 1.888904 | 1.462194 |
| Q501W9 | zgc:112501 | DNA-direct | 3 | 1.789711 | 0.3641 |
| Q566S4 | zgc:11252C | NADH deh | 6 | 1.504109 | 0.42628 |
| Q6Q419 | zgc:11393 | 40S riboso | 1 | 1.491148 | 0.323362 |
| Q4VBU7 | zgc:11416 | Cytochrom | 2 | 1.690076 | 0.382667 |
| Q5BJA2 | zgc:114172 | NADH deh | 2 | 1.828943 | 1.342528 |
| Q32LT0 | zgc:123293 | Sorting nex | 2 | 2.546899 | 0.637975 |
| A0A0R4IJL | zgc:123333 | Alpha-gala | 3 | 1.725596 | 0.954128 |
| Q29RE8 | zgc:13662 | Neurofilam | 5 | 1.464601 | 0.372147 |
| Q29RA2 | zgc:13690 | Transitiona | 8 | 1.710908 | 0.631692 |

| | | | | | |
|---|---|---|---|---|---|
| Q08CD8 | zgc:153231 | GTP-bindin | 4 | 1.703998 | 0.686681 |
| Q0P459 | zgc:15339C | 28S ribosor | 1 | 1.493054 | 0.838777 |
| Q0P449 | zgc:153411 | Zgc:153411 | 1 | 1.86709 | 1.085432 |
| Q08BF2 | zgc:15386C | Zgc:15386C | 2 | 1.826434 | 0.761326 |
| A0JML6 | zgc:153955 | Tuftelin 1a | 1 | 2.149037 | 0.690668 |
| Q05AI8 | zgc:154009 | Zgc:154009 | 7 | 1.770036 | 0.536634 |
| Q0P3U1 | zgc:154087 | Dehydroge | 3 | 1.447885 | 0.864158 |
| Q08BV6 | zgc:154095 | creatine kir | 11 | 1.397061 | 0.326662 |
| A1A5V2 | zgc:158252 | Zgc:158252 | 2 | 3.676147 | 0.960655 |
| A1L1S0 | zgc:158286 | Leucine ricl | 1 | 1.653267 | 0.753204 |
| A1L1T1 | zgc:158319 | Niban apop | 4 | 2.414785 | 0.388764 |
| A1A5X1 | zgc:158342 | non-specifi | 4 | 1.629794 | 0.611425 |
| A1L1V1 | zgc:158397 | Farnesyltra | 7 | 1.627261 | 0.477567 |
| A4IG34 | zgc:16215C | Perilipin 6 | 2 | 1.920577 | 0.795444 |
| A3KNN5 | zgc:162272 | 26S protea | 3 | 2.150111 | 0.324715 |
| A3KNN6 | zgc:162277 | Diphthamic | 3 | 1.499011 | 0.359481 |
| A3KP28 | zgc:162895 | Checkpoint | 1 | 1.445968 | 1.04193 |
| A4QN91 | zgc:163073 | Zgc:163073 | 1 | 1.591081 | 1.324933 |
| B0JZC1 | zgc:171911 | 39S ribosor | 4 | 2.006862 | 0.334628 |
| A8E5P7 | zgc:174917 | Zgc:174917 | 1 | 1.765273 | 1.349973 |
| Q5U393 | zgc:175013 | Cleavage ar | 3 | 1.900126 | 0.659219 |
| B3DJF3 | zgc:194985 | Zgc:194985 | 7 | 1.830227 | 0.324484 |
| Q7ZW05 | zgc:55441 | YTH domai | 3 | 1.632811 | 0.417585 |
| Q7ZVW5 | zgc:55557 | Zgc:55557 | 1 | 1.725314 | 0.980894 |
| A0A8M3AL | zgc:55582 | myomegali | 4 | 1.85958 | 0.412681 |
| Q7ZYY0 | zgc:55874 | Acyl-CoA d | 4 | 1.993709 | 0.506924 |
| Q803B3 | zgc:55949 | Ubiquitin c | 6 | 5.322512 | 0.435204 |
| A0A8M9PI! | zgc:56064 | Serine/thre | 10 | 1.389135 | 0.511451 |
| Q7ZU93 | zgc:56066 | Ribonuclop | 2 | 2.197201 | 0.691181 |
| Q7ZUU4 | zgc:56091 | DnaJ (Hsp4 | 2 | 2.211865 | 0.475391 |
| Q7ZUS7 | zgc:56227 | Serine/thre | 3 | 2.058502 | 0.50585 |
| A0A8M9PV | zgc:56304 | uncharacte | 2 | 1.311182 | 0.529175 |
| Q7ZUZ2 | zgc:56317 | Actin relate | 6 | 1.817392 | 0.625773 |
| Q7ZUJ8 | zgc:56374 | Proteasom | 5 | 1.466853 | 1.229375 |
| Q7ZU69 | zgc:56377 | 26S protea | 17 | 1.801272 | 0.686406 |
| Q6NV33 | zgc:56380 | Isocitrate d | 10 | 1.594474 | 0.51104 |
| Q6IQE3 | zgc:56411 | Tetratricop | 2 | 2.18989 | 0.495391 |
| Q7ZWE5 | zgc:56471 | 26S protea | 19 | 1.987462 | 0.332306 |
| E9QC42 | zgc:56506 | Synaptobre | 2 | 2.160743 | 0.944487 |
| Q7ZWD1 | zgc:56516 | Proteasom | 1 | 1.688254 | 1.944745 |
| Q7ZW92 | zgc:56677 | Serine/thre | 4 | 1.435723 | 0.722678 |
| Q7ZW89 | zgc:56686 | SEC22 vesi | 1 | 1.403968 | 0.586203 |
| Q7SY50 | zgc:63505 | Zgc:63505 | 2 | 1.330144 | 2.041796 |
| F1RA91 | zgc:63532 | Argininosu | 4 | 1.669359 | 0.477235 |
| F1QXM3 | zgc:63624 | TBC1 doma | 2 | 1.631806 | 1.290935 |
| Q7T3B1 | zgc:63995 | 26S protea | 21 | 2.08099 | 0.440073 |
| Q6PFT0 | zgc:64103 | Flotillin | 5 | 1.877754 | 0.328161 |

| | | | | | |
|---|---|---|---|---|---|
| A9JT54 | zgc:64136 | Zgc:64136 | 1 | 3.119033 | 0.361521 |
| Q6PHK7 | zgc:65921 | 26S protea | 24 | 2.256196 | 0.327982 |
| Q6PHE7 | zgc:65973 | DNA-direct | 3 | 1.724167 | 0.460687 |
| Q6PHI7 | zgc:65981 | phenylalan | 19 | 1.396994 | 0.40543 |
| Q6PC76 | zgc:73072 | Golgi reass | 2 | 1.750369 | 0.695116 |
| Q6PC49 | zgc:73107 | NADH dehy | 10 | 1.787884 | 0.923234 |
| Q6NYL6 | zgc:76981 | Zgc:76981 | 3 | 1.443106 | 0.330514 |
| Q7ZTZ0 | zgc:76983 | DNA-direct | 3 | 1.598517 | 0.389545 |
| Q6P0I2 | zgc:77139 | Proteasom | 9 | 2.206988 | 0.48152 |
| Q6P4V4 | zgc:77673 | Catenin, be | 5 | 1.996484 | 0.878243 |
| Q7ZUG6 | zgc:77683 | 60S ribosor | 11 | 1.479245 | 0.793012 |
| Q6P6F0 | zgc:77702 | 40S ribosor | 5 | 1.525455 | 0.549504 |
| Q7ZUB0 | zgc:77943 | Atrial myos | 17 | 1.587272 | 0.606343 |
| Q6NWI5 | zgc:85685 | methylcrot | 12 | 1.408814 | 0.430958 |
| Q6NWE0 | zgc:85789 | Ester hydrc | 2 | 1.480177 | 0.583694 |
| Q6NSM6 | zgc:85963 | Zgc:85963 | 9 | 1.383339 | 0.422289 |
| Q6IQH8 | zgc:86757 | Zgc:86757 | 5 | 2.497334 | 1.106118 |
| Q6IQH4 | zgc:86762 | 26S protea | 7 | 1.638261 | 0.918255 |
| Q7ZV16 | zgc:86802 | Elongation | 10 | 1.528672 | 0.460503 |
| Q6IQC6 | zgc:86833 | 26S protea | 8 | 2.156084 | 0.667438 |
| Q5I0F8 | zgc:86860 | Cell divisior | 3 | 1.670069 | 0.675758 |
| Q68EH3 | zgc:91894 | E74-like fac | 1 | 2.205114 | 0.876222 |
| Q7SZZ7 | zgc:91931 | AP-4 comp | 1 | 2.629182 | 0.509114 |
| Q6DHT4 | zgc:92082 | Aldehyde d | 21 | 1.653173 | 0.424762 |
| Q6DHM1 | zgc:92229 | DNAation f | 2 | 1.480583 | 0.840127 |
| Q5PQY9 | zgc:92239 | Muscle-rela | 16 | 1.429886 | 0.743646 |
| Q6DHK2 | zgc:92259 | Nucleopori | 4 | 1.462947 | 0.541758 |
| A0A8M9PK | zgc:92287 | uncharacte | 4 | 1.634919 | 0.860672 |
| Q6P3G3 | zgc:92437 | UBX domai | 4 | 2.238191 | 0.388665 |
| Q6DH80 | zgc:92608 | Lysozyme g | 4 | 2.874685 | 0.531714 |
| Q6DH35 | zgc:92663 | Zgc:92663 | 1 | 3.317085 | 0.760497 |
| Q6DH25 | zgc:92674 | Ependymin | 1 | 1.920114 | 1.341136 |
| Q6DGY8 | zgc:92716 | Proteasom | 12 | 2.702683 | 0.408721 |
| Q502P2 | zmp:00000 | Zgc:11196( | 11 | 1.900136 | 0.735523 |
| A0A8M9QE | zmp:00000 | A-kinase ar | 3 | 2.307844 | 0.76543 |
| A0A8M9PZ | zmp:00000 | uncharacte | 1 | 1.623123 | 0.596724 |
| A0A8M9Q9 | zmp:00000 | uncharacte | 2 | 2.950537 | 1.652827 |
| A0A8M9Q3 | znf131 | zinc finger | 1 | 1.30525 | 1.07412 |
| Q08CN9 | znrf2 | E3 ubiquitir | 1 | 1.75207 | 0.326635 |
| Q6DHC4 | Bf-2 | Compleme | 1 | 1.536419 | -0.47537 |
| Q8UVG6 | CRBP | Cellular ret | 5 | 2.062156 | -0.77433 |
| Q803J0 | Cyp2J1 | Cyp2j25 pr | 1 | 1.481037 | -0.37306 |
| Q7T3D6 | Gpc1b | Glypican-1 | 2 | 2.118783 | -0.39693 |
| Q1LVG4 | IRb | Tyrosine-pr | 6 | 1.33583 | -0.3971 |
| A0A8M1RS | LOC100007 | protein PF( | 2 | 2.658231 | -0.432 |
| A0A8M9P9 | LOC10014 | formin-like | 1 | 2.025283 | -0.68039 |
| A0A8M9QE | LOC10015( | metal trans | 3 | 1.882154 | -0.4465 |

| | | | | | |
|---|---|---|---|---|---|
| A0A8M9Q( | LOC10033( | uncharacte | 28 | 1.368243 | -0.52059 |
| A0A8M1P4 | LOC100332 | uncharacte | 2 | 1.389924 | -0.48356 |
| A0A8M9PT | LOC10033: | integrin alp | 1 | 1.885004 | -0.39658 |
| A0A8M1RF | LOC100334 | solute carri | 3 | 2.342339 | -0.45122 |
| A0A8M1RS | LOC10053! | uncharacte | 16 | 4.733054 | -0.65957 |
| A0A8M9P4 | LOC10053( | uncharacte | 3 | 3.003095 | -0.60532 |
| A0A8M1RF | LOC100537 | solute carri | 1 | 2.034976 | -0.54617 |
| A0A8M9Q8 | LOC10188: | ATP-bindin | 1 | 1.895103 | -0.44195 |
| A0A8M2BH | LOC10188: | uncharacte | 1 | 2.008505 | -0.54927 |
| A0A8M9PC | LOC101884 | collagen alp | 5 | 1.6219 | -0.33496 |
| A0A8M9PK | LOC103908 | uncharacte | 1 | 1.919581 | -0.56912 |
| A0A8M9P6 | LOC103908 | MYND-type | 2 | 3.715451 | -0.63039 |
| A0A8M9PN | LOC10390! | synaptotag | 1 | 1.690825 | -0.35836 |
| A0A8M9Q( | LOC10391( | huntingtin- | 3 | 2.010131 | -0.4296 |
| A0A8M9P9 | LOC10817! | immune-as | 1 | 1.444062 | -1.04702 |
| B3DJB7 | LOC571819 | Similar to a | 2 | 1.32484 | -0.77587 |
| E7FGA4 | NRG2b | Neuregulin | 1 | 1.423437 | -0.38922 |
| Q7ZUX4 | RDHB | Retinol deh | 7 | 1.52462 | -0.41951 |
| A3KH13 | S100A1 | Protein S1( | 1 | 1.834666 | -0.41604 |
| Q9PUQ6 | Sk-Tmod | Sk-tropomo | 14 | 2.987344 | -0.53431 |
| D3XD61 | Ugt1b2 | UDP-glucur | 3 | 2.354456 | -0.69461 |
| D3XD87 | Ugt2b3 | UDP-glucur | 5 | 2.091718 | -0.60717 |
| A0A8M3BE | a1cf | APOBEC1 c | 7 | 1.4516 | -0.47814 |
| A0A8M2B3 | abca5 | ATP-bindin | 4 | 1.376731 | -0.33081 |
| A0A8M2BF | acsl1a | Long-chain | 8 | 1.937705 | -0.39294 |
| Q7SYD3 | actn3 | Actinin alpl | 33 | 3.586535 | -0.6321 |
| A0A8M1RT | adcy9 | adenylate c | 2 | 1.542872 | -0.38071 |
| Q7T2J4 | adh8b | Alcohol del | 17 | 2.373289 | -0.52782 |
| Q7ZW00 | agk | Acylglycerc | 1 | 1.734651 | -0.58924 |
| Q6NYE4 | agmo | Alkylglycer | 2 | 1.370107 | -0.45637 |
| A0JMM2 | agpat2 | 1-acyl-sn-g | 1 | 1.313812 | -0.69822 |
| A0A8M2BG | apba2b | amyloid be | 1 | 2.442829 | -1.4285 |
| A0A8M2BF | apbb2b | amyloid be | 2 | 3.642348 | -0.42513 |
| A0A8M1NA | apoda.1 | Apolipopro | 4 | 1.46547 | -0.36928 |
| A0A8M2B7 | aqp7 | aquaporin- | 2 | 1.485984 | -0.71965 |
| A0A8M2BD | arhgef25b | rho guanin | 1 | 2.988217 | -0.38616 |
| Q7ZWH4 | arpc5b | Actin-relate | 3 | 2.247925 | -0.53915 |
| A0A8M9QI | atg7 | Ubiquitin-li | 3 | 1.451747 | -0.37933 |
| A0A8M1M: | atp13a1 | manganese | 5 | 1.720477 | -0.35365 |
| A0A8M2BL | atp9a | Phospholip | 2 | 1.821965 | -0.47696 |
| A8WGA6 | b44 | Tnnt3b pro | 46 | 4.008556 | -1.16501 |
| E3W9A6 | bZ1N7.1 | Sema dom | 1 | 3.447964 | -0.41558 |
| A0A8M1PR | bnip1b | vesicle trar | 1 | 2.279422 | -0.33004 |
| A0A8M3AF | bnip2 | BCL2/aden | 1 | 1.338913 | -0.6092 |
| Q58EQ7 | bnip3l | BCL2 intera | 2 | 1.624294 | -0.90672 |
| A0A8M2BG | brpf3a | bromodom | 1 | 1.403634 | -0.40852 |
| A0A8M2BF | cacna1sa | Voltage-de | 4 | 2.035062 | -0.5271 |

| | | | | | |
|---|---|---|---|---|---|
| A0A8M9P5 | camta1a | calmodulin | 1 | 1.848845 | -0.45807 |
| F8W3K2 | card15 | NOD2a pro | 1 | 1.887043 | -0.38715 |
| Q6DHU3 | cb112 | Zgc:92061 | 22 | 1.446381 | -0.55697 |
| B8A568 | cb20 | Myosin, he | 34 | 1.937313 | -0.61541 |
| Q6P980 | cb463 | Superoxide | 8 | 2.229921 | -0.43809 |
| Q7ZUC4 | ccdc6 | Ccdc6a pro | 7 | 2.734926 | -0.43307 |
| A0A8M2B6 | ccsapa | centriole, c | 1 | 1.310777 | -0.4667 |
| A0A8M2BL | cd151 | Tetraspanin | 1 | 2.500255 | -0.35434 |
| Q6DBW9-2 | cd99l2 | Isoform 2 c | 1 | 1.389446 | -0.49247 |
| A0A0R4IVA | cdc14ab | Dual specif | 1 | 1.491613 | -0.52483 |
| A6H8U1 | cdip1 | Cell death-i | 1 | 1.635158 | -0.32463 |
| A0A8M1RF | cers3a | ceramide s | 1 | 1.371299 | -0.7241 |
| A0A8M3AV | clcn6 | Chloride ch | 1 | 1.532278 | -0.41563 |
| Q7T2E7 | cldn15l | Claudin | 1 | 1.709804 | -0.3922 |
| Q6AXK7 | cldnd1b | Zgc:100913 | 1 | 2.91018 | -0.4444 |
| Q6NXB5 | clic5 | Chloride in | 7 | 1.348528 | -0.46864 |
| Q5U3P5 | clic5a | Chloride in | 5 | 2.160382 | -0.3467 |
| Q7ZW34 | cntn5 | Contactin-5 | 1 | 1.610805 | -0.42402 |
| A0A8M2B6 | col17a1a | collagen, ty | 12 | 2.759287 | -0.47759 |
| A0A8M9Q1 | col19a1 | collagen alp | 1 | 2.729682 | -0.82728 |
| A0A8M9PC | col4a3bpb | collagen ty | 1 | 1.493129 | -0.379 |
| A0A8N7T6I | colgalt1 | procollagen | 3 | 1.38088 | -0.4839 |
| A4IG53 | comtb | Catechol O | 3 | 1.902765 | -0.40444 |
| Q6DH88 | cox20 | Cytochrom | 1 | 1.783382 | -0.34394 |
| B8JLQ9 | cpo | Carboxypep | 4 | 1.465036 | -0.55648 |
| F6P7Z9 | cpt1al | carnitine O | 1 | 1.71836 | -0.55798 |
| A0A8M9PF | cpt1cb | carnitine O | 4 | 1.836099 | -0.4181 |
| Q7SZC2 | cse1l | Exportin-2 | 18 | 1.702936 | -0.37358 |
| A0A8M1N7 | cst3 | cystatin C p | 3 | 2.500517 | -0.40159 |
| Q0P487 | cyb5r2 | NADH-cyto | 2 | 1.374487 | -0.40947 |
| Q8UUR3 | cygb1 | Cytoglobin- | 6 | 1.451179 | -0.40118 |
| Q6PGV7 | cyp2j22 | Cytochrom | 2 | 1.911116 | -0.97491 |
| Q5TZ81 | cyp2j29 | Cytochrom | 1 | 1.503889 | -0.56536 |
| A0A8M1N2 | cyp2k16 | cytochrome | 4 | 1.597929 | -0.54569 |
| A0JMQ6 | cyp4v2 | Cyp4v2 pro | 1 | 2.644641 | -0.77326 |
| Q1LXJ7 | cyt1l | type I cytok | 57 | 1.966635 | -0.46364 |
| Q6P3J0 | dgat1 | O-acyltrans | 2 | 1.707784 | -0.71856 |
| Q4V9F0 | dgat2 | Diacylglyce | 2 | 2.379564 | -0.83445 |
| Q7SXF1 | dhcr7 | 7-dehydroc | 4 | 1.896339 | -0.8502 |
| A0A8N7TE5 | dhx30 | RNA helicas | 1 | 1.657611 | -0.38046 |
| A1L1P7 | dnlz | DNL-type z | 2 | 1.641889 | -0.40983 |
| A0A8M1RK | dnttip1 | Deoxynucle | 2 | 1.496498 | -0.56963 |
| A0A8M9QI | ela3l | elastase 3 l | 11 | 1.389987 | -0.44134 |
| A0A8M3AY | elavl3 | ELAV-like p | 4 | 1.462915 | -0.5233 |
| Q8JG61 | epb41 | Protein 4.1 | 4 | 1.97267 | -0.3467 |
| A0A8M2B9 | epha2b | receptor pr | 1 | 1.662107 | -0.53044 |
| A0A8M3AV | epha3 | receptor pr | 2 | 2.457473 | -0.46365 |

| | | | | | |
|---|---|---|---|---|---|
| A0A8M3B8 | eps8l1 | eps8-like1 i | 1 | 1.750652 | -0.42472 |
| Q4V8Y6 | ergic1 | Endoplasm | 2 | 2.535776 | -0.3528 |
| E7FD79 | erich1 | Glutamate | 1 | 1.592188 | -0.78322 |
| A0A8M6YV | erp27 | endoplasm | 5 | 1.552179 | -0.72057 |
| Q66I72 | f11r | Junctional a | 5 | 2.117959 | -0.35561 |
| Q6NX10 | fa22e07 | ADP/ATP tr | 46 | 1.748686 | -0.35193 |
| Q4VBT1 | fabp1b.1 | Fatty acid b | 15 | 1.694858 | -0.69812 |
| A0A8M6Z9 | fam135a | protein FAI | 1 | 1.588579 | -0.51917 |
| A0A8M1PD | fastkd1 | FAST kinase | 1 | 1.467653 | -0.46347 |
| A0A8N7UV | fastkd5 | FAST kinase | 3 | 2.087509 | -0.46637 |
| A3KP44 | fb59c09 | Zgc:162938 | 8 | 1.720925 | -0.36724 |
| Q2HPG2 | fb78h12 | LGIa (Fragn | 1 | 1.721924 | -0.33923 |
| Q6NUW9 | fggy | FGGY carbo | 1 | 1.380812 | -0.37945 |
| A8WG70 | fj47b06 | DnaJ (Hsp4 | 1 | 1.337447 | -0.46772 |
| F1R3E6 | foxo4 | Forkhead b | 3 | 2.593898 | -0.34408 |
| Q6PH19 | gatm | Glycine am | 9 | 2.237195 | -0.51085 |
| A0A8M2BK | gdpd2_1 | glyceropho | 1 | 1.970743 | -0.69342 |
| A0A8M6Z9 | gdpd3b | glyceropho | 2 | 1.615233 | -0.6119 |
| A0A8M9PR | glipr2 | ancylostom | 2 | 2.545217 | -0.8299 |
| A0A8M1PF | gmds | GDP-manno | 6 | 2.238039 | -0.36923 |
| A0A8M9PT | golga7 | golgin subf | 1 | 1.846962 | -0.52828 |
| Q6DG38 | gpat3 | Glycerol-3- | 2 | 1.487247 | -0.4626 |
| A0A8M1P8 | gpatch2 | G patch do | 1 | 1.499796 | -0.49167 |
| A0A8M1NI | gpx3 | Glutathione | 1 | 2.345846 | -0.32862 |
| Q68EG5 | grb10 | Grb10 prot | 2 | 1.362623 | -0.4574 |
| A0A8N7TD | gtf3c4 | general tra | 1 | 2.028666 | -0.35053 |
| A0A8M2BE | gtpbp1 | GTP-bindin | 4 | 2.280206 | -0.57861 |
| Q5BLF6 | hbbe3 | Hemoglobi | 15 | 2.298039 | -0.51704 |
| A0A8M3B4 | hectd4 | probable E3 | 9 | 1.372015 | -0.32411 |
| A0A8M1NE | hexdc | beta-N-ace | 3 | 2.039813 | -0.33604 |
| A0A8M2B8 | hm13 | minor histo | 2 | 1.993975 | -0.37942 |
| Q9I9N0 | hm:zeh018 | Dap1b | 1 | 1.736527 | -0.69512 |
| A6H8Q3 | hm:zeh105 | Zgc:165344 | 39 | 3.319957 | -0.8409 |
| Q6PUF3 | hsd11b1l | Hydroxyste | 6 | 1.428035 | -0.60449 |
| A0A8M3AX | hsd11b1lb | hydroxyste | 1 | 1.838003 | -0.70532 |
| A0A8M3AN | htr3a | 5-hydroxyt | 2 | 2.342221 | -0.42895 |
| A0A8M9PN | igfn1.1_1 | immunoglo | 2 | 3.369336 | -0.95227 |
| A0A8M9QA | igfn1.1 | immunoglo | 39 | 3.853428 | -0.94838 |
| A0A8M2BC | igfn1.3 | immunoglo | 38 | 4.06083 | -0.74074 |
| A0A8M1RS | il17rel | putative int | 1 | 1.809619 | -1.05647 |
| Q7T162 | im:715302 | F-box and l | 1 | 2.260388 | -0.42955 |
| Q568D1 | im:715508 | Tetraspanin | 2 | 2.182073 | -0.44071 |
| A0A8M1RN | itga10 | integrin alp | 1 | 1.940713 | -0.7293 |
| B3DIV2 | itgb1_1 | Integrin be | 8 | 2.051766 | -0.38515 |
| A3KPA0 | jam3b | Junctional a | 2 | 2.052639 | -0.60483 |
| B3DIV9 | klhl40a | Kelch-like p | 1 | 3.575372 | -1.61799 |
| F1QEG2 | klhl41b | Kelch-like p | 11 | 4.39544 | -2.87011 |

| | | | | | |
|---|---|---|---|---|---|
| F1REV3 | krit1 | Krev intera | 1 | 1.636689 | -0.91771 |
| Q7SXJ0 | lgals1l3 | Galectin | 2 | 1.420784 | -2.05417 |
| Q1LWG4 | lpcat1 | Lysophospl | 1 | 1.633029 | -0.45342 |
| A0A8M3B7 | man2c1 | alpha-manı | 6 | 1.386912 | -0.4621 |
| A0A8M1NU | map3k5 | mitogen-ac | 2 | 2.009631 | -0.35021 |
| A0A8M1P7 | mbd1a | methyl-CpC | 1 | 1.584327 | -0.46871 |
| A0A8M1NZ | mbpb | Myelin bas | 7 | 1.883368 | -0.34993 |
| A0A8N7TB | mctp2a | multiple C2 | 1 | 1.866263 | -0.36966 |
| Q7ZW91 | mobkl2a | MOB1, Mp | 2 | 1.660039 | -0.46607 |
| A0A8M1P3 | mogat2 | Acyltransfe | 6 | 1.3028 | -0.51743 |
| A0A8M1P7 | mon1a | Vacuolar fu | 1 | 2.031468 | -0.72748 |
| A0A8M9PT | mslna | mesothelin | 4 | 1.448891 | -0.92615 |
| Q9MIX8 | mt-cyb | Cytochrom | 1 | 1.748492 | -0.60241 |
| Q58EJ9 | mtarc1 | Mitochond | 7 | 1.446931 | -0.44916 |
| A0A0R4IVV | mttp | Microsoma | 17 | 1.325971 | -0.59995 |
| Q5RKQ2 | mtx1 | Metaxin | 1 | 1.675756 | -0.46982 |
| A0A8M9Q1 | mybpc2a | myosin-bin | 7 | 1.478873 | -1.30256 |
| A0A8M1N4 | mybpc2b | myosin-bin | 30 | 2.449293 | -0.60357 |
| A0A8M6Z3 | mybpha | uncharacte | 7 | 1.529181 | -0.71539 |
| A0A8M3BC | mylka | Myosin ligh | 3 | 2.506372 | -0.62032 |
| A0A8M3BA | mylkb | myosin ligh | 5 | 1.739814 | -0.44258 |
| E7F9L8 | myo1d | Unconvent | 10 | 1.643126 | -0.36415 |
| A0A8M3AF | myo5ab | unconventi | 1 | 2.615823 | -0.40621 |
| A0A8M1NU | myom1a | myomesin | 44 | 4.245964 | -0.89107 |
| A0A8M1PA | myom2a | uncharacte | 48 | 2.483652 | -0.58443 |
| A0A8M1RF | myom2b | myomesin- | 34 | 3.254915 | -0.85389 |
| A5PLA8 | nadsyn1 | Glutamine- | 1 | 1.44551 | -0.69395 |
| A0A8M3B1 | naif1 | Nuclear ap | 2 | 4.843004 | -2.57961 |
| Q0P4A4 | nat14 | Probable N | 1 | 1.348855 | -0.38764 |
| E7FAW3 | nbeal2 | Neurobeac | 2 | 1.358218 | -0.43938 |
| Q7SXL4 | ndpkz2 | Nucleoside | 41 | 3.458789 | -0.55179 |
| A0A8M9PY | neb | nebulin iso | 50 | 3.902817 | -2.13272 |
| A0A8M1RF | negaly6 | urokinase p | 2 | 1.775998 | -0.39681 |
| Q9IAD3 | nme2b1 | Nucleoside | 12 | 1.502513 | -0.32452 |
| A0A8M9PJ | nme3 | Nucleoside | 6 | 1.573098 | -0.59775 |
| A0A8M1P8 | nrap | nebulin-rel | 25 | 2.276484 | -1.04454 |
| A1XQX8 | nrxn3a | Neurexin-3 | 2 | 1.385337 | -0.37442 |
| B8A569 | ns:zf-e68 | Myosin, he | 396 | 2.633317 | -0.58729 |
| A0A8M1P7 | ogfr | opioid grov | 1 | 1.664511 | -0.34737 |
| A0A8M9PN | palm3 | uncharacte | 1 | 2.134852 | -0.49347 |
| A0A8M1P4 | pank4 | 4'-phospho | 2 | 2.641683 | -0.3464 |
| Q4VBJ8 | pbld | Zgc:11221C | 2 | 1.626926 | -0.5546 |
| Q7ZVK1 | pdcd4 | Programme | 2 | 2.302416 | -0.76917 |
| Q6P7E4 | pdlim7 | PDZ and LII | 6 | 2.323411 | -0.49286 |
| A0A8M9Q5 | pik3cg | phosphatid | 1 | 1.41283 | -0.54755 |
| P50392 | pla2g4a | Cytosolic p | 4 | 1.427095 | -0.36397 |
| A0A8M9P9 | pla2g4f.1 | Phospholip | 2 | 2.580417 | -0.43009 |

| | | | | | |
|---|---|---|---|---|---|
| A0A8M1N9 | pold3 | DNA polym | 2 | 1.801768 | -0.66332 |
| A0A8M1N6 | pon3.1 | Paraoxonas | 10 | 1.973732 | -0.32548 |
| Q568K2 | ppp1r11 | E3 ubiquitir | 1 | 1.897343 | -0.56644 |
| Q567W8 | ppp1r14a | Protein pho | 1 | 1.414385 | -0.34493 |
| A0A8N7T6 | prex1 | phosphatid | 2 | 1.480978 | -0.39651 |
| A0A8M3B9 | prkcq | Protein kin | 5 | 2.376456 | -0.33007 |
| A0A8M2BA | prr33 | muscle M-l | 17 | 1.307091 | -0.50437 |
| Q6IQG6 | prtfdc1 | Hypoxanth | 4 | 1.789592 | -0.52006 |
| Q803C9 | ptdss1 | Phosphatid | 1 | 1.784969 | -0.56882 |
| Q5U3W6 | pte2bl3 | Acyl-CoA th | 1 | 1.697003 | -1.0214 |
| A0A8M3B2 | ptgfrn | prostaglan | 9 | 2.502874 | -0.34968 |
| Q502I0 | ptk7a | Zgc:112211 | 4 | 1.48838 | -0.38715 |
| Q6NX92 | ptp4a3 | Protein tyr | 1 | 2.1263 | -0.86812 |
| A0A8M9QJ | ptprea | receptor-ty | 3 | 2.368457 | -0.37195 |
| A5PKS5 | ptps | 6-pyruvoyl | 2 | 1.400667 | -0.46817 |
| Q1LVA5 | rab25 | Novel prote | 4 | 1.621258 | -0.37565 |
| Q5XJS4 | rab34 | RAB34, me | 1 | 1.684303 | -0.50498 |
| A0A8M9PT | rapgef1b | rap guanine | 2 | 1.34194 | -0.46384 |
| E7F372 | rasa1b | RAS p21 pr | 5 | 1.618205 | -0.3636 |
| A0A8M2B9 | rerea | arginine-glu | 1 | 1.683613 | -0.56094 |
| A8Y5U1 | rimoc1 | RAB7A-inte | 3 | 1.852826 | -0.35469 |
| F1QC45 | rp2 | Protein XRI | 1 | 1.549391 | -0.405 |
| A0A0R4ICI5 | rrn3 | RNA polym | 3 | 1.680885 | -0.34173 |
| A0A8M9PV | ryr2a | ryanodine r | 2 | 2.710756 | -0.36676 |
| F1R442 | sb:cb171 | Arachidona | 7 | 2.145361 | -0.55535 |
| B1WB79 | sb:cb624 | Storkhead | 2 | 1.640079 | -0.46068 |
| A0A8M3AT | scn8ab | Sodium cha | 2 | 1.551328 | -0.32483 |
| A0A8M1ND | sec14l8 | SEC14-like | 1 | 1.523859 | -0.4891 |
| A0A8M1NV | sft2d3 | Vesicle tran | 1 | 2.152248 | -0.59883 |
| E7EZW5 | sgc1a1 | guanylate c | 1 | 1.588517 | -0.51213 |
| A0A8M2B2 | sh2d4a | SH2 domai | 1 | 1.744872 | -0.61517 |
| A0A8N7US | si:ch211-11 | cytochrome | 7 | 1.337951 | -0.38384 |
| A0A8M2B5 | si:ch211-12 | neuroblast | 45 | 2.058045 | -0.36565 |
| A0A8N7TF | si:ch211-17 | olfactomed | 1 | 1.778139 | -0.39222 |
| A7MD64 | si:ch211-17 | Si:ch211-17 | 3 | 1.740585 | -0.63858 |
| A0A8M1PS | si:ch211-19 | NACHT, LRI | 3 | 1.379646 | -0.44286 |
| A0A8M1PS | si:ch211-19 | probable N | 1 | 1.338619 | -0.65148 |
| A0A8M9PN | si:ch211-19 | si:ch211-19 | 1 | 1.705247 | -0.41192 |
| A0A8M2B9 | si:ch211-23 | uncharacte | 1 | 1.397296 | -0.51489 |
| A0A8M1RL | si:ch211-23 | SITS-bindin | 2 | 2.339902 | -0.33381 |
| Q4QRJ9 | si:ch211-26 | alkaline ph | 8 | 2.461122 | -0.59823 |
| A0A8M9P0 | si:ch211-26 | Triadin | 19 | 2.426775 | -0.64163 |
| Q503G4 | si:ch211-27 | Zgc:110621 | 1 | 1.334164 | -0.40077 |
| A0A8M1PS | si:ch211-81 | probable N | 1 | 1.458104 | -0.38717 |
| A0A8M9PR | si:ch211-87 | leucine-rich | 1 | 4.481368 | -1.00554 |
| A0A8M9PZ | si:ch211-93 | uncharacte | 10 | 1.761216 | -0.63078 |
| A0A8M9PJ | si:ch73-27 | uncharacte | 1 | 2.253029 | -0.50504 |

| | | | | | |
|---|---|---|---|---|---|
| A0A8M1RH | si:ch73-308 | lisH domair | 7 | 1.664074 | -0.48529 |
| A0A8M2BC | si:ch73-368 | histone H1- | 11 | 1.82757 | -0.34071 |
| A0A8M3AN | si:ch73-44r | uncharacte | 1 | 1.835302 | -0.3935 |
| A0A8M9Q9 | si:ch73-72k | netrin rece | 1 | 1.792518 | -0.54973 |
| A0A8M1PV | si:dkey-127 | serine/argi | 2 | 1.429695 | -0.47061 |
| A0A8M9PU | si:dkey-13a | uncharacte | 1 | 1.608624 | -0.6452 |
| A0A8M2BC | si:dkey-174 | uncharacte | 1 | 1.709923 | -0.7225 |
| Q5TZ78 | si:dkey-183 | Cytochrom | 6 | 2.937367 | -0.54126 |
| A0A8M2B6 | si:dkey-194 | piggyBac tr | 2 | 1.417912 | -0.60138 |
| A0A8M1RK | si:dkey-248 | uncharacte | 6 | 1.924605 | -0.70801 |
| A0A8M1RT | si:dkey-74k | uncharacte | 3 | 1.528238 | -0.38464 |
| A0A8M9PJ | si:dkey-79f | interferon- | 5 | 1.467761 | -1.01229 |
| A0A8M9P5 | si:dkey-9l2 | uncharacte | 1 | 2.202868 | -2.68604 |
| A0A8M3AN | si:dkeyp-50 | Calpain-1 c | 2 | 1.840266 | -0.56367 |
| A0A8M1RF | si:rp71-36a | uncharacte | 1 | 1.646516 | -0.48283 |
| A0A0R4IIC7 | si:zfos-1425 | IKAROS fan | 2 | 3.597259 | -0.80241 |
| Q7ZWG9 | sigmar1 | Sigma non- | 3 | 1.819664 | -0.43375 |
| A0A8M1N9 | slc12a3 | solute carri | 2 | 2.622279 | -0.5697 |
| A0A8M9QE | slc15a1a | solute carri | 4 | 1.807502 | -0.56328 |
| A1L1W9 | slc16a10 | Monocarbo | 1 | 3.802831 | -0.61252 |
| Q6P4V2 | slc1a3 | Amino acid | 3 | 1.374771 | -0.39615 |
| A0A8M9Q1 | slc25a23a | calcium-bir | 8 | 1.710431 | -0.46165 |
| A0A8M6Z9 | slc2a9l2 | solute carri | 1 | 1.710757 | -0.61301 |
| A0A8M1RH | slc44a1a | Choline tra | 7 | 1.780723 | -0.36886 |
| A0A8M2BD | slc44a2_1 | choline trai | 4 | 3.356017 | -0.59519 |
| A0A8M1PC | slit3 | slit homolo | 1 | 1.402701 | -0.36807 |
| A0A8M3AS | smarca2 | probable gl | 5 | 1.529363 | -0.35664 |
| B2ZFP3 | smarcal1 | SWI/SNF-re | 2 | 1.879801 | -1.43516 |
| A0A8N1TU | smc4 | Structural r | 8 | 1.716759 | -0.6493 |
| A0A0R4IZ8 | smg6 | Telomerase | 1 | 1.79829 | -0.32405 |
| Q7ZUG0 | snrpe | Small nucle | 1 | 1.480578 | -0.54085 |
| A0A8M1N7 | sord | Sorbitol de | 7 | 1.411421 | -0.60793 |
| A0A8M1NA | spint2 | uncharacte | 2 | 1.653435 | -0.46592 |
| A0A8M1P4 | stt3b | dolichyl-dip | 4 | 1.451501 | -0.40384 |
| A0A8M2B4 | tcea3 | transcriptic | 2 | 2.030921 | -0.50191 |
| A0A8M2BH | tcf3b | transcriptic | 2 | 1.922229 | -0.50345 |
| A0A8M2BD | tead3b | Transcriptio | 2 | 1.429976 | -0.37277 |
| A0A8M2B5 | tecpr2 | tectonin be | 1 | 1.435225 | -0.40737 |
| Q1L8P7 | thbs3b | Thrombosp | 6 | 1.576977 | -0.50674 |
| Q05AK9 | tma7 | Translation | 2 | 1.537206 | -0.47746 |
| A5PN43 | tmem175 | Endosomal | 1 | 1.352146 | -0.37263 |
| Q5CZV0 | tmem182a | Transmeml | 5 | 1.574752 | -0.34943 |
| A0JMD5 | tmprss4 | Transmeml | 1 | 1.415018 | -1.33256 |
| A0A8M2B4 | tmprss4a | transmemb | 4 | 1.799586 | -0.49235 |
| Q9I8U8 | tnnc | Fast skeleta | 32 | 2.856734 | -1.0296 |
| A0A8M2BA | tnni2a.4 | troponin I, | 19 | 1.714204 | -1.00591 |
| Q6DHP2 | tnni2b.2 | Troponin I | 12 | 2.551005 | -0.64167 |

| | | | | | |
|---|---|---|---|---|---|
| Q8AVB1 | tnnt1 | Slow tropo | 3 | 1.693654 | -1.38892 |
| A0A8M2BA | tnnt3a | troponin T | 12 | 3.824413 | -3.47148 |
| A0A8N7TD | tnpo1 | transportin | 3 | 1.580759 | -0.45536 |
| Q568B8 | tor4a | Torsin-4A | 1 | 1.878558 | -0.40527 |
| P13104 | tpma | Tropomyos | 138 | 2.361728 | -0.91271 |
| A0A8M9PC | trdn | Triadin | 21 | 1.847851 | -0.35806 |
| A0A8M9PT | trim67 | tripartite m | 1 | 2.858868 | -1.18189 |
| A0A8M1RK | tsg101b | tumor susc | 4 | 2.633987 | -1.20634 |
| A0A8M3AK | tspan9a | tetraspanir | 1 | 2.042302 | -0.49484 |
| A0A8M9Q( | ttn.1 | titin isoforr | 96 | 3.497563 | -0.86396 |
| A0A8M9Q | ttn.2 | titin | 114 | 3.858154 | -0.80457 |
| A0A8M1N8 | ugt1a1 | UDP-glucur | 2 | 2.121253 | -1.03206 |
| A0A8N1Z2I | ugt1ab | UDP glucur | 8 | 1.822732 | -0.81245 |
| A0A8M3B1 | unm_hu79 | aspartyl/as | 2 | 1.825271 | -0.49897 |
| A0A8M6YZ | upf2 | LOW QUAL | 1 | 1.425799 | -0.43641 |
| A0A8M9QL | usp2a | Ubiquitin c | 3 | 1.455354 | -0.4441 |
| F1QZW1 | vmhc | Myosin hea | 42 | 3.057036 | -0.39612 |
| A0A8M1NL | vsig8b | V-set and ir | 2 | 1.414253 | -0.49186 |
| Q1LWK5 | vwa9 | Integrator ( | 2 | 1.972926 | -0.5338 |
| F1QEB7 | wdr11 | WD repeat | 5 | 1.442766 | -0.4849 |
| A0A8M9Q( | wdr19 | WD repeat | 4 | 1.542807 | -0.36168 |
| A0A8N7UZ | wdr35 | WD repeat | 3 | 1.42953 | -0.56153 |
| A0A8N7TF | wfs1a | wolframin | 12 | 1.522458 | -0.46517 |
| Q7T389 | wu:fa03b0 | Phosphatid | 2 | 1.578445 | -0.41869 |
| Q7ZWB1 | wu:fa10c0 | Cyclin-depe | 2 | 1.491882 | -0.42983 |
| Q7T313 | wu:fa14b1 | Serine/thre | 2 | 1.606302 | -0.3479 |
| F1QBM2 | wu:fa55g0 | Programme | 9 | 1.763887 | -0.3224 |
| Q802U8 | wu:fb12d0 | H1 histone | 5 | 1.345485 | -0.44938 |
| A0A8M6Z1 | wu:fb16f0 | eukaryotic | 4 | 1.468718 | -0.50329 |
| Q9I9E6 | wu:fb33g0 | Trifunction | 10 | 2.065581 | -0.34353 |
| E7FAG0 | wu:fb77d1 | Vacuolar pr | 5 | 1.529111 | -0.37776 |
| Q08CV8 | wu:fc56h0 | Rps6ka1 pr | 1 | 1.311212 | -0.35946 |
| A6H8R5 | wu:fd12f1 | Zgc:16543( | 1 | 1.315507 | -0.49856 |
| Q7ZWC9 | wu:fd13g0 | 1-acyl-sn-g | 5 | 1.780539 | -0.6116 |
| A0A0R4IRS | wu:fe48d0 | Polynucleo | 5 | 1.842147 | -0.32972 |
| Q1LYI0 | wu:fi13b0 | UBX domai | 3 | 2.46641 | -0.41036 |
| A8CYR1 | wu:fi85h05 | Bromodom | 2 | 1.630083 | -0.3711 |
| Q7T067 | wu:fj61d12 | Rh blood gı | 4 | 1.316467 | -0.62843 |
| Q6PC10 | wu:fj66g02 | Branched-c | 1 | 1.727837 | -0.63901 |
| Q5BLF7 | wu:fl20e08 | Zgc:11308 | 17 | 2.227142 | -0.39871 |
| A0A8M1N/ | xpc | DNA repair | 3 | 2.228016 | -0.38885 |
| Q5SPJ8 | xpot | Exportin-T | 1 | 1.494434 | -0.43889 |
| Q5U3V6 | zgc:101592 | YTH domai | 1 | 2.135237 | -0.41105 |
| A0A8M9PK | zgc:101853 | uncharacte | 1 | 1.617014 | -0.41767 |
| Q5XJB5 | zgc:103457 | Zgc:103457 | 4 | 1.466595 | -0.35883 |
| Q5PR62 | zgc:103458 | Zgc:10345 | 3 | 1.590909 | -0.59446 |
| A0A8M2B8 | zgc:103482 | intracellula | 1 | 2.001113 | -0.59976 |

| | | | | | |
|---|---|---|---|---|---|
| Q5U3H4 | zgc:103524 | ribose-5-ph | 6 | 1.471283 | -0.49573 |
| Q5PR45 | zgc:103559 | allantoinas | 1 | 1.596084 | -1.27292 |
| Q800S8 | zgc:103687 | Fetuin-A | 3 | 2.656285 | -0.81827 |
| Q9I8U7 | zgc:109832 | Fast skeleta | 24 | 2.258038 | -1.1204 |
| A0A8N7UY | zgc:110216 | uncharacte | 1 | 1.655271 | -0.39998 |
| Q503H4 | zgc:110595 | Caspase 7, | 2 | 2.2268 | -0.63187 |
| Q567W1 | zgc:110715 | Troponin I, | 1 | 2.353458 | -1.82326 |
| A3KQN9 | zgc:111823 | Integrator ( | 2 | 2.101318 | -0.37359 |
| Q567D7 | zgc:112138 | Zgc:112138 | 13 | 1.723103 | -0.35414 |
| Q4V9I5 | zgc:112290 | Signal reco | 5 | 1.594428 | -0.40762 |
| Q4V8U2 | zgc:114143 | Secretory c | 3 | 1.523453 | -0.35152 |
| Q32PT2 | zgc:123217 | Zgc:123217 | 1 | 2.006543 | -0.79308 |
| Q32PS1 | zgc:123262 | Acyl-CoA th | 2 | 3.424503 | -0.68812 |
| Q08CC5 | zgc:153269 | Zgc:153269 | 2 | 1.678626 | -0.47824 |
| Q08C97 | zgc:153353 | Zgc:153353 | 3 | 1.782065 | -0.36935 |
| Q08C83 | zgc:153387 | Solute carr | 3 | 2.728212 | -0.33015 |
| A0JMG3 | zgc:153446 | Zgc:153446 | 1 | 1.333133 | -0.40117 |
| A0JMP1 | zgc:154079 | Endothelin- | 3 | 1.63157 | -0.36959 |
| Q1LUD0 | zgc:154080 | Interferon- | 6 | 2.017744 | -0.37167 |
| A2RUZ7 | zgc:158520 | Si:ch211-22 | 1 | 1.516797 | -0.51962 |
| A0A8M3BD | zgc:162816 | uncharacte | 2 | 1.340516 | -0.3422 |
| A3KP84 | zgc:163053 | D-beta-hyd | 2 | 1.459243 | -0.38259 |
| B3DHE5 | zgc:194224 | Zgc:194224 | 2 | 1.454517 | -0.38063 |
| Q803X9 | zgc:55364 | Voltage-de | 1 | 1.955037 | -1.07795 |
| Q802W1 | zgc:55670 | Chic2 prote | 1 | 1.519224 | -0.48065 |
| Q7ZUJ4 | zgc:56389 | Endoplasm | 6 | 1.882087 | -0.36688 |
| Q7ZUH8 | zgc:56518 | Microsoma | 1 | 2.419725 | -0.51165 |
| A0A8M6Z2 | zgc:63694 | high affinit | 2 | 1.372034 | -0.45424 |
| Q7T2B6 | zgc:64087 | Acyl-coenz | 5 | 1.555166 | -0.41815 |
| Q6P696 | zgc:66035 | Lactoylglut | 5 | 1.896196 | -0.32932 |
| Q6NWX4 | zgc:73160 | Serine/thre | 6 | 2.217202 | -0.33104 |
| Q6PEI4 | zgc:77437 | Tyrosine-pr | 1 | 1.349539 | -0.38582 |
| Q6P2T7 | zgc:77739 | UPF0462 p | 1 | 1.898675 | -0.37118 |
| Q6P2T6 | zgc:77740 | Cysteine ar | 1 | 1.708961 | -0.3477 |
| E7FG48 | zgc:77934 | Tubulin pol | 6 | 1.441077 | -0.40245 |
| Q6IQL9 | zgc:86709 | Novel actin | 68 | 2.248388 | -0.73135 |
| Q6IQE6 | zgc:86798 | small monc | 3 | 1.987927 | -0.35776 |
| Q6IQD5 | zgc:86813 | Ribokinase | 2 | 1.445843 | -0.63603 |
| F1RAK0 | zgc:92083 | Long-chain | 12 | 1.597732 | -0.43779 |
| A0A8M2B4 | zgc:92630 | dehydroger | 1 | 1.346074 | -0.47771 |
| Q6DGI1 | zgc:92912 | Zgc:92912 | 1 | 2.284654 | -0.65476 |
| Q6DGH3 | zgc:92922 | Protein tra | 3 | 1.879269 | -0.3408 |
| A0A8N7US | zmp:00000 | suppressor | 2 | 1.553566 | -0.52213 |
| A0A8M2B5 | zmp:00000 | nuclear fac | 1 | 2.538319 | -0.55783 |
